# Experimental change in personality: Overexpression of GDNF in the rat striatum converts the low exploratory phenotype into highly explorative

**DOI:** 10.1101/2025.03.16.643460

**Authors:** Jaanus Harro, Aet O’Leary, Marianna Norden, Kristi Liiver, Margus Kanarik, J. Arturo García-Horsman, Sven Sakson, Pilleriin Kupper, Karita Laugus, Sirin Korulu, Indrek Teino, Tõnis Org, Ilmari Parkkinen, Tanel Kaart, Päivi Lindholm, Sophie Imbeault, Helen Poska, Brandon K. Harvey, Mikko Airavaara, Ruth Shimmo, Mart Saarma

## Abstract

Major vulnerability factors for psychiatric disorders such as depression, that often prevent complete remission and lead to relapses, are temperamental. In a rat model of clustered persistent high anxiety/low motivation, we have found that overexpression of glial-cell-line-derived neurotrophic factor (GDNF) by intra-striatally administered adeno-associated virus vector strikingly converts the passive coping style of low exploratory rats into an active one, similar to high exploratory rats. This conversion of behavioural strategy developed gradually over repeated testing, and was associated with increased catecholamine metabolism in several brain regions and changes in the regulation of serotonin neurotransmission. An increase in *in vivo* dopamine transporter availability in the striatum was necessary for the phenotype conversion. Associated changes in striatal gene expression included key players in monoamine storage and epitranscriptomic regulation. The increase in GDNF signalling also caused alterations in levels and regional covariation of oxidative metabolism, indicative of persistent reorganization of neural activity throughout the brain. Thus, neurotrophic factors, GDNF in particular, may play a pivotal role in the development, persistence and alteration of personality traits, and therefore constitute a potential target for treatment of chronic, relapsing psychiatric disorders.

## Introduction

Mental health problems develop in a cascade where vulnerability embedded in temperament/personality traits will augment the negative impact of psychosocial and socioeconomic adversities [Gold et al. 1988; Wasserman et al. 2007; Harro and Kiive 2011; Rice et al. 2019; Kang et al. 2023]. Such a maladaptive development becomes sustained by the persistency of underlying neural networks. Understanding how such maladaptive neural networks are maintained, with the ultimate goal of providing tools to re-establish the suboptimal connectivity, is an important challenge for health sciences. A reductionistic approach to the factors that predict higher vulnerability to a variety of psychiatric disorders has identified the significant contribution of personality traits reflective of high negative emotionality, e.g., neuroticism, and low positive emotionality, e.g., extraversion or openness [Fava and Kendler 2000; Bienvenu and Stein 2003; Sanchez-Roige et al. 2011; Kang et al. 2023; Yang et al. 2024]. All this leads to the question of whether and how temperament/personality, defined as a characteristic behavioural pattern of reacting to environment stimuli, can be persistently altered. Recent studies on psychedelic-assisted therapy have suggested that a reduction in neuroticism and an increase in extraversion/openness are associated with therapeutic benefits [Holze et al. 2024; Pagni et al. 2025]. Understanding of the neurobiology that underlies a persistent shift in temperament would greatly benefit from the availability of an animal model.

One evolutionary behavioural strategy fundamental for survival is exploratory behaviour, which is represented in everyday search for vital resources, as well as in orientational response to any stimulus pattern that is novel to some degree [Berlyne 1966]. Exploratory behaviour in rodents is inter-individually largely variable [Rägo et al., 1988; Piazza et al. 1989], it is expressed as a persistent trait that is related to general health [Cavigelli and McClintock 2003], and inter-individual differences emerge even in genetically identical group-housed animals [Freund et al. 2013]. Lower exploratory behaviour predicts stress susceptibility [Milic et al. 2021]. Two of the core symptoms of depression, high anxiety and low motivation, cluster together in animals with persistently low exploratory activity [Mällo et al. 2007], as measured in the exploration box test [Harro et al. 1995; Otter et al. 1997]. In humans a constellation of high anxiety and low motivation would define the syndrome of melancholia, and if expressed trait-wise, represents the co-occurrence of high neuroticism and low extraversion. Such a personality type should be the strongest predictor of clinical depression [Kendler et al. 2006]. Rats with low exploratory activity in this test (LE-rats) exhibit high anxiety and anhedonia in a number of anxiety and depression-related tests, but these behavioural differences subside with handling and stress, while the defining phenotype itself is very resistant to any treatment and surviving at least to late adulthood [Mällo et al. 2007; Raudkivi et al. 2012]. LE-rats have lower extracellular dopamine levels in the striatum [Mällo et al. 2007, Alttoa et al. 2009], a smaller proportion of dopamine D_2_ receptors in high-affinity state [Alttoa et al. 2009], and a lower capacity of antidepressants to potentiate amphetamine-induced dopamine release [O’Leary et al. 2016]. In a mouse study, dopamine fluctuations in the dorsal striatum were shown to occur during spontaneous exploration, while despite genetic identity and common housing conditions, some animals were less sensitive to dopamine than others [Markowitz et al. 2023]. Low striatal dopamine function compromises effort-related behaviours and by this means underlies the emergence of depression [Salamone et al. 2016]. In remitted patients of recurrent depression, stressful stimuli selectively overactivate the striatum [Admon et al. 2015], suggesting that the striatum is a hub in the neurobiology of the traits on which vulnerability to mood and anxiety disorders is based.

Striatal dopaminergic neurotransmission depends, *inter alia*, on glial-cell-line-derived neurotrophic factor (GDNF). GDNF is a secreted growth factor that by binding to co-receptor GFRα1 activates several signalling pathways through the transmembrane tyrosine kinase receptor RET, regulating neuronal survival, neurite branching and synaptic plasticity [Airaksinen and Saarma 2002; Sidorova and Saarma 2020]. GDNF has been shown to support dopamine neuron survival and to stimulate dopamine release *in vitro* [Lin et al., 1993]. GDNF administration can increase dopaminergic activity in the mesocorticolimbic pathways [Salvatore et al. 2004; Wang et al. 2010], and alleviates symptoms of Parkinson’s disease [Gill et al., 2003; Kirik et al. 2017; Whone et al. 2019a; Whone et al. 2019b]. Endogenous GDNF-RET signalling plays an important role in the survival and function of dopamine neurons, and regulates dopamine transporter function and synaptic dopamine levels [Kramer et al. 2007; Pascual et al. 2008; Kumar et al. 2015; Kopra et al. 2017; Espinoza et al. 2019]. GDNF overexpression in the striatum protects and regenerates midbrain dopaminergic neurons, and stimulates dopamine release [Sidorova and Saarma 2020; Barker et al. 2024]. Dysregulated expression of GDNF and its co-receptor GFRα1 has been implicated in clinical depression [Maheu et al. 2015] and depression-like behaviours in animals [Zhang et al. 2019]. Given the dopaminergic deficit in the LE-rats and that endogenous GDNF enhances dopamine neuronal function and regulation of synaptic dopamine levels [Kopra et al. 2017], we considered the overexpression of GDNF a possible strategy of intervention. In this study, we first demonstrated that overexpression of GDNF induced with intra-striatal administration of adeno-associated virus (AAV1) vector reliably converts the passive coping strategy of LE-rats into an active one, similar to HE-rats, and subsequently identified the neurobiological mechanisms responsible for this change in a behavioural trait.

## Results

### GDNF overexpression elicits gradual conversion of a low exploratory to a high exploratory phenotype

We used a free exploration task, the exploration box test, (Fig. 1a) in which data distribution strongly deviates from normal (Fig. 1b). After screening in the exploration box test, rats were given viral vectors intrastriatally, repeatedly tested in the exploration box, thereafter submitted to *in vivo* experiments and subsequently sacrificed with brain tissues dissected for biochemical and histological studies (Fig. 1c). Treatment with AAV1-GDNF caused a gradual increase in exploration in the LE-rats over repeated testing, that reached the level typical of HE-rats (Fig. 1d). In order to exclude the possibility that the gradual increase in exploration in LE-rats after GDNF overexpression was simply related to time after administration of the vector, behavioural observations were started six weeks after administration of the viral vector in the first replication experiment, with essentially reproducing the finding in the previous experiment (Fig. 1e). This suggests that the profound increase in exploratory behaviour in LE-rats requires learning of a new behavioural strategy. We then used enzyme-linked immunosorbent assay (ELISA) to quantitatively measure the levels of the expression of GDNF. Viral vector mediated overexpression induced high and sustained levels of GDNF in the striatum (31.5±3.6 ng/mg tissue in the control group, AAV1-GFP (green fluorescent protein) treated rats vs. 3668.9±449.1 ng/mg in AAV1-GDNF treated groups) that were above the level that is required for the near-complete neuroprotection as previously described in the 6-hydroxydopamine lesion model [Georgievska et al. 2002]. We also detected an increase of GDNF also in the nucleus accumbens (45.0±15.4 ng/mg tissue in AAV1-GFP treated rats vs. 2180.1±616.4 ng/mg in AAV1-GDNF treated groups; Fig. 1f). Control HE-rats had higher levels of GDNF (49.1±0.9 ng/mg tissue) than LE-rats (25.6±2.5 ng/mg tissue) in the striatum but not in the accumbens (Fig. 1g). We then performed tyrosine hydroxylase (TH) immunostaining on striatal sections. Intensities of TH staining were higher in rats with the active approach to novelty, that is, in HE-rats and in LE-rats after AAV1-GDNF administration after excluding the few non-responders to this treatment (Fig. 1h). This suggests that RET-dependent signalling is important for exploratory behaviour.

**Fig. 1:**
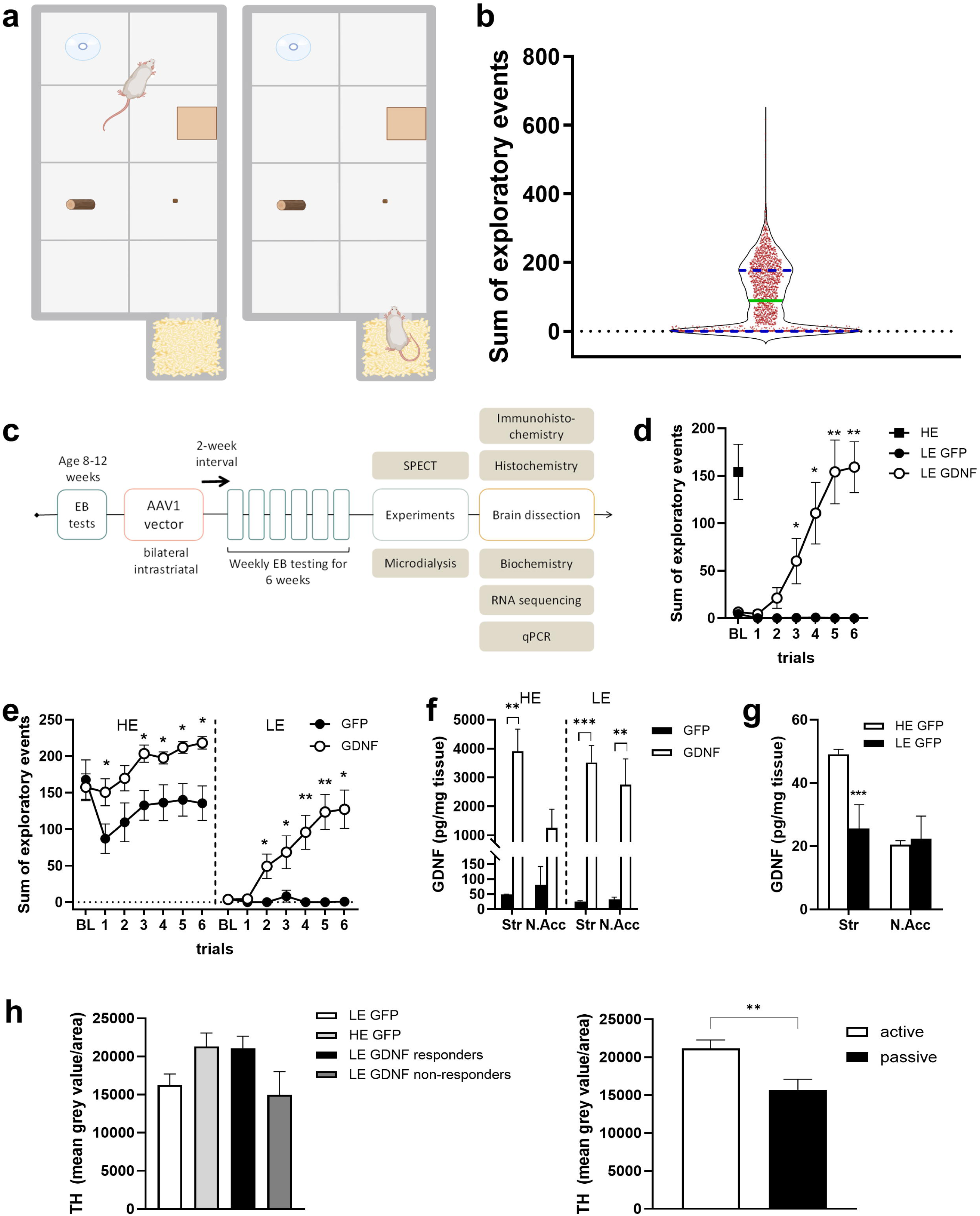
Exploratory behaviour of LE-rats increases dramatically after overexpression of GDNF. (a) A scheme of the exploration box test. (b) Exploratory activity in 2159 male rats, assessed in 2004-2024 in four different animal facilities, does not follow normal distribution (Shapiro-Wilks W=0,88; p < 0.0001). Green horizontal line - median, blue horizontal lines - quartiles; 27% of the animals did not enter the open area. (c) Experimental design of studies on the effect of GDNF overexpression in LE- and HE-rats. EB, exploration box; SPECT, single photon emission computerized tomography. (d) Exploratory behaviour of LE-rats before and starting from two weeks after treatment with AAV1 (repeated-measures ANOVA, LE-GFP vs. LE-GDNF, GDNF effect: F(1,14)=15.74; p=0.0014; LE-GFP n=7; LE-GDNF n=9; HE-GFP n=7). (e) Exploratory behaviour of rats treated with AAV1-GFP or AAV1-GDNF six or eight weeks before beginning of post-treatment observations (two-way ANOVA, Day × HE/LE × GDNF interaction: F(6,221)=2.863; p=0.011; LE-GFP n=11; LE-GDNF n=14; HE-GFP n=7; HE-GDNF n=9). (f) GDNF levels by ELISA in corpus striatum and nucleus accumbens after treatment with AAV1-GFP or AAV1-GDNF (two-way ANOVA, GDNF effect in striatum: F(1,21)=47.6; p<0.0001; GDNF effect in nucleus accumbens: F(1,21)=7.923; p=0.010; LE-GFP n=9; LE-GDNF n=8; HE-GFP n=3; HE-GDNF n=5). (g) Control LE-rats had lower levels of GDNF in striatum but not in accumbens (unpaired t-test, striatum: t=5.214, df=10; p=0.0004; nucleus accumbens: t=0.3542, df=7; p=0.734; LE-GFP n=9; HE-GFP n=3) (h) Tyrosine hydroxylase immunostaining in striatum was higher in actively exploring rats (one-way ANOVA, F(3,13)=2.808; p=0.081; LE-GFP n=4; HE-GFP n=4; LE-GDNF responders n=6; LE-GDNF non-responders n=3; active (n=10) vs. passive (n=7) groups, unpaired t-test, t=3.058; p=0.008). Data are presented as the mean ± s.e.m.; see Source Data for statistical details. *P < 0.05,**P < 0.01, ***P < 0.001 compared to respective control.

### GDNF overexpression affects dopamine and serotonin metabolism and dopamine overflow while behavioural conversion requires an increase in dopamine transporter availability

*Ex vivo* measurements revealed the effect of GDNF overexpression on monoamine neurotransmission in the striatum. Thus, dopamine turnover was increased as indicated by increased ratio of 3,4-dihydroxyphenylacetic acid (DOPAC) and DA in GDNF overexpressing rats (Fig. 2a), but also were increased the levels of 5-hydroxyindoleacetic acid (5-HIAA), the main metabolite of serotonin (Fig. 2b). Furthermore, monoamine metabolism was also affected in other brain regions. In the nucleus accumbens, GDNF overexpression increased the levels of metabolites of dopamine and serotonin, somewhat more prominently in LE-rats (Fig. 2c). The trend of an increase in dopamine and DOPAC after GDNF overexpression was also observed in the hippocampus (Fig. 2d), and in LE-rats, AAV1-GDNF treatment additionally increased 5-HIAA levels in the frontal cortex (Fig. 2e). In the hypothalamus, striatal GDNF overexpression caused higher levels of normetanephrine, indicative of increased noradrenergic activity (Fig. 2f), while in the raphe area, intrastriatal administration of AAV1-GDNF significantly decreased 5-HIAA levels and increased DOPAC/DA ratio, but only in HE-rats (Fig. 2g).

**Fig. 2:**
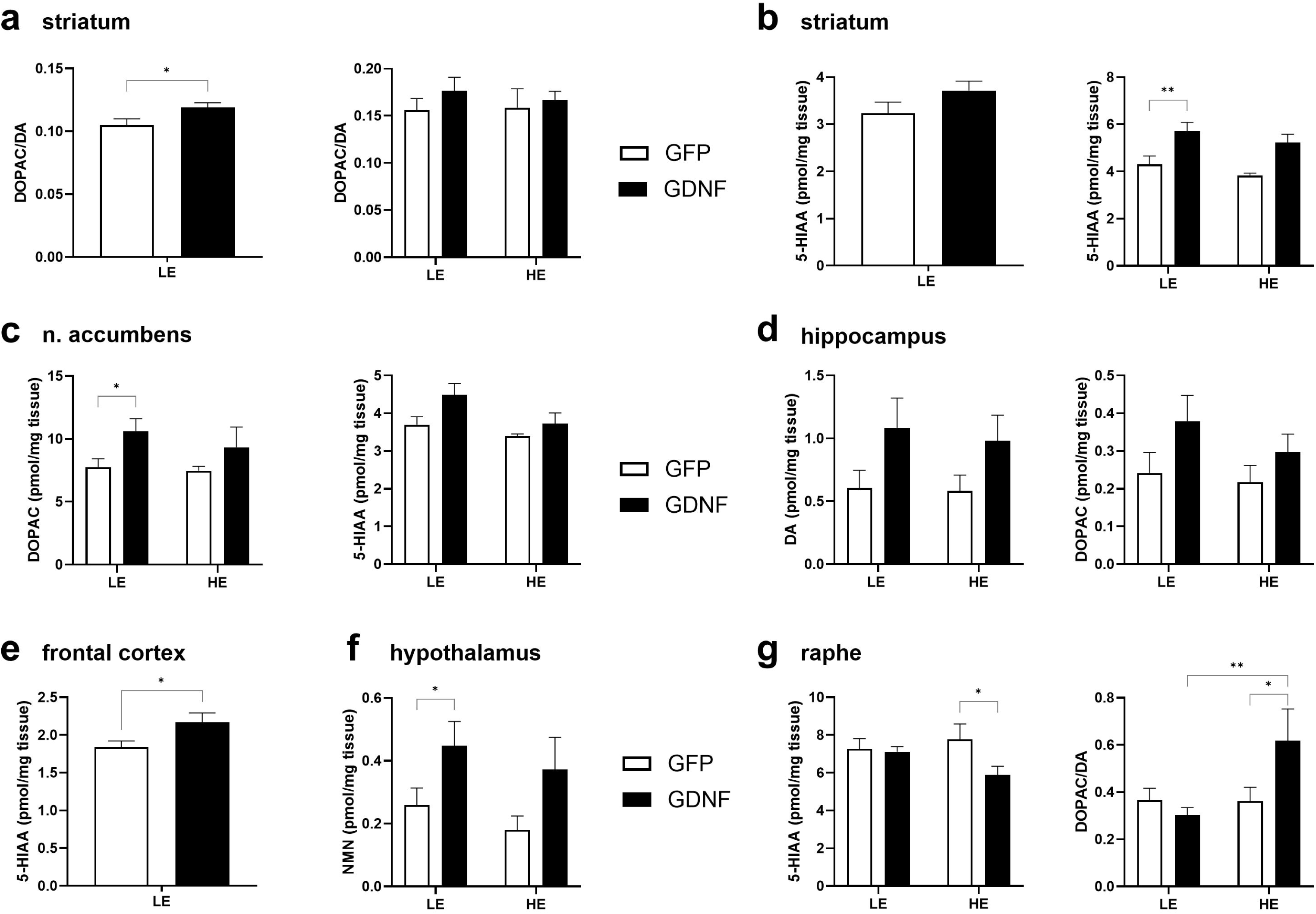
GDNF overexpression affects dopamin- and serotonergic neurotransmission in striatum and throughout the brain. (a) GDNF overexpression increased dopamine turnover in striatum, expressed as DOPAC/DA ratio. This was statistically significant in the first experiment (left; Student’s t-test, t=2.22, df=13; p=0.045; LE-GFP n=6; LE-GDNF n=9) and observed as a tendency when animals were sacrificed after microdialysis (right; LE-GFP n=13; LE-GDNF n=11; HE-GFP n=7; HE-GDNF n=9). (b) GDNF overexpression increased serotonin turnover in striatum, expressed as 5-HIAA/5-HT ratio. This was observed as a tendency in the first experiment (left; LE-GFP n=6; LE-GDNF n=9) and statistically significant when animals were sacrificed after microdialysis (right; two-way ANOVA, GDNF effect: F(1,36)=14.78; p=0.0005; LE-GFP n=13; LE-GDNF n=11; HE-GFP n=7; HE-GDNF n=9). (c) Striatal GDNF overexpression increased the levels of DOPAC (left; two-way ANOVA, GDNF effect: F(1,54)=4.27; p=0.043) and tended to increase 5-HIAA (right; two-way ANOVA, GDNF effect: F(1,54)=3.35; p=0.073) in the nucleus accumbens. (d) Striatal GDNF overexpression tended to increase the levels of dopamine (left; two-way ANOVA, GDNF effect: F(1,70)=4.98; p=0.029) and DOPAC (right; two-way ANOVA, GDNF effect: F(1,25)=3.45; p=0.075) in hippocampus. (e) Increase of 5-HIAA levels in LE-rats in frontal cortex after GDNF overexpression in striatum (Student’s t-test, t=2.16, df=40; p=0.036). (f) Increase of normetanephrine levels in the hypothalamus after GDNF overexpression in striatum (two-way ANOVA, GDNF effect: F(1,39)=6.31; p=0.016). (g) Striatal GDNF overexpression decreased the levels of 5-HIAA (left; two-way ANOVA, GDNF effect: F(1,61)=4.06; p=0.048) and increased DOPAC/DA ratio (right; two-way ANOVA, HE/LE × GDNF interaction: F(1,61)=5.27; p=0.025) in the raphe area in HE-rats. Data are presented as the mean ± s.e.m.; see Source Data for statistical details. *P < 0.05; **P < 0.01 compared to respective comparison group.

Previous studies, similarly to the experiments presented herewith, have not revealed any difference between LE-and HE-rats in the tissue levels of dopamine and intrastriatal administration of a low dose of 6-OHDA to HE-rats reduced their high level of exploration only modestly (Suppl. Fig. 1). However, higher levels of extracellular levels of dopamine in have consistently been detected in the striatum of HE-rats across multiple conditions [Mällo et al. 2007; Alttoa et al. 2009; O’Leary et al. 2016]. *In vivo* microdialysis in the striatum revealed a reduction in extracellular levels of dopamine after GDNF overexpression in HE-rats who naturally have higher levels of extracellular dopamine than LE-rats. This was particularly evident in release-stimulating conditions such as locally induced depolarization (Fig. 3a), or administration of amphetamine (Fig. 3b). However, if LE-rats were stratified according to their behavioural response to AAV1-GDNF, it was revealed that in LE GDNF-non-responders the reduction in the striatal extracellular dopamine levels by GDNF overexpression was similar to the AAV1-GDNF-treated HE-rats, but the small number of non-responders had masked the overall effect in LE-rats (Fig. 3c). Hence, in LE-rats that became highly exploratory after GDNF overexpression, the synaptic regulation of dopamine had been changed. GDNF non-responders also had lower 5-HT levels in the striatum (Fig. 3d).

**Fig. 3:**
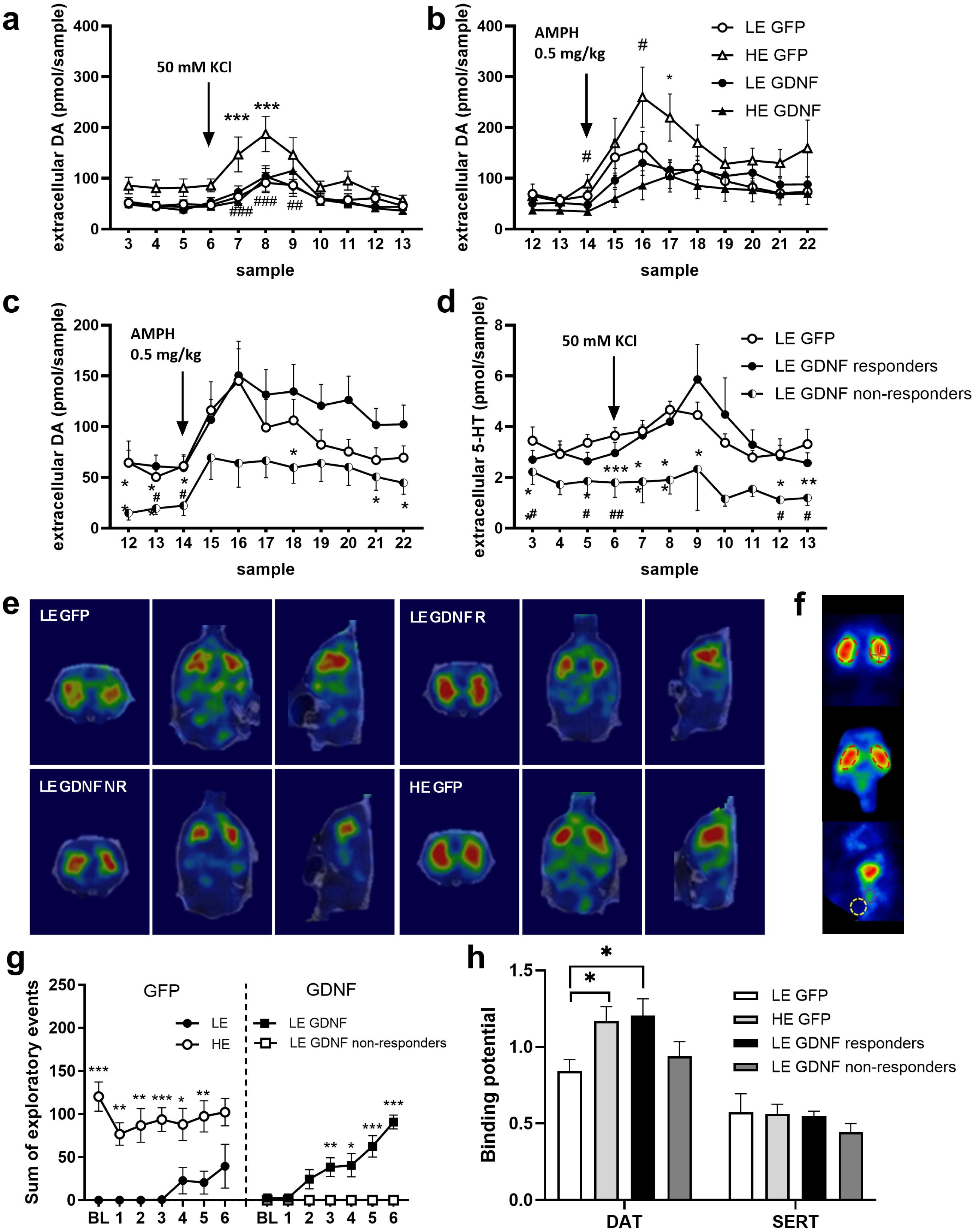
Behavioural trait conversion by GDNF overexpression requires alterations in synaptic regulation of dopamine. (a) Dopamine overflow in striatum was reduced by GDNF overexpression in HE-rats (LE-GFP n=19; LE-GDNF n=20; HE-GFP n=10; HE-GDNF n=11; two-way ANOVA with repeated measures, Time × HE/LE × GDNF interaction: F(10,560)=1.92; p=0.041;***P < 0.001 compared to LE-GFP rats; ^##^P < 0.01; ^###^P < 0.001 HE-GDNF compared to HE-GFP rats). (b) Effect of D-amphetamine (0.5 mg/kg intraperitoneally) on striatal dopamine release was also smaller after GDNF overexpression in HE-rats (LE-GFP n=10; LE-GDNF n=15; HE-GFP n=7; HE-GDNF n=8; repeated-measures two-way ANOVA, Time × HE/LE × GDNF interaction: F(10,360)=2.16; p=0.019; *p < 0.05 compared to LE-GFP rats; ^#^p < 0.05 HE-GDNF compared to HE-GFP rats). (c) In GDNF non-responder LE-rats, striatal extracellular dopamine levels were lower (LE-GFP n=10; LE-GDNF non-responders n=3; LE-GDNF responders n=12; repeated measures ANOVA, Time effect: F(4,037, 76,71)=7.87; p < 0.0001; *P < 0.05; **P < 0.01; ***P < 0.001 comparison of LE-GDNF responders and non-responders; ^#^P < 0.05; ^##^P<0.01 LE-GDNF non-responders compared to LE-GFP rats). (d) Extracellular levels of serotonin in striatum were also lower in GDNF non-responder LE-rats (LE-GFP n=10; LE-GDNF non-responders n=2; LE-GDNF responders n=8; mixed-effects model: Time effect: F (2,394, 54,59) = 3.872; p=0.021; *P < 0.05; **P < 0.01 comparison of LE-GDNF responders and non-responders; ^#^P < 0.05 LE-GDNF non-responders compared to LE-GFP rats). (e) SPECT imaging with ^123^I-FP-CIT: coronal, axial and sagittal views of representative rats of each group. (f) SPECT imaging with ^123^I-FP-CIT showing volumes of interest marked with dashed line, striatum (red) for DAT, midbrain (magenta) for SERT, and cerebellum as background (yellow). (g) Effect of GDNF overexpression on exploration box behaviour in the SPECT imaging experiment (repeated-measures ANOVA, Group effect: F(1,12)=16.42; p=0.0016; Day × Group interaction: F(6,72)=5.391; p < 0.0001; LE-GFP n= 7; HE-GFP n=7; LE-GDNF responders n=10; LE-GDNF non-responders n=4. (h) DAT binding potential (BP) in striatum was lower in LE-rats as compared to HE-rats; in LE-rats responding to GDNF overexpression, DAT BP was at the level of HE-rats, while this was not the case in non-responders (one-way ANOVA, Group effect: F(3,24)=3.04; p=0.048; *P < 0.05;**P < 0.01; ***P < 0.001 compared to LE-GFP rats; ^#^P < 0.05; ^##^P<0.01; ^###^P < 0.001 compared to LE-GDNF non-responders. Ligand binding to 5-HT transporters in the brainstem was not different between groups.

As a further experiment (Fig. 3g), we conducted SPECT imaging with ^123^I-FP-CIT that labels both dopamine and serotonin transporters, the former quantified by tracer binding in the striatum and the latter in the mid-brain (Fig. 3ef). The proportion of LE- and HE-rats varies by batches of animals substantially, and for this experiment, the number of HE-rats was sufficient only to form the control group. LE-rats had lower DAT availability than HE-rats, and GDNF overexpression increased DAT availability of LE-rats to the level of HE-rats (Fig. 3h). Importantly, in the four LE-rats that in this experiment failed to respond to GDNF overexpression, DAT availability was not significantly increased. Serotonin transporter availability in the midbrain was neither significantly different between LE- and HE-rats nor affected by GDNF overexpression.

### While the striatal transcriptomic landscape is divergent in LE-rats, GDNF overexpression has multiple effects on striatal gene expression that correspond to the changed behavioural trait

In order to find novel clues how GDNF overexpression in the striatum can convert the exploratory phenotype, we performed mRNA sequencing in rats from one further experiment in which the LE-phenotype was converted to HE-like by AAV1-GDNF treatment (Fig. 4a). On the basis of statistical significance, GDNF overexpression affected a larger number of genes in HE- as compared to LE-rats (at nominal p<0.05 level, n=869 vs 299, respectively; at p<0.001 level, n=53 vs. 9, respectively; Suppl. Table 1). A similar outcome was observed in GO pathway analysis: the number of pathways affected by GDNF overexpression (at nominal p<0.05 level) was 118 vs. 7 (Biological Processes), 62 vs. 6 (Cellular Component), 34 vs. 5 (Molecular Function) and 18 vs. 7 (KEGG pathway) for HE- and LE-rats, respectively. This was, however, related to the fact that the gene expression landscape of the LE-phenotype is significantly more diverse than that of the HE-phenotype (Fig. 4b). Similarly, KEGG pathway analysis revealed significant enrichment of neurotransmission-related pathways after GDNF overexpression in the HE-rats; in the LE-rats, only a modest number of signalling pathways were enriched after GDNF overexpression. Of note, the oxytocin signalling pathway and the calcium signalling pathway were enriched in both LE- and HE-rats (Fig. 4c). We then used the STRING database (https://string-db.org/) to explore protein-protein interactions in the sets of differentially expressed genes, and again found significantly enriched protein interactions only in the HE-GFP vs. HE-GDNF comparison. While the differentially expressed genes between the LE-GFP and LE-GDNF did not yield a significantly enriched STRING network, we took notice of a gene of interest (*Mettl3*; encoding methyltransferase 3, N6-adenosine-methyltransferase complex catalytic subunit) with its interaction partners (see below) (Suppl. Fig. 2).

**Fig. 4:**
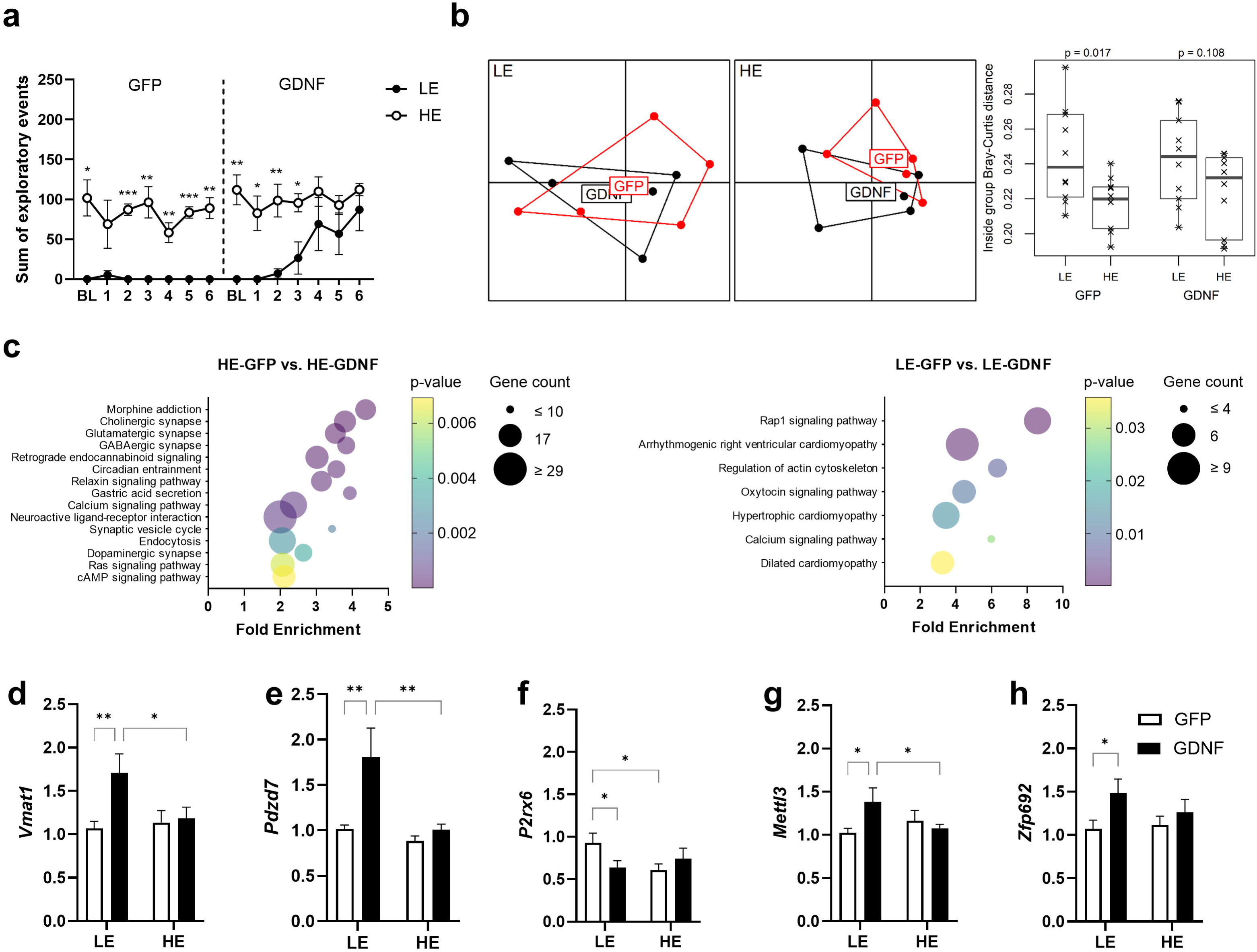
GDNF overexpression elicited changes in striatal gene expression landscape. (a) Behavioural effect of intrastriatal AAV1-GDNF treatment in the mRNA sequencing experiment (in each group n=5; two-way ANOVA with repeated measures: HE/LE effect: (1,16)=50.48; p < 0.0001; GDNF effect: F(1,16)=6.11; p=0.025; post-hoc *P < 0.05;**P < 0.01; ***P < 0.001 compared to the corresponding LE-group). (b) Principal component analysis of standardized expression of 21,657 genes that had data from all rats (PC1: 24.7%, PC2: 10.0%). Dots mark the location of samples according to their first two principal component scores sorted by exploring behaviour (LE and HE) and treatment (GFP and GDNF). Samples from the same group are surrounded by a line; treatment names in boxes mark the group centroids. Inside treatment (GFP and GDNF) and exploring behaviour (LE and HE) group Bray-Curtis distances calculated based on standardised expression data of 21,657 genes. Crosses mark single pairwise inside group distances and p-values above the figure indicate the statistical significance of LE- and HE-group difference inside treatment group (t-test). (c) Enriched KEGG pathways in the HE-GFP vs. HE-GDNF groups (left) and LE-GFP vs. LE-GDNF groups (right). (d) Independent replication of the increased expression of *Vmat1* by GDNF overexpression in LE-rats (two-way ANOVA, GDNF effect: F(1,88)=4.24; p=0.043; LE-GFP n=21; LE-GDNF n=29; HE-GFP n=21; HE-GDNF n=21). (e) Independent replication of the increased expression of *Pdzd7* by GDNF overexpression in LE-rats (two-way ANOVA, GDNF effect: F(1,77)=4.86; p=0.031; LE-GFP n=16; LE-GDNF n=25; HE-GFP n=19; HE-GDNF n=21). Independent confirmation that *P2rx6* is higher in LE-rats and was reduced by GDNF overexpression (two-way ANOVA, HELE × GDNF interaction: F(1,84)=4.43; p=0.038; LE-GFP n=22; LE-GDNF n=25; HE-GFP n=21; HE-GDNF n=21). (g) Independent replication of the increased expression of *Mettl3* by GDNF overexpression in LE-rats (two-way ANOVA, HE/LE x GDNF interaction: F(1,75)=4.02; p=0.049; LE-GFP n=17; LE-GDNF n=21; HE-GFP n=20; HE-GDNF n=21). (h) Independent replication of the increased expression of *Zfp692* by GDNF overexpression in LE-rats (two-way ANOVA, GDNF effect: F(1,83)=4.24; p=0.042; LE-GFP n=21; LE-GDNF n=25; HE-GFP n=21; HE-GDNF n=21). *P < 0.05;**P < 0.01, compared to respective comparison group.

Some genes highlighted in the RNA sequencing experiment and STRING analysis were further examined by RT-qPCR in striatal samples collected in several independent experiments. In the striatal mRNA sequencing experiment, GDNF overexpression in the LE-rats had the statistically most significant effect on *Vmat1*, vesicular monoamine transporter 1; an independent experiment revealed a similar increase in LE-rats (Fig. 4d). Expression of the second gene in the list ranked by statistical significance, *Pdzd7* (PDZ domain containing 7), was also statistically significantly increased by AAV1-GDNF treatment in other experiments (Fig. 4e). Another gene of interest, tested independently after a statistically significant effect of GDNF overexpression was found in mRNA sequencing, was the ATP receptor *P2rx6*: This gene was more highly expressed in LE-rats, and this was brought down to the HE-rat level by GDNF overexpression (Fig. 4f). Of other genes with nominally significantly altered expression, our interest was specifically evoked by the effect of GDNF overexpression on *Mettl3*, an RNA m^6^A methyl transferase gene [Meyer and Jaffrey 2017], because we had recently revealed a simultaneous behaviourally activating and anxiolytic-like effect of *Mettl3* activation [Kanarik et al. 2025], by use of the first-in-class METTL3-METTL14-WTAP complex activating compound [Selberg et al. 2019]. GDNF overexpression elicited an increase in the expression of *Mettl3* exclusively in LE-rats (Fig. 4g). In the STRING analysis, two genes that were significantly affected by GDNF overexpression, *Zfp692* (zinc finger protein 692) and *Lrpprc*, were associated with *Mettl3*. We examined the expression of these two genes in independent experiments and confirmed the upregulation of *Zfp692* (Fig. 4h), but not *Lrpprc*.

### Conversion of the explorative phenotype after GDNF overexpression occurs together with changes in oxidative metabolism and the covariation matrix of regional cerebral activity

The energy requirement of the nervous system is reflected in the production of ATP, that is dependent on the activity of cytochrome c oxidase (COX; EC 1.9.3.1; also complex IV) in the mitochondrial electron transport chain. COX activity is relatively stable in time and hence can serve as a reliable indicator for neural basis of behavioural traits [Wong-Riley et al. 1998]. COX histochemical mapping provides a snapshot of mainly excitatory neurotransmission of the entire brain. This method has revealed brain regions and patterns of regional co-variance that are shared across a number of mouse [Matrov et al. 2019] and rat [Harro et al. 2014] models of depression and psychiatric vulnerability. As with the SPECT experiment, we did not have a sufficient number of HE-rats for two treatment conditions, so AAV1-GDNF was administered only to LE-rats (Fig. 5a). COX histochemistry revealed several group differences in subcortical areas along the rostrocaudal axis (Fig. 5b; Suppl. Table 1).

**Fig. 5:**
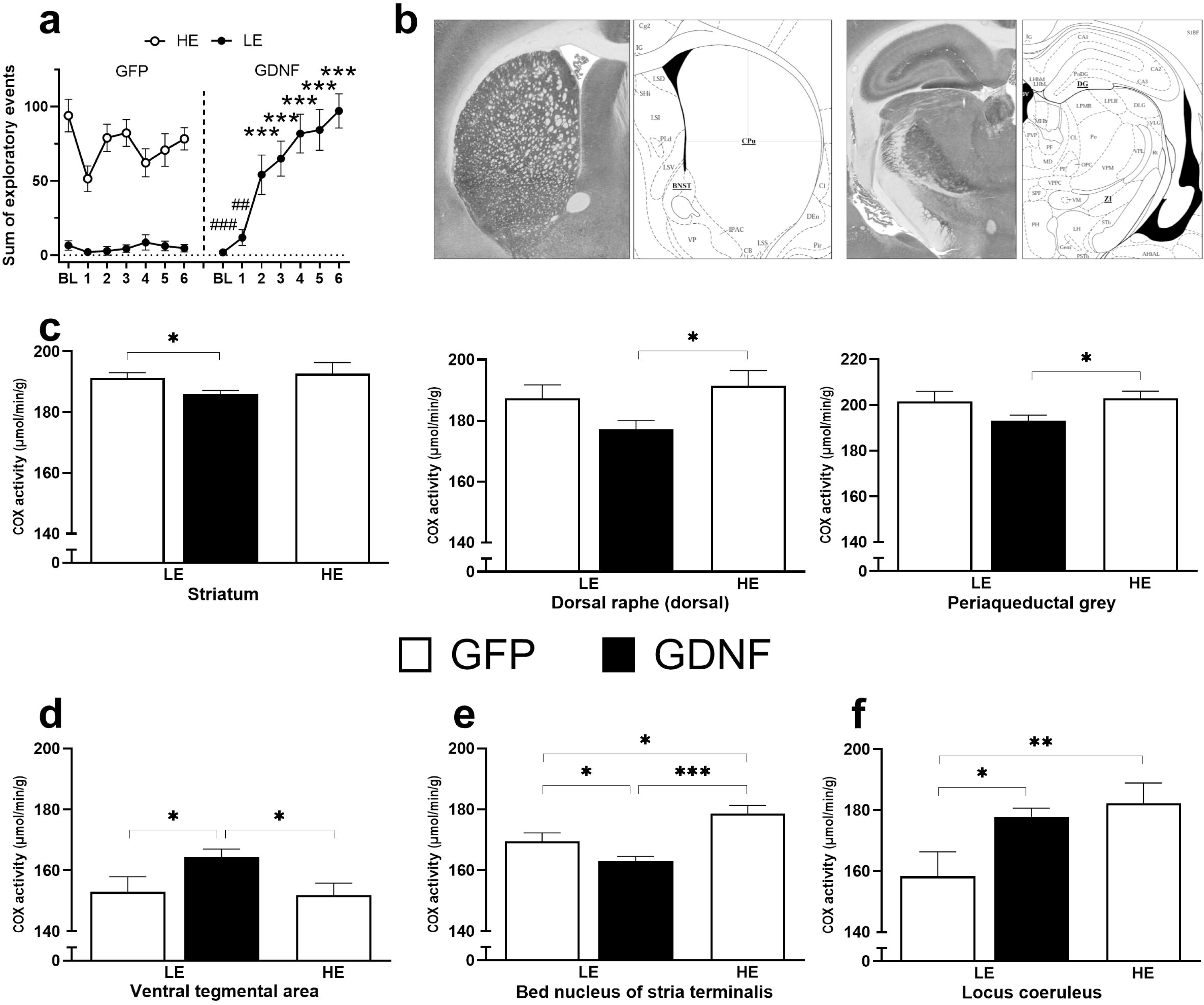
Conversion of the exploratory phenotype after GDNF overexpression occurs together with changes in cytochrome oxidase histochemistry indicative of oxidative metabolism in several brain regions, and in the regional covariation matrix of cerebral activity. (a) Behavioural effect of intrastriatal AAV1-GDNF treatment in the oxidative metabolism mapping experiment (LE-GFP n=10; LE-GDNF n=17; HE-GFP n=11; ANOVA with repeated measures, Group effect: F(2,35)=22.08; p < 0.0001; Time effect: F(6,210)=8.52; p < 0.0001; Time × Group interaction: F(12,210)=9.98; p<0.0001; post-hoc comparisons: ***P < 0.001 compared to LE-GFP group; ^##^P < 0.01, ^###^P < 0.001 compared to HE-GFP group). (b) Representative histochemical images of COX activity at the level of striatum (left) and hippocampus and zona incerta (right). (c) GDNF overexpression reduced oxidative metabolism in striatum (F(2,35)=2.76; p=0.077; comparison of LE-GFP and LE-GDNF, F(1,25)=5.63; p=0.026), dorsal raphe (F(2,35)=3.74; p < 0.05) and lateral periaqueductal grey matter (F(2,35)=3.41; p=0.044). (d) GDNF overexpression increased oxidative metabolism in VTA (F(2,35)=3.97; p < 0.05). (e) Lower oxidative metabolism of the LE-rats in the bed nucleus of stria terminalis was further reduced by GDNF overexpression (F(2,35)=12.90; p < 0.0001). (f) In locus coeruleus, GDNF overexpression increased oxidative metabolism of LE-rats to the level of HE-rats (F(2,35)=4.61; p < 0.05; *P < 0.05;**P < 0.01; ***P < 0.001 compared to the corresponding group). (g) A scheme of changes in regional activity covariation after GDNF overexpression (Created in Biorender.com). dmSTR, dorsomedial striatum; CL, claustrum; DG, dentate gyrus; ZI, zona incerta; MeA, medial nucleus of amygdala; CoA, cortical nucleus of amygdala; vlPAG, ventrolateral periaqueductal grey matter; DRD, dorsal part of dorsal raphe; LC, locus coeruleus. *P < 0.05 compared to LE-GFP; Pearson’s correlations compared with the z-test for r-to-z transformed coefficients.

LE- and HE-rats had similar levels of oxidative metabolism in the striatum, dorsal part of the dorsal raphe, and lateral periaqueductal grey matter, while GDNF overexpression reduced it (Fig. 5c). In contrast, striatal GDNF overexpression increased oxidative metabolism in the ventral tegmental area (VTA) (Fig. 5d). LE-rats had lower oxidative metabolism throughout the bed nucleus of stria terminalis (BNST), and this was further reduced by GDNF overexpression (Fig. 5e). In the locus coeruleus, LE-rats had lower levels of oxidative metabolism, and GDNF overexpression increased it to the level of HE-rats (Fig. 5f). Thus, GDNF overexpression had a complex effect on regional oxidative metabolism. Hence, we examined the correlation matrix of regional activities in AAV1-GFP treated HE- and LE-rats and in LE-rats with exploratory activity increased by GDNF overexpression (Suppl. Table 2). There were several potentially interesting differences between LE- and HE-rats and a variety of effects of GDNF overexpression, but hereby we have focused on those cases where the correlation between two regions was most clearly altered by GDNF overexpression, making it similar to the HE-rats. Thus, in control LE-rats, oxidative metabolism in the claustrum was in a strong positive correlation with that in the medial and cortical amygdala; these correlations were weakly negative in HE-rats and in LE-GDNF rats (Fig. 5g). Similarly, the strong positive correlation of the dorsomedial part of striatum with the dentate gyrus in LE was absent or weakly negative in HE-rats as well as in the LE-GDNF group; similar was its activity co-variation with zona incerta. In contrast, a strong negative correlation between COX activities of the locus coeruleus and zona incerta was present in control LE-rats, whereas the correlation was moderately positive in HE-rats, and in LE-rats after GDNF overexpression. In control LE-rats only a weak positive correlation was found between oxidative metabolism in the ventrolateral PAG and the dorsal part of dorsal raphe. However, this correlation was strongly positive in HE-rats, and in LE-rats after GDNF overexpression. Of the brain regions where striatal GDNF overexpression affected oxidative metabolism, the BNST had a distinct profile: First, control LE-rats had significantly lower oxidative metabolism than HE-rats, and AAV1-GDNF treatment reduced this further; second, both control LE- and HE-rats had a negligible correlation between activity in BNST and in the nucleus accumbens and claustrum, while administration of AAV1-GDNF resulted in strong positive correlations between BNST subregions and the accumbens, and with the claustrum (Suppl. Table 2).

## Discussion

We report that bilateral intrastriatal overexpression of GDNF elicits a gradual increase in exploratory behaviour in rats with persistent low-exploring (LE) phenotype. This behavioural conversion can serve as a model for further studies on experimental alteration of persistent behavioural traits, with implications to future therapies for chronic and relapsing disorders.

In previous studies, a variety of neurobiological differences between LE- and HE-rats have been found, that are consistent with the LE-rats being highly anxious and low in motivation [Alttoa et al. 2007; Alttoa et al. 2010; O’Leary et al. 2016]. Across experiments, the hallmark of the LE-phenotype has been a lower dopaminergic capacity in the striatum [Mällo et al. 2007; Alttoa et al. 2009; O’Leary et al. 2016]. Because GDNF expression in the rodent striatum in the postnatal period is necessary for the survival and functioning of midbrain dopaminergic neurons [Granholm et al. 2000; Sidorova and Saarma 2020], an isoform-specific decrease of the GDNF co-receptor GFRα1 has been found in depression [Maheu et al. 2015], in animal models, depression-like behaviours were associated with higher GDNF gene promoter methylation and lower GDNF mRNA levels [Zhang et al. 2019], and endogenous GDNF enhances dopamine transporter function and regulation of synaptic dopamine levels [Kopra et al. 2017], we considered the overexpression of GDNF as a possible strategy for intervention to rectify the dopaminergic deficit in the LE-rats.

In these experiments, most of the LE-rats did not leave their home base at screening. Only a few rats given AAV1-GFP had some increase in exploratory activity, possibly owing to unspecific effect of intracerebral manipulation or the effect of general anaesthesia. Indeed, a single-dose ketamine treatment can have long-lasting antidepressant effects in chronic, treatment-resistant depression [Krystal et al. 2013; Shiroma et al. 2020], and other anaesthetics have also shown antidepressant properties [Kohtala and Rantamäki 2021]. But in contrast to the infrequent behavioural change with the control AAV1-GFP treatment, after AAV1-GDNF administration the vast majority of LE-rats started to emerge from the small compartment of the exploration box and explore the large open area, gradually achieving the activity level of HE-rats. The fact that this phenotypic alteration developed in the course of repeated testing, suggests the involvement of learning a new behavioural strategy, which however first required the reduction of anxiety and an increase in exploratory drive caused by overexpression of GDNF in the striatum. Of note, control LE-rats had lower levels of GDNF in the striatum than HE-rats, but no differences were present between the groups in the nucleus accumbens. This suggests a functional specificity, that resembles the findings on extracellular dopamine levels that are lower in LE-rats in the striatum but not in the accumbens [Mällo et al. 2007]. On the other hand, overexpression of GDNF was also found after AAV1-GDNF in the few LE-rats with no increase in exploratory activity. Thus, further downstream effects are required for the behavioural conversion.

Previously, one study has attempted a similar approach, by increasing the expression of neurotrophin-3 in the dorsal amygdala and observing a degree of improvement in the anxious temperament of rhesus monkeys [Fox et al. 2019]. An increase of GDNF expression by viral vector delivery has been used in Parkinson’s disease patients with variable but so far modest effects on the main symptoms, possibly due to the advanced neurodegeneration present in trial subjects [Airavaara and Saarma 2024; Barker et al. 2024]. That expression of GDNF is relevant to mood and anxiety was highlighted in a recent study, where administration of GDNF into the medial prefrontal cortex mitigated the depression-like phenotype of mice in the MPTP model of Parkinson’s disease via local enhancement of dopamine neurotransmission [Liu et al. 2024]. Also, Ford and co-authors [2023] have demonstrated, in rhesus monkeys, that GDNF overexpression in the VTA resulted in prolonged alcohol abstinence and resistance to reintroduction challenges.

Overexpression of GDNF in the striatum elicited changes in monoamine systems, both in the striatum and beyond, including the brainstem, diencephalon, basal ganglia, hippocampus and cerebral cortex, whereas region-specific changes were found in all three major monoamine systems. These alterations were mostly in the levels of neurotransmitter metabolites, suggestive of changes, mostly an increase, in monoamine release. During exploration, both dopamine and serotonin release increase in A9 and A10 cell terminal fields in the dorsal striatum and nucleus accumbens [Broderick and Phelix 1997], and this is expected to support an active approach toward the environment. Novelty stimuli also increase dopamine release in the hippocampus from the nerve terminals originating from the VTA [Titulaer et al. 2021]. An increase in serotonin metabolism in the frontal cortex may help to respond to anticipatory cues [Merali et al. 2004] and reduce impulsive tendencies that disable persistent responding [Harro and Oreland 2016], whereas the increase in noradrenergic neurotransmission in the hypothalamus may serve as an optimization of stress response to novelty [Ma and Morilak 2005]. Interestingly, striatal GDNF overexpression affected dopamine and serotonin systems in the raphe area only in HE-rats, suggesting a neurobiological difference in the raphe between LE- and HE-rats. Thus, relevant changes in monoaminergic neurotransmission were elicited by striatal GDNF overexpression along the rostrocaudal axis of the brain. These were possibly necessary for the trait conversion but not sufficient, as these alterations mostly also occurred in the few LE-rats that did not become highly explorative.

Individual differences in novelty-related behaviour have a variety of neurobiological correlates [Pawlak et al. 2008; Harro 2010], but in the HE/LE-rat model a consistent finding is low extracellular striatal dopamine level in the LE-rats as measured by microdialysis [Mällo et al. 2007; Alttoa et al. 2009; O’Leary et al. 2016]. GDNF overexpression reduced extracellular dopamine levels in the HE-rats to the level of LE-rats, and had no effect in the LE-rats, both at baseline and after amphetamine treatment. This suggests that extracellular dopamine availability *per se* is not critically important for the exploration trait. However, in these experiments, a few LE-rats were GDNF non-responders, and a separate analysis revealed that in GDNF responders a tendency of higher extracellular dopamine levels was present, while in GDNF non-responders, dopamine levels were significantly reduced. These results suggest a specific alteration in the regulation of synaptic dopamine in GDNF-responder LE-rats, which includes the increased dopamine transporter availability as demonstrated by *in vivo* single photon emission computerized tomography (SPECT). To the extent the LE-rat represents a depression vulnerability model, these findings are consistent with the fairly reproducible finding of lower *in vivo* DAT availability in clinical depression [Mizuno et al. 2023], and with what may be even more relevant, the negative correlation of anxiety and psychomotor retardation in depressed patients with their DAT availability as measured by ^123^I-FP-SPECT [D’Onofrio et al. 2024]. In cell culture, application of GDNF enhances glycosylation and membrane trafficking of dopamine transporters and by this means appears to alleviate the behavioural deficits in the MPTP mouse model [Chengcheng et al. 2024]. DAT availability differences between LE- and HE rats are not explained by higher levels of gene expression in the latter, because in our previous studies, Western blotting did not reveal a significant difference in protein levels (unpublished) and in the present study, DAT mRNA levels were neither different between control LE- and HE-rats nor affected by GDNF overexpression. Instead, the transporter availability is likely to be the subject of membrane trafficking, known to impact dopamine-dependent behaviours [Bolden et al. 2025]. On the other hand, the GDNF non-response may have variable determinants, as lower striatal extracellular 5-HT levels and lower tyrosine hydroxylase levels in these animals suggest. It may be worth studying whether co-administration of serotonin reuptake inhibitors would lead to a synergistic effect.

RNA sequencing revealed extensive changes in gene expression in the striatum after AAV1-GDNF treatment. These appeared to be much more extensive in HE-rats as compared to LE-rats, but this difference represented an issue of statistical power - with regard to gene expression landscape, LE-rats are much more divergent than the HE-rats. This suggests that a variety of neurobiological mechanisms underlies the low exploratory phenotype. This appears similar to neuroticism-related psychiatric traits in humans that have multiple disease trajectories [Xia et al. 2024]. Nonetheless, GDNF overexpression can shift the low exploratory phenotype into a high exploratory one in the majority of these neurobiological types of low exploration. The pathways most consistently involved in this “personality change” include the oxytocin signalling pathway and the calcium signalling pathway, both implicated in anxiety regulation [Jurek and Neumann 2018; Montanez-Miranda et al. 2023]. We have further examined, in independent experiments, the expression of a few candidate genes that appeared in the hypothesis-free search to be altered by GDNF overexpression in LE-rats. The top gene in terms of statistical significance, *Slc18a1* or *Vmat1*, encodes the vesicular monoamine transporter type 1 (VMAT1), which is less common in CNS than VMAT2, but also expressed in the brain [Hansson et al. 1998], including the striatum [Ibanez-Sandoval et al. 2010], and plays a role in the accumulation of monoamine neurotransmitters in synaptic vesicles with a relatively higher affinity for serotonin [Brunk et al. 2006]. The highly conserved 136Thr locus of VMAT1 has human-unique gain-of-function substitutions under positive selection [Sato and Kawata 2018], these being associated with lower frequency or expression of bipolar disorder [Lohoff et al. 2006], anxiety-related personality traits [Lohoff et al. 2008], and anxiety disorders and depression [Vaht et al. 2016]. Humanized substitutions in *Vmat1* in mice also reduced anxiety-like behaviour [Sato et al. 2022]. Thus it appears that increased expression of *Vmat1*, possibly leading to higher efficacy of monoamine storage, is another mechanism in dopamine neurons responsible for the conversion of LE-rats into highly exploratory animals by GDNF overexpression. Next in the list of differentially expressed genes by GDNF overexpression in LE-rats was *Pdzd7*, a gene that has primarily been associated with hearing function, but possibly also contributing to brain development [Badshah et al. 2022]. An example of downregulation by GDNF overexpression in LE-rats to the level of HE-rats was provided by *P2rx6*. *P2rx6* encodes the ATP-gated ion channel, P2X6 purinoceptor, that has been found on GABA-ergic neurons in the substantia nigra and striatum, with increased expression after a 6-OHDA lesion as a possible compensatory reaction to low dopaminergic input [Amadio et al. 2007]. Hence the higher level of *P2rx6* expression in the LE-rats, and its downregulation by GDNF overexpression, may respectively reflect the lower dopaminergic state in LE-rats and its correction by GDNF. We were particularly intrigued by the finding of increased expression of *Mettl3* in LE-rats after GDNF overexpression. *Mettl3* encodes the main RNA m^6^A methyltransferase in mammalian cells, and RNA m^6^A methylation regulates splicing, transport, stability and translation of mRNA [Zaccara et al. 2019]. RNA m^6^A methylation participates in response to stress [Engel et al. 2018], and we have recently found that the first-in-class METTL3/METTL14 complex activating compound, CHMA1004, that increases RNA m^6^A methylation [Selberg et al. 2019], has a psychopharmacological profile of an activating and anxiolytic drug [Kanarik et al. 2025]. While the STRING analysis did not detect any significantly enriched network among the genes differentially expressed between the LE-GFP and LE-GDNF groups, *Mettl3* was found to be associated with two other such genes, and of these the effect of GDNF overexpression was confirmed for *Zfp692*. *Zfp692*, alternatively *Arebp* or *Znf692*, encodes the zinc finger protein 692, a transcription factor involved in immune traits [Zhang et al. 2015] and gluconeogenesis [Shirai et al. 2017] that serves as a scaffold in ribosome biogenesis [Lafita-Navarro et al. 2023]. Thus, converging evidence supports the involvement of RNA m^6^A methylation in selecting non-anxious, active behavioural strategies. Altogether, these experiments independently confirmed that expression of multiple genes is altered in striatum by GDNF overexpression, that may serve as the contributors to the conversion to high exploratory phenotype.

Thus, while the LE-rats have a large variety of neurobiological anxiety markers throughout the brain [Alttoa et al. 2007; Alttoa et al. 2010; O’Leary et al. 2016], striatal GDNF overexpression can elicit changes in neural circuits that lead to a conversion of a behavioural vulnerability trait into a more resilient one. We conducted a large-scale histochemical mapping of cytochrome oxidase activity in order to assess neuronal activity throughout the brain. Cytochrome c oxidase activity reflects overall neural functional activity similarly to 2-deoxyglucose uptake, but reflects activities over longer periods [Hevner et al. 1995], thus fitting to the studies on behavioural traits. Oxidative metabolism was reduced in the dorsomedial striatum by GDNF overexpression, and the strong negative correlation that existed in LE-rats between COX activity in this brain region with that in the dentate gyrus and zona incerta was abolished. Exposure to novel environments elicits a large immediate early gene response in the dorsomedial striatum, that habituates with repeated exposures [Struthers et al. 2005], and it is conceivable that the low exploratory phenotype is maintained by these relationships with other brain areas critically involved in novelty-related behaviour. Synaptic potentiation in the dentate gyrus occurs during exploration in order to facilitate learning about the novel environment [Moser et al. 1993], and interaction between the striatum and the hippocampus has a role in the emerging vulnerability to anxiety disorders during adolescence especially owing to the gains in attention regulation during this period [Lago et al. 2017]. Early developmental differences in cell proliferation have been reported between rats with low vs. high novelty-related activity [Clinton et al. 2011], and neuroplastic changes in the dentate gyrus have previously been found to occur together with a reduction of trait anxiety after deep brain stimulation [Schmuckermair et al. 2013]. Neural plasticity in the dentate gyrus has been found to depend on the integrity of dopaminergic neurotransmission [Suzuki et al. 2010]. Opposite to its effect in the striatum, GDNF overexpression increased oxidative metabolism in VTA. Balance of approach vs. avoidance behaviour has been linked with VTA neurons that receive prominent input from the dorsomedial striatum and exhibit substantial output to zona incerta [Wang et al. 2024]. In turn, zona incerta is thought to serve as a key communication hub of circuits that are dysfunctional in Parkinson’s disease, and deep brain stimulation of the zona incerta can alleviate this disorder by normalizing striatal output neuron function [Ossowska 2020]. Zona incerta integrates multiple neural circuits to support motor behaviours and survival-associated activities, and its stimulation suppressed fear generalization [Venkataraman et al. 2019] and may alleviate core symptoms of obsessive-compulsive disorder [Saluja et al. 2024]. It is critically involved in appetitively motivated behaviour [Wang et al. 2020], and appears to organize novelty-seeking behaviour in order to reduce uncertainty, interacting with dopamine neurons [Monosov et al. 2022; Monosov 2024]. GDNF overexpression altered the co-variation of intensity of oxidative metabolism in the dentate gyrus and zona incerta with that in the dorsal raphe and locus coeruleus, respectively, whereas activity co-variation of these regions with the ventrolateral part of periaqueductal grey matter was shifted from a negative to a positive correlation. The locus coeruleus is instrumental in immediate response to environmental novelty [Svensson 1987], including dopamine-dependent memory consolidation [Yamasaki and Takeuchi 2017], and lesioning of its projections has been demonstrated to severely impair novelty-related behaviour in the exploration box task [Harro et al. 1995]. In humans, higher signal intensity in the locus coeruleus corresponds to higher openness to experience, and higher IQ [Plini et al. 2024]. All this is compatible with the lower oxidative metabolism in LE-as compared to HE-rats, and the increase after GDNF overexpression, found in the present study. GDNF overexpression also led to positive co-variation of oxidative metabolism of both locus coeruleus and dorsal raphe with the ventrolateral part of the periaqueductal grey matter. Neurons of the ventrolateral periaqueductal grey matter are known to become active after both external fearful cues [Fendt 1998] and anxiogenic drug treatment [Singewald and Sharp 2000], and the ventrolateral periaqueductal grey matter is part of the network for appetitive/aversive prediction error together with the mesolimbic dopaminergic circuit, the amygdala, hippocampus and locus coeruleus [Iordanova et al. 2021]. Neurons in the ventrolateral periaqueductal grey and dorsal raphe, while activated by stressful stimuli, exert a synergistic anti-anxiety effect [Zhang et al. 2024]. Altogether these alterations in the regional activity co-variation produced by striatal GDNF overexpression suggest that functional reorganization in a network of brain regions implicated in active exploration of the environment, comprising the striatum, hippocampus, zona incerta, locus coeruleus, dorsal raphe and ventrolateral periaqueductal grey, underlies the profound change in the exploratory behaviour of LE-rats after striatal GDNF overexpression and repeated exposure to an environment that had previously been persistently aversive.

Another interesting finding concerns the co-variation of oxidative metabolism in the medial and cortical amygdaloid nuclei with that in the claustrum. Amygdalar activity, as measured by glucose metabolism, is a trait-like feature of the brain [Fox et al. 2008], and it co-varies with behavioural inhibition [Shackman et al. 2013]. The canonical model of anxiety has mostly focused on the central and basolateral nuclei [Fox and Shackman 2024], but other nuclei of the corpus amygdaloideum are also involved in ethologically relevant threat processing. While the medial amygdala is mostly recognized as a hub of anxiety regulation in social contexts, there is also evidence for its implication in anxiety and passive coping response to novel physical environment [Moreno-Santos et al. 2021], and electrical kindling of the medial amygdala has reduced exploration in an anxiety test [Adamec 1990]. Both the medial and cortical amygdala are involved in response to alarm pheromone detection in the rat [Kobayashi-Sakashita et al. 2023], and neurons of the cortical amygdala are responsive to intruder threat [Caffrey et al. 2010]. In turn, the claustrum has reciprocal connections with various cortical regions, is believed to serve the role of a multisensory integrator for a complex performance [Crick and Koch 2005], supports cognitive control under demanding conditions [White et al. 2020], and its activation by stressful stimuli can elicit anxiety-related responses [Niu et al. 2022]. It might be of relevance that the co-variance of oxidative metabolism in the claustrum with that of zona incerta and locus coeruleus in LE-rats was also shifted more HE-like by GDNF overexpression, even though this change did not reach statistical significance. However, the claustrum is directly connected to the corpus striatum [Milardi et al. 2015; Borra et al. 2024] and electrical stimulation of the claustrum in dogs can elicit both active and passive defensive behaviours [Vakolyuk et al. 1983]. In a joint analysis of COX activity mapping in four genetic mouse models of depression, the claustrum emerged as one of the top regions discriminating the genetically modified mice from the wildtype [Matrov et al. 2019]; it has been found less active in socially defeated rats [Kanarik et al. 2011], and in depressed humans in resting-state conditions [Fitzgerald et al. 2008]. A study in rhesus macaques has identified a network comprising the striatum, amygdala and claustrum being involved in processing object novelty, value and their combination [Ghazizadeh et al. 2020].

Yet another finding of interest was the downregulation of oxidative metabolism by GDNF overexpression in the BNST, and the appearance of BNST activity co-variation with the nucleus accumbens and with claustrum. The role of the BNST in anticipation of threat, stress response and anxiety disorders is well established [Lebow and Chen 2021]. Interestingly, in human post-traumatic stress disorder the functional connectivity of BNST with the ventral striatum and claustrum is altered [Rabellino et al. 2018], and dual-target combined stimulation of the BNST and nucleus accumbens was recently found effective in treatment-resistant depression [Wang et al. 2024]. Altogether it appears that striatal GDNF overexpression leading to the increase of exploratory behaviour in LE-rats and making the phenotype similar to the HE-rat is related to rearrangement of neural integration in specific circuits involving subregions of limbic and brainstem nuclei and claustrum, so that the functional coupling of these becomes HE-like.

In sum, bilateral overexpression of GDNF in the striatum of rats with predisposition of passive coping in novel environments could elicit a gradual but profound conversion of their exploration phenotype into an active coping style. This was based on multiple alterations in striatal transcriptomic landscape and at dopaminergic synapse, and associated with broad reorganization in regional cerebral activity patterns. Given that the LE-rat is less sensitive to conventional antidepressant treatments, GDNF deserves attention as a treatment option for chronic, recurrent and treatment-resistant mood and anxiety disorders.

## Methods

### Animals

Male Wistar rats (Scanbur BK AB, Sweden or Harlan Laboratories, the Netherlands) were housed 3-4 per cage in standard transparent polypropylene cages under controlled light cycle (lights on from 08:00 h to 20:00 h) and temperature (19–21 °C), with free access to tap water and food pellets (v1534-000 universal maintenance diet, ssniff Spezialdiäten GmbH, Soest, Germany). The experiments were in accordance with EU legislation (directive 2010/63/EU) and the studies were approved by the Ethical Committee for Animal Experiments of the Estonian Ministry of Rural Affairs (permission no 149; 27.09.2019) and the Finnish Project Authorisation Board (permission no ESAVI/13959/2019).

### Exploration phenotyping

This was carried out at age 2-3 months using the exploration box test [Otter et al. 1997] as previously described [Mällo et. al. 2007] (Fig. 1a). The exploration box was made of metal and comprised a 0.5 m x 1 m open area (the height of side walls 0.4 m) and a 20 cm x 20 cm x 20 cm compartment attached to one of the shorter sides of the open area. The open area was divided into eight equal-sized squares, and four objects, three unfamiliar (a glass jar, a cardboard box, and a wooden handle) and one familiar (a food pellet), were placed onto certain squares. The location of the objects remained the same throughout the experiment. The floor of the small compartment was covered with wood shavings. The open area was directly accessible from the small chamber through an opening (20 cm x 20 cm). The exploration box test was initiated by placing the rat into the small compartment which was then covered with a lid for the duration of the test. The following parameters were recorded by an observer: (a) latency of entering the open area with all four paws; (b) entries into the open area; (c) line crossings, (d) rearing; (e) exploration of the three unfamiliar objects in the open area; (f) the time spent exploring the open area. To provide an index of exploration considering the elements of both inquisitive and inspective exploration, the scores of line crossings, rearing and object investigations were summed for each animal. This measure was used throughout the presented analyses because in naive rats all measures are strongly correlated and the administration of viral vectors did not alter this. Objects investigation refers only to unfamiliar objects, because without any intervention, the familiar object is ignored. After each animal, the exploration box was wiped clean with a wet tissue. A single test session lasted 15 min and the experiments were carried out under dim light conditions (3-7 lx in the open area). All animals were exposed to the exploration box on two consecutive days. The classification of rats into groups of high or low explorers (HE or LE, respectively) was based on the sum of the exploratory events on the second exposure to the exploration box test. Activity distribution measured in that way strongly deviates from normal (Fig. 1b). LE-phenotype was defined as the Day 2 total exploration score <50 and no decline as compared to the first day, but in these experiments the majority of the LE-rats did not emerge from the starting box at all. Exploration was tested again two weeks after administration of vectors.

### Generation of viral constructs

To produce the self-complementary AAV (scAAV) vector expressing human pre-α-pro-GDNF or pre-ß-pro-GDNF, the cDNA fragment encoding human pre-α-pro-GDNF or pre-ß-pro-GDNF was produced by PCR using pAAV-pre-α-pro-GDNF or pAAV-pre-ß-pro-GDNF [Lonka-Nevalaita et al. 2010] as a template. PCR was performed with Phusion Hot-Start polymerase (ThermoFisher Scientific, Waltham, MA). PCR products were purified and digested by BamHI and NotI restriction enzymes (ThermoFisher Scientific, Waltham, MA) and ligated into a pscAAV-CMV vector using T4 DNA ligase (ThermoFisher Scientific, Waltham, MA). The plasmid pscAAV-CMV was obtained by cutting out the eGFP insert from pscAAV-CMV-eGFP using BamHI and NotI restriction sites. The cloned construct was verified by DNA sequencing. Primers used for cloning of pre-α-pro-GDNF into pscAAV-CMV were forward 511-TAGGATCCATGA AGTTATGGGATGTCGTGG-311 containing BamHI restriction site and reverse 511-TAGCGGCCGCTCAGATACATCCACACC TTTTA-311 containing NotI restriction site.

The self-complementary AAV vectors, scAAV-pre-α-pro-GDNF, scAAV-pre-β-pro-GDNF and scAAV-CMV-eGFP were packaged as serotype 1 [Howard et al. 2008], then purified and titered as described previously [Henderson et al. 2014]. The titer for the vector was 7.40 × 1013 vg/ml. AAV vector preparation was conducted by the Optogenetics and Transgenic Technology Core, NIDA IRP, NIH, Baltimore MD, USA.

### Intrastriatal administration of viral vectors

This was done as previously described [Penttinen et al. 2018]. All stereotaxic surgeries were performed under ketamine/medetomidine anaesthesia (60 and 0.5 mg/kg i.p., respectively), atipamezole (1 mg/kg, s.c.) was used for anaesthesia reversal. Lidocaine (1%, appr. 1 mg/kg, i.d.) was used for perioperative and meloxicam (1-2 mg/kg, s.c., for three days) as post-operative analgesic. For the viral vector injections, LE- and HE-rats were randomly allocated to treatment groups. Two μl of scAAV1-pre-α-pro-GDNF or scAAV1-eGFP were injected into the striatum bilaterally. The injection coordinates were A/P: +0.7 mm and L/M ±3.0 mm from bregma; D/V -6.0 from dura mater. Injections were done at a rate of 0.5 μl/min. The microinjection needle was kept in place for additional 4 min to avoid backflow of the solution.

### Immunohistochemistry

Right after SPECT imaging, and while the animals were still under isoflurane anaesthesia, they were transcardially perfused with phosphate buffered saline (PBS) and 4% paraformaldehyde (PFA) solution. Brains were removed and post-fixed overnight in 4% PFA at +4°C and transferred to sucrose series of 20 and 30% sucrose. The brains were cut in a freezing microtome in 40 μm thick sections in series of six. Free-floating sections were stained as previously described [Penttinen et al. 2018]. In brief, the sections were washed and treated with 0.3% hydrogen peroxide solution. After incubation in the blocking solution (4% bovine serum albumin and 0.1% Triton X-100 in PBS) the sections were incubated with the primary antibody overnight at +4 °C. Next, the sections were incubated with biotinylated secondary antibodies (anti-rat, anti-mouse, or anti-rabbit, Vector Laboratories, Burlingame, CA) and the staining was reinforced with avidin-biotin-complex (Vector Laboratories, Burlingame, CA) and visualized with 311,311 diaminobenzidine. The stained images of the brain slices were obtained by Aizure Biosystems, Sapphire FL Biomolecular Imager IS4000.

For the TH immunofluorescence staining in the striatum, the images were converted to an 8-bit gray scale format. This allowed for the measurement of the Mean Gray Value parameter of the selected areas, which were defined using freehand selection and the ROI manager in the ImageJ program.

### Tissue levels of GDNF by ELISA

Enzyme-linked immunosorbent assay (ELISA) was used to reveal the level of expression of GDNF in the brains of animals in association of the treatment response. Immediately after sacrifice and dissection, the fresh brain snap-frozen brain samples (striatum or nucleus accumbens) were weighed, and RIPA buffer (150 mM NaCl, 1% NP-40, 0.5% sodium deoxycholate, 0.1% SDS, 50 mM Tris; pH 8.0) with protease inhibitor cocktail (cOmplete™, Roche) was added directly to frozen brain sample. Ten µl of buffer was used for 1 mg of tissue. Brain tissue was homogenized in a cold room at 4 °C by sonication with Sonopuls HD2070 homogenizer (Bandelin Electronic, Germany) 5 times (0.5 s on, 0.5 s off for 10 s, then left to stand for 30 s and repeated). Sonication was followed by incubation and rotation of samples in 1.5 ml tubes for 20 min at cold room. Homogenates were centrifuged at 16,100 g for 20 min at 4 °C and supernatant was collected and aliquots were made, and stored at -80 °C. Protein quantification was done with Pierce BCA Protein Assay Kit (Thermo Scientific) and absorbance was measured in 96-well clear F-bottom plates (Greiner) using FLUOStar Optima plate reader (BMG Labtech, Germany). GDNF levels were determined from the supernatants by commercial enzyme-linked immunosorbent assay (ELISA) kit according to the manufacturer’s recommendations (R&D Systems, Minneapolis, MN).

### Behaviour of HE-rats after 6-OHDA lesion

A restricted dopaminergic lesion was inflicted with intrastriatal injections of 6-hydroxydopamine (6-OHDA) according to a protocol modified from Lindholm et al. [2007]. 6-OHDA was dissolved in saline with 0.02% ascorbic acid on ice and injected bilaterally into striata. The solution (1.5 µl, containing 2 µg of 6-OHDA) was injected per hemisphere with a rate of 0.5 µl/min, the coordinates of the injection were: AP +0.7; ML 3.0; DV -6.0 (Paxinos and Watson 1986). Baseline exploratory activity was measured at 11 weeks of age, followed by 6-OHDA injection at 14 weeks of age (n=13 for 6-OHDA and n=12 for control group). After a 2-week recovery period animals were tested seven times in the exploration box, the first three postoperative test were two weeks apart, the latter four tests two months apart from each other (see Methods, Exploration phenotyping for the description of the exploration test). In between the first and second exploration test, locomotor activity was measured, with the lighting conditions identical to the measurement of explorative behaviour. Animals were habituated with the testing room for 30 min, after which locomotor activity was measured in a cage identical to the animals’ home cage. The floor of the cage (area 40×50 cm), covered with bedding, was divided into six segments, crossings of segment borders in a 30-min test were counted from video recordings.

### Ex vivo measurement of monoamines and metabolites

Monoamines and their metabolites were assayed by HPLC with electrochemical (amperometric) detection as described previously [Kõiv et al. 2019]. Rat brain tissues were homogenized with an ultrasonic homogenizer (Bandelin Sonopuls, Bandelin Electronic, Berlin, Germany) in ice-cold solution of 0.1 M perchloric acid (30 μl/mg for the nucleus accumbens, amygdala and frontal cortex; 50 μl/mg for striatum) containing 5 mM of sodium bisulfite and 0.4 mM EDTA to avoid oxidation. The homogenate was then centrifuged at 14,000 rpm for 10 min at 4 °C. Aliquots (10 μl) of the supernatant obtained were chromatographed on a Luna C18(2) column (150×2 mm, 5 μm). The separation was done in isocratic elution mode at column temperature of 30 °C, using the mobile phase containing 0.05 M sodium citrate buffer at pH 3.7, 0.02 mM EDTA, 1 mM KCl, 1 mM sodium octanesulfonate and 7.5% acetonitrile. The chromatography system consisted of an isocratic pump (HP1100, Agilent, Waldbronn, Germany and Shimadzu LC-20AD, Japan), a temperature-regulated autosampler (HP1100, Agilent, Waldbronn, Germany and Shimadzu SIL-20AC, Japan), a temperature-regulated column compartment and an electrochemical detector (HP 1049 Agilent, Waldbronn, Germany and Antec Decade II, Netherlands) with a glassy carbon electrode (HP flow cell and VT-03 flow cell, Antec, Netherlands). The measurements were done at an electrode potential of +0.7 V versus the Ag/AgCl reference electrode. The limits of detection at signal-to-noise ratio=3 were as follows (expressed as pmol/mg tissue for each): 0.08 for DA, 0.10 for homovanillic acid (HVA), 0.05 for 3,4-dihydroxyphenylacetic acid (DOPAC), 0.08 for 5-HT, 0.04 for 5-hydroxyindoleacetic acid (5-HIAA), 0.07 for noradrenaline (NA), 0.03 for normetanephrine (NMN), and 0.01 for 3-methoxytyramine (3-MT).

### In vivo microdialysis

In vivo microdialysis was carried out to reveal changes in extracellular neurotransmitter levels at baseline and after local depolarization or depolarization-independent release capacity, essentially as previously described [O’Leary et al. 2016]. The animals were anaesthetized with ketamine/medetomide anaesthesia (60 and 0.5 mg/kg i.p., respectively) and mounted in a Stoelting stereotactic frame. A self-made concentric Y-shaped microdialysis probe with 7 mm shaft length and 3 mm active tip was implanted into the left dorsal striatum according to the following coordinates: AP +0.7; ML +3.0; DV –7.0, according to Paxinos and Watson (1986). The dialysis membrane used was polyacrylonitrile/sodium methalyl sulphonate copolymer (Filtral 12; i.d.: 0.22 mm; o.d.: 0.31 mm; AN 69, Hospal, Bologna, Italy). Two stainless steel screws and dental cement was used to fix the probe to the scull. After the surgery, anaesthesia was reversed by administration of atipamezole (1 mg/kg, s.c.) and meloxicam (1-2 mg/kg, s.c.) was administered for post-operative analgesia. The rats were placed in 21 cm × 36 cm × 18 cm individual cages in which they remained throughout the microdialysis experiment. Rats were given about 24 h for recovery and microdialysis procedure was conducted in awake freely moving animals. The microdialysis probe was connected to a syringe pump (SP101, World Precision Instruments, Inc., Sarasota, FL, USA) and to a refrigerated microsampler (Univentor 820, Univentor Limited, BLB029 Bulebel Industrial Estate, Zejtun ZTN 3000, Malta) and perfused with Ringer solution (147 mM NaCl, 4 mM KCl, 1.2 mM CaCl_2_, 1.0 mM MgCl_2_, 1.0 mM Na_2_HPO_4_; pH 7.20–7.22) at a constant rate of 1.5 μl/min. Connections to the infusion pump and microfraction collector were made with flexible FEP tubing (i.d. 0.12 mm, AgnTho’s AB, Lidingö, Sweden). After connecting the animal to the microdialysis system, the perfusate was discarded during the first 60 min to allow stabilization. Five baseline samples were collected, followed by 30-min perfusion with Ringer solution that contained 50 mM KCl at the beginning of collection of the sixth sample. At the beginning of the 14th sample (two hours after the start of the 50 mM KCl), some animals received an i.p. injection of amphetamine (0.5 mg/kg, i.p.), after which another nine samples were collected. All samples were collected with 15-min intervals. Sample vials were prefilled with 7.5 μl of 0.02 M acetic acid to prevent oxidation of dopamine. Upon completion of the experiment the animals were deeply anaesthetized with isoflurane and decapitated; the brains were removed, immediately frozen in ice-cold methylbutane and kept at –80 °C. The brains were sectioned on a cryostatic microtome (Microm GmbH, Walldorf, Germany) and probe placements were determined according to the atlas of Paxinos and Watson [1986]. The quantity of dopamine and serotonin in the microdialysis samples was determined by high performance liquid chromatography with electrochemical detection. The chromatography system consisted of a Shimadzu LC-20AD pump and CBM-20A controller (Shimadzu Corporation, Kyoto, Japan), a Luna C18(2) 5 μm column (150 x 2 mm) kept at 30 °C and Decade II digital electrochemical amperometric detector (Antec Leyden BV, the Netherlands) with electrochemical flow cell VT-03 (2 mm GC WE, ISAAC reference electrode, Antec Leyden BV, the Netherlands). The mobile phase consisted of 0.05 M sodium citrate buffered to pH 5.3, 2 mM KCl, 0.02 mM EDTA, 3.5 mM sodium octyl sulfonate and 14% acetonitrile. The mobile phase was filtered through a 0.22 μm pore size filter (type GV, Millipore, USA) and was pumped through the column at a rate of 0.2 ml/min. 5-HT and DA eluted from the column were measured with a glassy carbon working electrode maintained at a potential of +0.4 V versus Ag/AgCl reference electrode. Data were acquired using a Shimadzu LC Solution system.

### Single photon emission computerized tomography (SPECT)

Nine weeks after viral transfection and two weeks after the last exploration box test, 28 rats (290-350 g), in random order received an intravenous injection of 40 to 50 MBq ^123^I-labeled N-(3-fluoropropyl)-2beta-carbomethoxy-3beta-(4-iodophenyl)nortropane ([^123^I]-FP-CIT; GE, Healthcare, Finland). Two hours later, the rats were anaesthetised with isoflurane (4% induction, 1.7% maintenance), and imaged with a four-head nanoSPECT/CT (Mediso Ltd., Budapest), featuring 2.5-mm multi-pinhole rat apertures. Imaging was performed blind of the identity of each rat group, under isoflurane anaesthesia (2% to 3%), and the body temperature was maintained at 36±0.5 °C using a heated animal bed (Minerve, France).

Brain SPECT images were collected in 24 projections using a time per projection of 90 s, resulting in a total acquisition time of 25 min. CT imaging was carried out with a 45-kVp tube voltage in 180 projections. SPECT images were reconstructed, and fused, with CT datasets with Fusion software (Mediso, Hungary). Reconstructed SPECT images were reoriented and analysed with the same software using CT data as a reference. Elliptical volume of interest was defined around the striatum of an intact rat and used in all subsequent analyses. The mean striatal activity (activity-to-volume ratio) was counted and corrected for background activity from the cerebellum. Striatal volumes were considered for DAT binding potential, while mid-brain volumes as SERT binding potential. SPECT research team was unaware of the group allocation of the animals.

### RNA extraction and mRNA sequencing

RNA was extracted from rat striatal samples using FavorPrep^TM^ Tissue Total RNA Mini Kit (FATRK 001-2, Favorgen, Taiwan) according to the manufacturer’s protocol. Briefly, the FARB lysis buffer, containing ß-mercaptoethanol, was added to the tissue sample, grinding was performed with a micropestle followed by homogenization through a 20-G needle syringe. The lysate was cleared by using the filter column followed by addition of one volume of 70% ethanol. The sample was then transferred to the FARB Mini Column, washed, and dried by centrifugation at 18,000 g for three minutes. Total RNA was eluted in 30 µl of nuclease-free water, concentration was measured by NanoDrop 1000 spectrophotometer (ThermoFisher Scientific Inc., Waltham, MA, USA) and stored at -80°C. Sequencing was performed by Novogene Co., Ltd. (Beijing, China). Quality control and preprocessing of the fastq files, containing raw reads from Illumina 150bp paired-end sequencing, was conducted with fastp v0.23.2 using default parameters [Chen et al. 2018]. Strand specific alignment to the rat rn7 genome was carried out using HISAT2 v2.2.1. (paired end, Strand -FR).

FeatureCounts v1.6.4 [Liao et al. 2014] was used for transcript counting using *Rattus_norvegicus*.mRatBN7.2.108.gtf annotation from Ensembl database as a reference [Martin et al. 2023]. Differential gene expression between pairwise groups was performed with DeSeq2 v22.11.40.6 [Love et al. 2014] using posCounts method for size factor estimation. Genes with p-value smaller than 0.05 were considered differentially expressed. SynGO portal was used for ID conversion [Koopmans et al. 2019]. GO enrichment analysis was performed with DAVID 2021 [Sherman et al. 2022]. The analysis included 5 rats per group. The Volcano plot was created using R ggplot2 based tool in Galaxy (Version 0.0.3) using output from DESeq2.

### Quantification of mRNA levels by realllltime quantitative PCR (RTlllqPCR)

Total RNA was isolated using RNeasy Mini Kit (QIAGEN, #74106). RNA purity was validated by optical density (OD) absorption ratio (OD 260 nm/280 nm) using IMPLEN Nanophotometer (Germany). Then, total RNA was reverse transcribed into cDNA in a 20 μL reaction volume using RevertAid First Strand cDNA Synthesis Kit (Thermo Scientific, #K1622) according to the manufacturer’s instructions. Real-time quantitative PCRs were performed using Power SYBR® Green PCR Master Mix (Applied Biosystems, #4368706) and StepOnePlus™ real-time PCR system (Applied Biosystems). The specific primer sequences used for target genes, and reference gene (*GAPDH*) are shown in Suppl. Table 2 and were ordered from Invitrogen, Thermo Fisher Scientific (USA). Genes were amplified in triplicated RT-qPCR runs using a 96-well plate loaded with 1 μL of cDNA and either 250 or 500 nM of each forward and reverse primer in a final volume of 10 μL. RT-qPCR cycling conditions consisted of an initial step of 95 °C for 10 min, and 40 cycles of 95 °C for 15 s and 60 °C for 1 min. The amount of total mRNA present in each reaction was normalized to *GAPDH* in parallel samples. Results were expressed as relative levels of mRNA, referred to as control samples (“LE, GFP, Control”) and calculations were performed according to the comparative CT method (2–ΔΔCT) [Livak and Schmittgen 2001).

### Cytochrome c oxidase staining and image analysis

Brains were flash-frozen in isopentane on dry ice followed by storage at -80 °C. Sections of 40 µm were sliced on a Leica CM1520 cryostat onto VWR Superfrost Plus slides and stored at -80 °C until use. Cytochrome c oxidase staining and image analysis were performed as described previously [Matrov et al. 2020], the protocol being a modification of earlier work of other authors [Gonzalez-Lima and Cada 1998]. Coronal brain sections were pre-incubated for 10 min with 0.0275% cobalt chloride (w/v) and 0.5% dimethyl sulphoxide (v/v) in 0.05 M Tris buffer with 10% sucrose (w/v) adjusted to pH to 7.4 with 0.1% HCl (v/v). The sections were then incubated for one hour at 37 °C in a solution consisting of 0.05% 3,3 - diaminobenzidine tetrahydrochloride (AppliChem), 0.0075% cytochrome c (Sigma), 5% sucrose, 0.002% catalase (Sigma) and 0.25% dimethyl sulphoxide (v/v) in sodium phosphate buffer (pH 7.4). Image analysis was conducted using the Image J 1.34s freeware on the blue channel (resulting from a RGB split) of the background subtracted image. Brain regions were detected from stained images according to the rat brain atlas (Paxinos and Watson, 2007). Enzyme activity was derived from optical density measurements of a histochemical reaction product within each brain region. The optical density values were converted to enzyme activity by using external standardization: sections made of brain homogenate with spectrophotometrically measured enzyme activity were included in all incubation baths.

## Statistical analysis

Data are presented as mean ± s.e.m. Exploration data were submitted to Shapiro-Wilk normality test. Data of intervention experiments were analyzed with one- or two-way analysis of variance (ANOVA) including repeated measures as appropriate, with Fisher’s LSD post hoc comparison), or with Student’s t-test. Dependent on data distribution, Pearson or Spearman correlation analysis was applied. Pearson’s correlations were compared with the z-test for r-to-z transformed coefficients with cocor, a software package for R [Diedenhofen and Musch 2015]. To compare the samples according to their general genetic patterns, the principal component analysis with standardised gene expression data was performed. To measure the genetic differences between rats, the Bray-Curtis distances based on standardised gene expression data were calculated. The distributions of inside treatment and exploring group Bray-Curtis distances were visualized with box plot and the average inside group distances between two exploring groups by treatments were compared with t-test. The R package vegan was used to calculate Bray-Curtis distances. Whenever possible, experiments and measurements were blinded to group allocation during analysis. Statistical analyses were performed using Prism v.10 (GraphPad Software) and SigmaPlot v.15.0 (Inpixion).

## Acknowledgements

This study was funded by the Estonian Research Council (PRG1213) to J.H., the Sigrid Jusélius Foundation grant to M.S., and the TLU Centre of Excellence in Behavioural and Neural Sciences and the School of Natural Sciences and Health Research Fund (R.S.). J.H. was supported by Visiting Professorship of the Sigrid Jusélius Foundation. T.O. was supported by the European Union through Horizon 2020 research and innovation programme under grant number 810645. I.T. was funded by EU Marie Sklodowska Curie postdoctoral fellowship 101068830. M.S. was supported from the Cure Parkinson’s Trust, UK grant (project code MS02). KLi was supported by “Dora Pluss” program funded by the European Union through the European Regional Development Fund, and Kristjan Jaak Stipend provided by the Estonian Ministry of Education and Research. Technical assistance of Sari Tynkkynen, Julia Johansson and Marju Vahter is gratefully acknowledged.

## Author Contributions

J.H. conceptualized the study with seminal input from M.S. M.N., K.Li., M.K., A.G.-H., A.O., I.T., T.O., P.L., M.A. and B.K.H. made essential contributions to methodology. B.H. supplied the vectors. M.N., K.Li., M.K., A.G.-H., A.O., K.La., P.K., S.S., I.P., S.I., S.K. and H.P. performed the experiments. M.N., K.Li., M.K., A-G.-H., A.O., K.La., P.K., S.S., I.T., T.O., T.K. and P.L. analyzed the data. J.H., M.A., R.S. and M.S. supervised the study. J.H. wrote the manuscript with input from all coauthors. J.H., M.A, R.S. and M.S. provided funding for the study.

## Supplementary Figures

**Suppl. Fig. 1:** Dopaminergic lesion by bilateral intrastriatal 6-OHDA injection (2 µg per hemisphere) to HE-rats reduced their high level of exploration only modestly. (a) Exploratory activity in the exploration box test. The first three postoperative trials were two weeks apart, the last four tests two months apart (6-OHDA effect F(1,23)=6.70; p<0.016; repeated measurement effect F(7,161)=14.88; p<0.0001) 1b: Locomotor activity on an open field, measured after the first postoperative exploration box test. *P < 0.05;**P < 0.01 compared to the vehicle-treated control group.

**Suppl. Fig. 2:** STRING database analysis of protein-protein interactions in the set of differentially expressed genes between the LE-GFP and LE-GDNF groups. Note the interaction of *Mettl3* with *Zfp692* and *Lrpprc*.

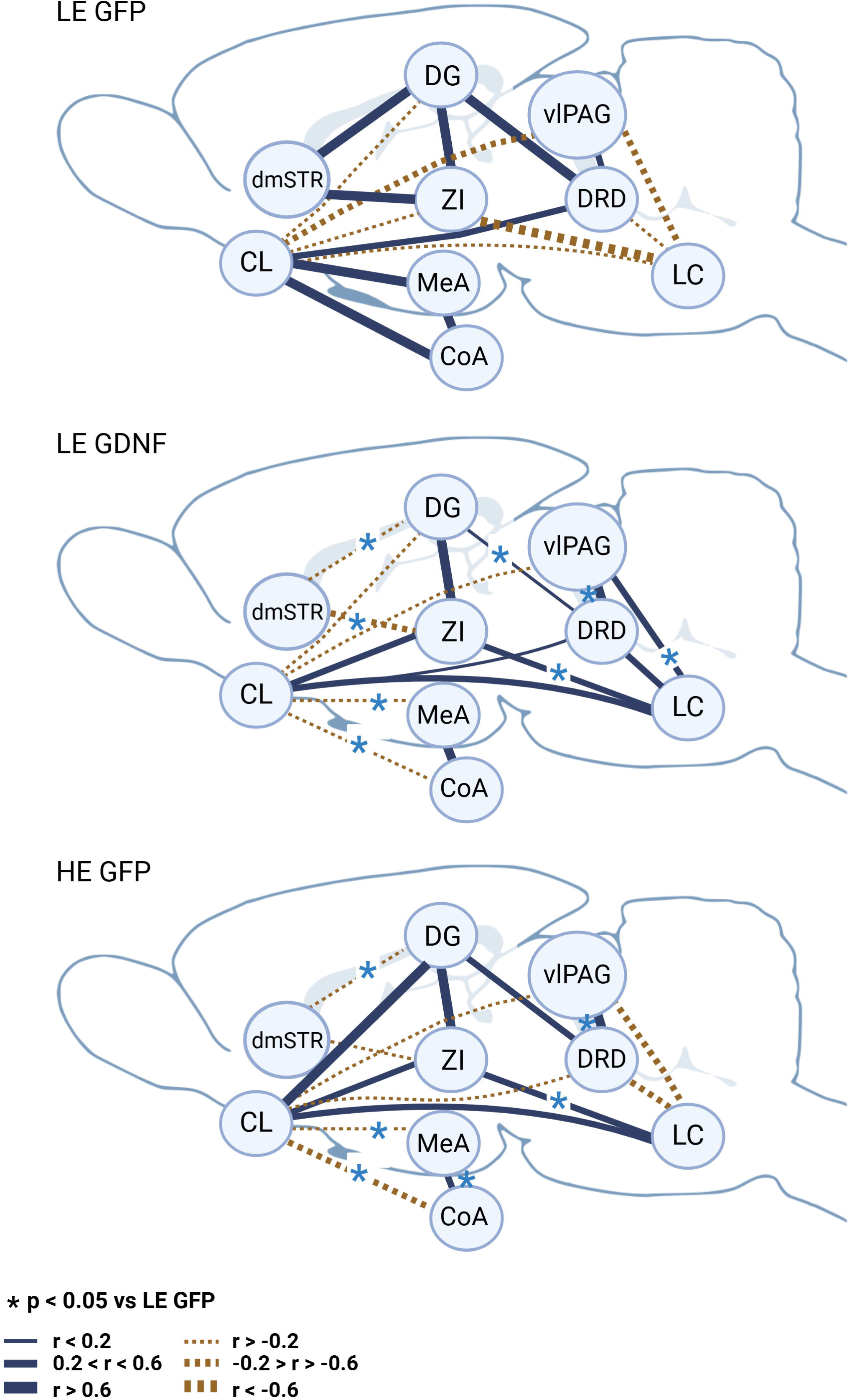

