## Supplemental Table 1 for "Experimental change in personality: Overexpression of GDNF in the rat striatum converts the low exploratory phenotype into highly explorative"

**Supplementary Table 1.** List of differentially expressed genes (p<0.05) based on RNA-seq analysis.

**LE-GFP vs LE-GDNF**

| **Gene symbol** | **Average norm read counts LE-GFP** | **Average norm read counts LE-GDNF** | **P-value** | **Fold Change** |
| --- | --- | --- | --- | --- |
| *Slc18a1* | 11,77 | 32,39 | 4,79E-05 | 2,75 |
| *Pdzd7* | 68,41 | 107,85 | 6,25E-05 | 1,58 |
| *Pdrg1* | 28,07 | 61,20 | 1,69E-04 | 2,18 |
| *Msantd2* | 107,38 | 149,46 | 1,99E-04 | 1,39 |
| *RGD1565987* | 15,30 | 5,54 | 2,93E-04 | -2,76 |
| *ENSRNOG00000006324* | 42,97 | 23,81 | 4,81E-04 | -1,81 |
| *ENSRNOG00000063514* | 40,05 | 77,68 | 7,58E-04 | 1,94 |
| *Phyhd1* | 125,97 | 183,16 | 7,64E-04 | 1,45 |
| *RGD1564941* | 12,13 | 27,28 | 7,90E-04 | 2,25 |
| *Ryr1* | 102,91 | 146,42 | 0,0010 | 1,42 |
| *AC135409.1* | 54,12 | 99,98 | 0,0011 | 1,85 |
| *Fabp4* | 1,89 | 8,48 | 0,0011 | 4,49 |
| *Arhgap4* | 85,17 | 123,03 | 0,0014 | 1,44 |
| *ENSRNOG00000070093* | 2,76 | 9,48 | 0,0019 | 3,43 |
| *Art4* | 10,10 | 20,75 | 0,0019 | 2,05 |
| *Slc16a4* | 0,75 | 5,89 | 0,0022 | 7,81 |
| *ENSRNOG00000071040* | 65,73 | 91,23 | 0,0022 | 1,39 |
| *ENSRNOG00000062681* | 1,16 | 6,75 | 0,0025 | 5,83 |
| *Cacng8* | 1026,47 | 865,09 | 0,0027 | -1,19 |
| *P2ry2* | 68,19 | 45,57 | 0,0027 | -1,50 |
| *Amer3* | 357,17 | 276,89 | 0,0031 | -1,29 |
| *Mgarp* | 1,93 | 7,38 | 0,0034 | 3,82 |
| *Dgka* | 635,54 | 771,36 | 0,0036 | 1,21 |
| *Snrnp70* | 2372,61 | 2839,75 | 0,0039 | 1,20 |
| *Fmod* | 67,85 | 41,63 | 0,0041 | -1,63 |
| *LOC120093114* | 29,67 | 52,18 | 0,0041 | 1,76 |
| *ENSRNOG00000070818* | 43,07 | 65,03 | 0,0045 | 1,51 |
| *Luc7l3* | 1439,34 | 1686,60 | 0,0048 | 1,17 |
| *Nr1i3* | 22,76 | 39,44 | 0,0052 | 1,73 |
| *LOC102550851* | 126,24 | 164,78 | 0,0055 | 1,31 |
| *Sgsh* | 88,25 | 60,50 | 0,0057 | -1,46 |
| *ENSRNOG00000065338* | 62,00 | 90,53 | 0,0059 | 1,46 |
| *Mon1b* | 62,04 | 42,98 | 0,0059 | -1,44 |
| *Nek5* | 50,86 | 77,22 | 0,0064 | 1,52 |
| *ENSRNOG00000069955* | 3,88 | 10,03 | 0,0067 | 2,59 |
| *Zfp161* | 350,49 | 285,14 | 0,0067 | -1,23 |
| *ENSRNOG00000063555* | 17,29 | 7,07 | 0,0068 | -2,45 |
| *Zfp958* | 136,90 | 196,07 | 0,0070 | 1,43 |
| *Ccer2* | 10,75 | 23,73 | 0,0071 | 2,21 |
| *RGD1561157* | 30,16 | 18,94 | 0,0074 | -1,59 |
| *ENSRNOG00000066393* | 8,41 | 18,26 | 0,0077 | 2,17 |
| *Emcn* | 95,13 | 126,72 | 0,0079 | 1,33 |
| *Clk1* | 1125,13 | 1386,36 | 0,0082 | 1,23 |
| *ENSRNOG00000067311* | 37,28 | 21,50 | 0,0083 | -1,73 |
| *Tmem129* | 226,88 | 279,85 | 0,0083 | 1,23 |
| *Tex9* | 74,61 | 102,34 | 0,0084 | 1,37 |
| *LOC100912291* | 4,57 | 1,26 | 0,0087 | -3,64 |
| *Esrra* | 66,92 | 46,71 | 0,0089 | -1,43 |
| *Pdzd9* | 22,65 | 37,52 | 0,0090 | 1,66 |
| *Hoxd1* | 5,00 | 1,18 | 0,0091 | -4,24 |
| *ENSRNOG00000068540* | 0,21 | 3,70 | 0,0091 | 17,31 |
| *AABR07022168.1* | 16,25 | 8,80 | 0,0098 | -1,85 |
| *Cenatac* | 113,08 | 148,01 | 0,0100 | 1,31 |
| *Plekhn1* | 131,93 | 173,52 | 0,0104 | 1,32 |
| *Stx2* | 139,29 | 177,05 | 0,0108 | 1,27 |
| *ENSRNOG00000033611* | 20,65 | 34,36 | 0,0109 | 1,66 |
| *Lsp1* | 53,92 | 78,87 | 0,0111 | 1,46 |
| *RGD1561381* | 11,10 | 23,58 | 0,0111 | 2,12 |
| *Zfp692* | 253,78 | 314,61 | 0,0115 | 1,24 |
| *ENSRNOG00000066115* | 0,43 | 3,58 | 0,0117 | 8,28 |
| *Kcnrg* | 40,64 | 27,55 | 0,0117 | -1,48 |
| *Itga8* | 62,57 | 104,41 | 0,0118 | 1,67 |
| *Defb41* | 5,89 | 1,68 | 0,0118 | -3,52 |
| *ENSRNOG00000070938* | 165,42 | 224,93 | 0,0119 | 1,36 |
| *ENSRNOG00000070285* | 1205,89 | 1533,39 | 0,0124 | 1,27 |
| *Hist1h2ao* | 31,37 | 19,25 | 0,0125 | -1,63 |
| *C10h17orf67* | 7,54 | 2,59 | 0,0127 | -2,92 |
| *Pidd1* | 52,65 | 36,48 | 0,0129 | -1,44 |
| *Colq* | 77,40 | 52,39 | 0,0131 | -1,48 |
| *Selenow* | 228,11 | 363,57 | 0,0137 | 1,59 |
| *Bdkrb2* | 2,61 | 7,23 | 0,0139 | 2,77 |
| *Ibsp* | 5,91 | 1,82 | 0,0140 | -3,25 |
| *LOC108350764* | 22,65 | 13,69 | 0,0140 | -1,65 |
| *Zfyve19* | 160,60 | 203,88 | 0,0141 | 1,27 |
| *Clk3* | 725,04 | 854,79 | 0,0141 | 1,18 |
| *Gosr2* | 1178,65 | 1040,62 | 0,0144 | -1,13 |
| *Mmachc* | 207,23 | 162,71 | 0,0146 | -1,27 |
| *Prss53* | 72,74 | 104,75 | 0,0146 | 1,44 |
| *Gnas* | 186,33 | 270,23 | 0,0148 | 1,45 |
| *Panx3* | 1,40 | 5,59 | 0,0158 | 3,99 |
| *Irf4* | 15,44 | 24,15 | 0,0162 | 1,56 |
| *Gpr27* | 174,89 | 138,96 | 0,0163 | -1,26 |
| *ENSRNOG00000062328* | 10,96 | 23,17 | 0,0163 | 2,11 |
| *Pkm-ps20* | 789,49 | 645,01 | 0,0168 | -1,22 |
| *Cartpt* | 151,21 | 258,79 | 0,0170 | 1,71 |
| *Slc39a5* | 6,17 | 13,32 | 0,0171 | 2,16 |
| *AABR07044388.2* | 271,69 | 216,12 | 0,0171 | -1,26 |
| *ENSRNOG00000066808* | 15,16 | 24,85 | 0,0173 | 1,64 |
| *ENSRNOG00000070948* | 26,67 | 39,53 | 0,0173 | 1,48 |
| *ENSRNOG00000070541* | 24,89 | 35,89 | 0,0174 | 1,44 |
| *Milr1* | 1,73 | 7,47 | 0,0176 | 4,31 |
| *Epm2a* | 145,51 | 102,78 | 0,0177 | -1,42 |
| *Dux4* | 25,27 | 38,84 | 0,0179 | 1,54 |
| *ENSRNOG00000070442* | 56,10 | 26,03 | 0,0180 | -2,15 |
| *Pnpla1* | 279,23 | 216,35 | 0,0180 | -1,29 |
| *Ccnl1* | 314,03 | 375,98 | 0,0180 | 1,20 |
| *Mtrf1* | 193,60 | 244,92 | 0,0181 | 1,27 |
| *MGC114499* | 0,41 | 3,26 | 0,0188 | 8,03 |
| *Ap5z1* | 393,72 | 451,64 | 0,0188 | 1,15 |
| *Neurl4* | 1430,94 | 1253,73 | 0,0192 | -1,14 |
| *AABR07009978.1* | 15,67 | 26,68 | 0,0192 | 1,70 |
| *Crtc3* | 98,07 | 73,52 | 0,0195 | -1,33 |
| *Map3k15* | 27,27 | 41,89 | 0,0196 | 1,54 |
| *Mrtfa* | 544,37 | 473,53 | 0,0203 | -1,15 |
| *Entrep2* | 731,82 | 627,71 | 0,0203 | -1,17 |
| *Stk38* | 305,37 | 366,57 | 0,0204 | 1,20 |
| *Tcea1l1* | 121,43 | 194,90 | 0,0204 | 1,61 |
| *Recql4* | 19,97 | 31,52 | 0,0207 | 1,58 |
| *Lrpprc* | 1448,49 | 1276,42 | 0,0211 | -1,13 |
| *Cyp2d2* | 2,71 | 0,45 | 0,0213 | -6,00 |
| *Snora2l1* | 4,06 | 1,18 | 0,0219 | -3,45 |
| *Ube2o* | 2165,95 | 1947,03 | 0,0223 | -1,11 |
| *Prph2* | 0,40 | 2,91 | 0,0223 | 7,36 |
| *Hp* | 13,34 | 26,91 | 0,0223 | 2,02 |
| *Scrt1* | 608,44 | 464,50 | 0,0225 | -1,31 |
| *RGD1563835* | 55,70 | 87,41 | 0,0225 | 1,57 |
| *Haus7* | 153,61 | 193,21 | 0,0228 | 1,26 |
| *LOC689816* | 2,84 | 8,47 | 0,0228 | 2,98 |
| *ENSRNOG00000043210* | 226,68 | 293,06 | 0,0230 | 1,29 |
| *Dusp5* | 66,31 | 89,31 | 0,0230 | 1,35 |
| *Kif20b* | 38,38 | 26,64 | 0,0236 | -1,44 |
| *ENSRNOG00000069588* | 4100,97 | 5041,77 | 0,0240 | 1,23 |
| *Tdrd1* | 5,97 | 2,00 | 0,0241 | -2,98 |
| *LOC120102603* | 7,63 | 2,44 | 0,0242 | -3,12 |
| *Actg1* | 12188,97 | 10428,26 | 0,0245 | -1,17 |
| *Fgf5* | 4,73 | 1,47 | 0,0247 | -3,22 |
| *ENSRNOG00000069893* | 101,19 | 133,67 | 0,0249 | 1,32 |
| *Acr* | 1,42 | 4,61 | 0,0251 | 3,24 |
| *Sumo3* | 72,82 | 92,58 | 0,0252 | 1,27 |
| *Mir3064* | 67,97 | 92,49 | 0,0253 | 1,36 |
| *ENSRNOG00000068896* | 1,61 | 5,03 | 0,0254 | 3,12 |
| *Snord19* | 0,61 | 3,79 | 0,0254 | 6,22 |
| *Aurkb* | 36,98 | 20,93 | 0,0256 | -1,77 |
| *ENSRNOG00000063937* | 41,32 | 56,74 | 0,0257 | 1,37 |
| *ENSRNOG00000067662* | 5,34 | 10,85 | 0,0257 | 2,03 |
| *ENSRNOG00000071138* | 435,59 | 508,50 | 0,0261 | 1,17 |
| *Tmprss6* | 43,38 | 60,68 | 0,0263 | 1,40 |
| *Fcgr2b* | 68,13 | 94,35 | 0,0269 | 1,38 |
| *Lrriq1* | 38,20 | 21,94 | 0,0269 | -1,74 |
| *Crispld2* | 42,85 | 28,74 | 0,0271 | -1,49 |
| *Tbxa2r* | 26,65 | 17,73 | 0,0272 | -1,50 |
| *Tecpr1* | 1317,58 | 1108,59 | 0,0273 | -1,19 |
| *Myo9b* | 742,32 | 891,29 | 0,0274 | 1,20 |
| *Mir29b2* | 1,16 | 4,39 | 0,0276 | 3,78 |
| *ENSRNOG00000066571* | 6,94 | 2,52 | 0,0277 | -2,75 |
| *Slc22a14* | 1,58 | 5,13 | 0,0277 | 3,24 |
| *LOC120095871* | 144,99 | 181,20 | 0,0282 | 1,25 |
| *Ftsj3* | 406,29 | 456,14 | 0,0286 | 1,12 |
| *Cmtr1* | 1059,47 | 924,71 | 0,0288 | -1,15 |
| *ENSRNOG00000067467* | 76,12 | 54,29 | 0,0288 | -1,40 |
| *AABR07044383.1* | 65,90 | 47,38 | 0,0288 | -1,39 |
| *LOC680385* | 125,61 | 161,40 | 0,0289 | 1,28 |
| *Eepd1* | 384,95 | 332,65 | 0,0289 | -1,16 |
| *AABR07053065.1* | 3,44 | 0,74 | 0,0291 | -4,65 |
| *Mgat5b* | 1114,84 | 971,02 | 0,0296 | -1,15 |
| *Thada* | 233,10 | 197,20 | 0,0296 | -1,18 |
| *ENSRNOG00000065864* | 12,92 | 6,64 | 0,0298 | -1,94 |
| *Orai2* | 1118,27 | 977,62 | 0,0298 | -1,14 |
| *Anks6* | 75,02 | 96,25 | 0,0299 | 1,28 |
| *Tjap1* | 405,16 | 488,09 | 0,0299 | 1,20 |
| *Cd300le* | 7,18 | 2,90 | 0,0301 | -2,47 |
| *Perp* | 23,52 | 36,24 | 0,0306 | 1,54 |
| *Adcy4* | 34,81 | 47,73 | 0,0308 | 1,37 |
| *Prpf39* | 311,24 | 398,30 | 0,0309 | 1,28 |
| *Selplg* | 622,34 | 803,14 | 0,0310 | 1,29 |
| *Chp2* | 17,72 | 10,02 | 0,0310 | -1,77 |
| *Ceacam4* | 4,39 | 10,90 | 0,0312 | 2,48 |
| *Syngr1* | 748,37 | 570,07 | 0,0313 | -1,31 |
| *Aph1b* | 17,15 | 25,77 | 0,0313 | 1,50 |
| *Lrrcc1* | 103,75 | 126,18 | 0,0314 | 1,22 |
| *Hspa8-ps25* | 78,27 | 53,45 | 0,0320 | -1,46 |
| *Kcnb2* | 52,02 | 34,53 | 0,0321 | -1,51 |
| *Sap25* | 21,35 | 37,67 | 0,0321 | 1,76 |
| *Ppip5k2* | 337,43 | 429,01 | 0,0322 | 1,27 |
| *Tph1* | 4,21 | 11,68 | 0,0323 | 2,77 |
| *Isca2-ps1* | 36,31 | 54,92 | 0,0325 | 1,51 |
| *ENSRNOG00000063687* | 6,85 | 14,71 | 0,0325 | 2,15 |
| *LOC120095727* | 0,94 | 4,24 | 0,0326 | 4,50 |
| *Fam120b* | 1410,42 | 1272,27 | 0,0327 | -1,11 |
| *C8g* | 77,22 | 101,38 | 0,0328 | 1,31 |
| *ENSRNOG00000067425* | 2,63 | 0,44 | 0,0330 | -6,00 |
| *Tmem150a* | 119,83 | 145,24 | 0,0332 | 1,21 |
| *ENSRNOG00000065219* | 3,18 | 9,04 | 0,0336 | 2,85 |
| *Msantd5* | 18,46 | 27,26 | 0,0342 | 1,48 |
| *Gpr26* | 40,60 | 27,52 | 0,0347 | -1,47 |
| *Nostrin* | 52,18 | 72,06 | 0,0348 | 1,38 |
| *Tgfb1* | 128,36 | 156,66 | 0,0352 | 1,22 |
| *Frmpd2* | 31,51 | 18,44 | 0,0355 | -1,71 |
| *Rpl7-ps3* | 15,11 | 8,87 | 0,0355 | -1,70 |
| *Eppk1* | 4,66 | 1,77 | 0,0356 | -2,64 |
| *Nsmaf* | 224,16 | 177,14 | 0,0359 | -1,27 |
| *ENSRNOG00000066253* | 1,95 | 6,69 | 0,0362 | 3,44 |
| *LOC102556098* | 3,94 | 8,88 | 0,0362 | 2,25 |
| *Pigbos1* | 57,19 | 78,21 | 0,0367 | 1,37 |
| *ENSRNOG00000067458* | 161,82 | 195,18 | 0,0369 | 1,21 |
| *Slc26a6* | 78,53 | 103,21 | 0,0370 | 1,31 |
| *ENSRNOG00000067586* | 3,46 | 0,90 | 0,0371 | -3,82 |
| *LOC100910341* | 7,04 | 12,42 | 0,0371 | 1,77 |
| *Zmym6* | 294,33 | 343,99 | 0,0372 | 1,17 |
| *Maff* | 11,81 | 20,70 | 0,0375 | 1,75 |
| *Brinp1* | 2005,40 | 1740,19 | 0,0378 | -1,15 |
| *Atp2a3* | 61,56 | 47,23 | 0,0379 | -1,30 |
| *Zcchc7* | 247,50 | 325,23 | 0,0380 | 1,31 |
| *Acp3* | 4,94 | 1,16 | 0,0382 | -4,26 |
| *ENSRNOG00000068224* | 7,48 | 13,08 | 0,0382 | 1,75 |
| *Nup214* | 653,35 | 575,55 | 0,0382 | -1,14 |
| *Mir21* | 3,44 | 8,83 | 0,0382 | 2,57 |
| *Nr5a1* | 0,54 | 3,66 | 0,0384 | 6,78 |
| *Kdm4d* | 49,54 | 36,01 | 0,0384 | -1,38 |
| *Fbxo24* | 2,21 | 0,37 | 0,0386 | -6,00 |
| *Pcdh1* | 2467,76 | 2010,20 | 0,0389 | -1,23 |
| *Thsd1* | 147,17 | 173,28 | 0,0389 | 1,18 |
| *Calr3* | 45,69 | 60,50 | 0,0394 | 1,32 |
| *Tmem121b* | 141,80 | 97,94 | 0,0394 | -1,45 |
| *Dhx38* | 562,63 | 497,00 | 0,0394 | -1,13 |
| *Pnpla6* | 677,58 | 591,84 | 0,0397 | -1,14 |
| *AABR07072207.1* | 203,08 | 293,11 | 0,0398 | 1,44 |
| *Nmnat1* | 226,32 | 272,02 | 0,0399 | 1,20 |
| *Olig2* | 591,05 | 677,71 | 0,0403 | 1,15 |
| *Sbds* | 1100,80 | 1257,36 | 0,0403 | 1,14 |
| *Furin* | 588,47 | 513,23 | 0,0404 | -1,15 |
| *Mir135a* | 0,58 | 3,14 | 0,0405 | 5,37 |
| *Mir1249* | 1,55 | 5,87 | 0,0405 | 3,79 |
| *Cthrc1* | 9,04 | 4,73 | 0,0407 | -1,91 |
| *Hps3* | 212,25 | 177,50 | 0,0410 | -1,20 |
| *Trim68* | 221,89 | 185,72 | 0,0410 | -1,19 |
| *Tifab* | 152,22 | 206,05 | 0,0410 | 1,35 |
| *Rsrp1* | 3149,03 | 3693,05 | 0,0412 | 1,17 |
| *LOC100911253* | 25,39 | 55,26 | 0,0412 | 2,18 |
| *LOC102551788* | 13,70 | 6,94 | 0,0413 | -1,97 |
| *Farsb* | 626,19 | 544,91 | 0,0414 | -1,15 |
| *Acap1* | 0,18 | 2,19 | 0,0414 | 12,05 |
| *Gbp1* | 10,23 | 3,54 | 0,0414 | -2,89 |
| *Ccdc57* | 146,09 | 184,69 | 0,0415 | 1,26 |
| *LOC120102832* | 49,17 | 69,82 | 0,0416 | 1,42 |
| *ENSRNOG00000068587* | 7,49 | 15,67 | 0,0418 | 2,09 |
| *Retsat* | 1524,49 | 1197,82 | 0,0419 | -1,27 |
| *ENSRNOG00000064734* | 14,50 | 36,03 | 0,0422 | 2,49 |
| *Foxh1* | 22,18 | 34,04 | 0,0423 | 1,53 |
| *ENSRNOG00000064849* | 64,97 | 85,40 | 0,0423 | 1,31 |
| *Rasal3* | 25,38 | 36,00 | 0,0424 | 1,42 |
| *Ccdc62* | 77,94 | 99,10 | 0,0425 | 1,27 |
| *LOC120102634* | 0,61 | 3,65 | 0,0426 | 5,93 |
| *Zfp61* | 407,58 | 493,05 | 0,0428 | 1,21 |
| *ENSRNOG00000065144* | 16,69 | 28,16 | 0,0428 | 1,69 |
| *Tmem120b* | 155,98 | 122,28 | 0,0429 | -1,28 |
| *ENSRNOG00000063787* | 14,16 | 23,82 | 0,0430 | 1,68 |
| *Jph3* | 6378,94 | 5658,59 | 0,0435 | -1,13 |
| *Col7a1* | 53,51 | 74,70 | 0,0435 | 1,40 |
| *ENSRNOG00000065245* | 15,53 | 24,94 | 0,0436 | 1,61 |
| *ENSRNOG00000030963* | 3445,51 | 2563,94 | 0,0437 | -1,34 |
| *ENSRNOG00000063961* | 32,96 | 51,94 | 0,0437 | 1,58 |
| *Tra2a* | 629,10 | 740,19 | 0,0439 | 1,18 |
| *LOC102549716* | 1,75 | 5,20 | 0,0441 | 2,96 |
| *LOC120098677* | 2,18 | 5,54 | 0,0442 | 2,54 |
| *ENSRNOG00000067218* | 5,02 | 9,44 | 0,0443 | 1,88 |
| *Ankrd1* | 7,56 | 13,65 | 0,0444 | 1,81 |
| *Gas8* | 100,73 | 125,03 | 0,0448 | 1,24 |
| *Ryr3* | 545,11 | 740,71 | 0,0448 | 1,36 |
| *Scarf1* | 52,43 | 67,75 | 0,0449 | 1,29 |
| *Pmch* | 2,89 | 8,11 | 0,0450 | 2,80 |
| *Tppp2* | 9,23 | 4,69 | 0,0452 | -1,97 |
| *ENSRNOG00000062892* | 0,92 | 5,16 | 0,0454 | 5,58 |
| *Ssh3* | 196,67 | 238,35 | 0,0455 | 1,21 |
| *C1s* | 97,80 | 74,62 | 0,0456 | -1,31 |
| *ENSRNOG00000069214* | 27,97 | 18,64 | 0,0457 | -1,50 |
| *Scmh1* | 708,13 | 636,21 | 0,0457 | -1,11 |
| *Ltk* | 281,57 | 346,43 | 0,0460 | 1,23 |
| *RGD1565297* | 7,58 | 14,10 | 0,0463 | 1,86 |
| *Wtip* | 55,50 | 72,56 | 0,0464 | 1,31 |
| *Eya3* | 426,18 | 350,05 | 0,0465 | -1,22 |
| *Pfn4* | 1,75 | 4,99 | 0,0467 | 2,85 |
| *P3h1* | 96,26 | 117,28 | 0,0467 | 1,22 |
| *Tcim* | 17,37 | 27,10 | 0,0469 | 1,56 |
| *Lat* | 7,07 | 14,42 | 0,0471 | 2,04 |
| *ENSRNOG00000069867* | 10,13 | 16,65 | 0,0471 | 1,64 |
| *Dnah1* | 53,06 | 27,31 | 0,0471 | -1,94 |
| *Dglucy* | 98,63 | 79,98 | 0,0472 | -1,23 |
| *ENSRNOG00000063355* | 3,57 | 0,92 | 0,0473 | -3,87 |
| *Albfm1* | 15,70 | 28,96 | 0,0475 | 1,84 |
| *Prdm11* | 49,35 | 30,46 | 0,0475 | -1,62 |
| *Pp2d1* | 0,43 | 2,94 | 0,0479 | 6,78 |
| *Clk4* | 374,23 | 445,26 | 0,0480 | 1,19 |
| *Slc16a4* | 41,00 | 57,29 | 0,0481 | 1,40 |
| *Mr1* | 61,96 | 42,55 | 0,0483 | -1,46 |
| *ENSRNOG00000068841* | 6,27 | 2,38 | 0,0484 | -2,64 |
| *Mettl3* | 440,02 | 520,77 | 0,0485 | 1,18 |
| *Bbs5* | 176,19 | 203,44 | 0,0485 | 1,15 |
| *Slc26a10* | 268,31 | 328,03 | 0,0486 | 1,22 |
| *Snord56* | 2,32 | 6,21 | 0,0486 | 2,68 |
| *Ccdc9b* | 56,02 | 40,18 | 0,0488 | -1,39 |
| *ENSRNOG00000070157* | 0,10 | 1,66 | 0,0488 | 16,60 |
| *Stard5* | 97,17 | 77,98 | 0,0490 | -1,25 |
| *Smim19* | 647,70 | 725,59 | 0,0493 | 1,12 |
| *Akap8l* | 1414,99 | 1707,78 | 0,0495 | 1,21 |
| *Ier3ip1-ps1* | 2,00 | 0,33 | 0,0495 | -6,00 |
| *Olr1* | 19,13 | 11,79 | 0,0498 | -1,62 |
| *Sdf2l1* | 133,58 | 168,85 | 0,0499 | 1,26 |
| *ENSRNOG00000062644* | 552,11 | 766,37 | 0,0499 | 1,39 |

**HE-GFP vs HE-GDNF**

| **Gene symbol** | **Average norm read counts HE-GFP** | **Average norm read counts HE-GDNF** | **P-value** | **Fold Change** |
| --- | --- | --- | --- | --- |
| *Sel1l3* | 353,04 | 259,83 | 8,57E-06 | -1,36 |
| *Tshz3* | 225,70 | 141,67 | 9,40E-06 | -1,59 |
| *Ssbp3* | 1261,23 | 984,73 | 1,10E-05 | -1,28 |
| *Apoa5* | 19,89 | 6,77 | 1,51E-05 | -2,94 |
| *Cybb* | 5,74 | 25,73 | 1,68E-05 | 4,48 |
| *Cadm3* | 4815,41 | 3838,10 | 1,82E-05 | -1,25 |
| *ENSRNOG00000056855* | 49,33 | 88,15 | 2,19E-05 | 1,79 |
| *Myadm* | 693,96 | 538,61 | 4,69E-05 | -1,29 |
| *Cyp26b1* | 173,12 | 103,85 | 5,23E-05 | -1,67 |
| *Gng2* | 4113,07 | 2975,86 | 6,41E-05 | -1,38 |
| *Sh3gl2* | 1567,64 | 1117,36 | 6,74E-05 | -1,40 |
| *Plaur* | 28,70 | 14,37 | 8,25E-05 | -2,00 |
| *Rtn1* | 18432,86 | 16108,04 | 1,04E-04 | -1,14 |
| *Tmem108* | 218,52 | 153,66 | 1,16E-04 | -1,42 |
| *Prss12* | 278,47 | 175,87 | 1,18E-04 | -1,58 |
| *Hspa1a* | 43,86 | 113,78 | 1,29E-04 | 2,59 |
| *Gabra3* | 496,94 | 338,94 | 1,35E-04 | -1,47 |
| *Pcsk1* | 256,02 | 181,68 | 1,43E-04 | -1,41 |
| *Vstm2b* | 536,03 | 432,08 | 1,59E-04 | -1,24 |
| *Cbln1* | 258,86 | 179,04 | 1,88E-04 | -1,45 |
| *Dkk3* | 3276,71 | 2375,85 | 2,58E-04 | -1,38 |
| *Faxdc2* | 132,47 | 189,79 | 2,73E-04 | 1,43 |
| *Lypd6* | 105,77 | 70,65 | 2,79E-04 | -1,50 |
| *Syt17* | 247,71 | 147,80 | 3,48E-04 | -1,68 |
| *Ggta1* | 86,19 | 134,62 | 3,76E-04 | 1,56 |
| *Tnfrsf21* | 758,54 | 628,63 | 4,07E-04 | -1,21 |
| *Exph5* | 59,12 | 31,54 | 4,34E-04 | -1,87 |
| *Smap2* | 3564,92 | 3089,47 | 4,57E-04 | -1,15 |
| *Asap2* | 571,29 | 434,89 | 4,58E-04 | -1,31 |
| *Sync* | 100,47 | 61,19 | 4,73E-04 | -1,64 |
| *Cx3cl1* | 12522,25 | 11039,78 | 4,92E-04 | -1,13 |
| *Plcxd2* | 508,20 | 349,28 | 5,16E-04 | -1,45 |
| *Tp53i11* | 836,39 | 598,66 | 5,32E-04 | -1,40 |
| *Ppif* | 212,06 | 271,28 | 5,35E-04 | 1,28 |
| *Trim9* | 1902,60 | 1525,09 | 5,35E-04 | -1,25 |
| *Pwp1* | 198,68 | 258,25 | 5,83E-04 | 1,30 |
| *Gfra1* | 248,10 | 171,73 | 5,84E-04 | -1,44 |
| *Tvp23a* | 515,58 | 624,55 | 5,91E-04 | 1,21 |
| *Csnk1e* | 1161,18 | 983,99 | 6,36E-04 | -1,18 |
| *Necap1* | 2860,21 | 2521,94 | 6,57E-04 | -1,13 |
| *RT1-M1-2* | 6,97 | 1,32 | 6,58E-04 | -5,29 |
| *LOC120097345* | 369,03 | 510,73 | 6,71E-04 | 1,38 |
| *RGD1561661* | 10,55 | 3,17 | 6,99E-04 | -3,33 |
| *ENSRNOG00000068601* | 97,87 | 136,98 | 7,00E-04 | 1,40 |
| *Cbln2* | 313,34 | 174,08 | 7,51E-04 | -1,80 |
| *Faap20* | 155,29 | 210,54 | 7,54E-04 | 1,36 |
| *Khsrp* | 1074,10 | 911,50 | 7,62E-04 | -1,18 |
| *Bmp3* | 46,34 | 21,05 | 7,80E-04 | -2,20 |
| *Neurod2* | 360,15 | 203,65 | 8,20E-04 | -1,77 |
| *Nptxr* | 6883,51 | 4669,45 | 8,82E-04 | -1,47 |
| *Gdi1* | 10360,26 | 9104,53 | 9,08E-04 | -1,14 |
| *Nptx2* | 581,18 | 402,02 | 9,58E-04 | -1,45 |
| *Basp1* | 6009,95 | 4411,73 | 9,77E-04 | -1,36 |
| *Pigc* | 243,11 | 304,00 | 0,0010 | 1,25 |
| *Pitpna* | 3427,97 | 2985,81 | 0,0010 | -1,15 |
| *Aif1* | 279,37 | 363,75 | 0,0011 | 1,30 |
| *Tk2* | 336,66 | 434,27 | 0,0011 | 1,29 |
| *Grm2* | 35,03 | 18,72 | 0,0011 | -1,87 |
| *Rasgrf1* | 4662,79 | 3992,84 | 0,0011 | -1,17 |
| *Tnc* | 226,08 | 154,24 | 0,0011 | -1,47 |
| *Zfp775* | 156,40 | 203,57 | 0,0012 | 1,30 |
| *Prxl2c* | 362,25 | 431,93 | 0,0012 | 1,19 |
| *Fap* | 27,94 | 15,44 | 0,0012 | -1,81 |
| *Slit1* | 555,64 | 367,73 | 0,0012 | -1,51 |
| *Agmat* | 46,76 | 70,02 | 0,0013 | 1,50 |
| *Gabbr2* | 2444,53 | 1802,15 | 0,0013 | -1,36 |
| *Arhgdib* | 290,72 | 368,43 | 0,0013 | 1,27 |
| *Rtn4ip1* | 289,94 | 381,97 | 0,0014 | 1,32 |
| *Tmem200a* | 83,91 | 50,95 | 0,0014 | -1,65 |
| *Gna14* | 27,30 | 14,26 | 0,0014 | -1,91 |
| *Snap91* | 2765,34 | 2161,58 | 0,0015 | -1,28 |
| *Bdnf* | 207,75 | 142,68 | 0,0016 | -1,46 |
| *ENSRNOG00000067311* | 32,28 | 18,22 | 0,0016 | -1,77 |
| *Tsnaxip1* | 5,99 | 15,72 | 0,0017 | 2,63 |
| *Srek1ip1-ps1* | 99,16 | 135,18 | 0,0017 | 1,36 |
| *Hivep1* | 294,63 | 233,42 | 0,0017 | -1,26 |
| *Chrm3* | 544,19 | 399,91 | 0,0017 | -1,36 |
| *R3hdm1* | 1577,83 | 1318,49 | 0,0018 | -1,20 |
| *C1ql3* | 1278,64 | 836,52 | 0,0019 | -1,53 |
| *Mapk10* | 1930,19 | 1574,88 | 0,0019 | -1,23 |
| *Pgm2l1* | 2058,81 | 1640,58 | 0,0019 | -1,25 |
| *Ackr1* | 120,10 | 89,33 | 0,0020 | -1,34 |
| *Relt* | 67,89 | 94,99 | 0,0021 | 1,40 |
| *Fcsk* | 143,72 | 179,20 | 0,0021 | 1,25 |
| *Cdk14* | 1410,81 | 1110,60 | 0,0021 | -1,27 |
| *Yipf4* | 838,66 | 960,66 | 0,0021 | 1,15 |
| *Prrt1* | 2628,23 | 2256,94 | 0,0022 | -1,16 |
| *Mmp16* | 268,77 | 212,22 | 0,0022 | -1,27 |
| *Gpsm1* | 1266,57 | 1486,18 | 0,0023 | 1,17 |
| *Bhlhe22* | 108,46 | 56,45 | 0,0023 | -1,92 |
| *Pcdhb3* | 121,59 | 89,45 | 0,0023 | -1,36 |
| *Trabd2b* | 51,27 | 26,16 | 0,0024 | -1,96 |
| *Albfm1* | 28,78 | 14,16 | 0,0024 | -2,03 |
| *Mccc2* | 557,88 | 670,88 | 0,0024 | 1,20 |
| *R3hdm4* | 2379,20 | 2073,83 | 0,0025 | -1,15 |
| *Syngr1* | 811,14 | 638,72 | 0,0026 | -1,27 |
| *Pla2g2d* | 16,52 | 6,95 | 0,0026 | -2,38 |
| *Etv5* | 1018,58 | 804,02 | 0,0027 | -1,27 |
| *Ddah1* | 712,18 | 566,12 | 0,0027 | -1,26 |
| *Gabra5* | 692,44 | 506,18 | 0,0028 | -1,37 |
| *Ifi44l* | 6,76 | 16,03 | 0,0028 | 2,37 |
| *Scube1* | 370,05 | 250,04 | 0,0028 | -1,48 |
| *ENSRNOG00000062478* | 39,93 | 16,74 | 0,0029 | -2,38 |
| *Septin3* | 5383,25 | 4742,01 | 0,0029 | -1,14 |
| *Hs3st2* | 445,83 | 285,17 | 0,0029 | -1,56 |
| *Arhgef28* | 430,99 | 352,32 | 0,0029 | -1,22 |
| *Srebf2* | 2929,61 | 2454,17 | 0,0030 | -1,19 |
| *ENSRNOG00000063824* | 151,79 | 105,79 | 0,0030 | -1,43 |
| *LOC120099454* | 13,95 | 29,60 | 0,0030 | 2,12 |
| *Dpysl3* | 265,23 | 183,16 | 0,0031 | -1,45 |
| *Fezf2* | 202,56 | 141,59 | 0,0031 | -1,43 |
| *Lingo1* | 2866,03 | 2098,49 | 0,0031 | -1,37 |
| *Dpp10* | 688,98 | 526,57 | 0,0031 | -1,31 |
| *Zfp367* | 241,27 | 197,02 | 0,0032 | -1,22 |
| *Ntng2* | 427,12 | 282,45 | 0,0033 | -1,51 |
| *LOC120093363* | 0,10 | 7,74 | 0,0033 | 77,37 |
| *Fxyd6* | 1232,33 | 931,02 | 0,0033 | -1,32 |
| *Plxna1* | 503,57 | 363,29 | 0,0034 | -1,39 |
| *LOC498154* | 229,46 | 291,92 | 0,0035 | 1,27 |
| *Lhfpl3* | 234,58 | 186,39 | 0,0035 | -1,26 |
| *Ywhah* | 12607,94 | 10570,86 | 0,0037 | -1,19 |
| *Smim11* | 147,08 | 190,50 | 0,0037 | 1,30 |
| *Kcnc4* | 502,14 | 352,23 | 0,0037 | -1,43 |
| *Ret* | 57,78 | 38,93 | 0,0038 | -1,48 |
| *Mfsd2a* | 287,54 | 352,14 | 0,0038 | 1,22 |
| *Psmb10* | 242,32 | 316,93 | 0,0039 | 1,31 |
| *Gpr12* | 518,71 | 418,73 | 0,0039 | -1,24 |
| *Gna15* | 55,22 | 78,19 | 0,0039 | 1,42 |
| *Medag* | 140,89 | 95,62 | 0,0039 | -1,47 |
| *Cckbr* | 334,96 | 274,01 | 0,0040 | -1,22 |
| *Pcdhb8* | 114,71 | 81,29 | 0,0040 | -1,41 |
| *Acyp2* | 448,55 | 548,01 | 0,0040 | 1,22 |
| *Gspt1* | 1242,41 | 1115,78 | 0,0041 | -1,11 |
| *B3gat1* | 1416,67 | 1177,68 | 0,0042 | -1,20 |
| *B3gnt5* | 7,17 | 2,37 | 0,0042 | -3,02 |
| *ENSRNOG00000067620* | 180,45 | 225,97 | 0,0043 | 1,25 |
| *Satb2* | 261,37 | 153,65 | 0,0043 | -1,70 |
| *Rspo3* | 108,04 | 67,60 | 0,0044 | -1,60 |
| *Gins3* | 68,78 | 94,15 | 0,0045 | 1,37 |
| *ENSRNOG00000066171* | 54,85 | 75,02 | 0,0045 | 1,37 |
| *Cadm2* | 967,22 | 791,10 | 0,0046 | -1,22 |
| *Smyd1* | 84,77 | 49,25 | 0,0046 | -1,72 |
| *Pcdhb7* | 81,98 | 60,25 | 0,0046 | -1,36 |
| *Cyp11b1* | 43,08 | 19,02 | 0,0046 | -2,26 |
| *RT1-CE16* | 449,48 | 685,06 | 0,0047 | 1,52 |
| *Rnasel* | 68,78 | 48,56 | 0,0047 | -1,42 |
| *Kcnh7* | 71,60 | 44,45 | 0,0048 | -1,61 |
| *Syn1* | 6540,28 | 5446,10 | 0,0049 | -1,20 |
| *Pip5kl1* | 5,57 | 1,56 | 0,0049 | -3,56 |
| *Nppc* | 56,94 | 36,65 | 0,0049 | -1,55 |
| *Mas1* | 65,48 | 39,10 | 0,0049 | -1,67 |
| *Rims1* | 1952,30 | 1589,43 | 0,0050 | -1,23 |
| *Atp2b4* | 3037,74 | 2081,32 | 0,0050 | -1,46 |
| *Nppa* | 136,33 | 85,16 | 0,0051 | -1,60 |
| *Ppm1e* | 363,11 | 247,12 | 0,0051 | -1,47 |
| *Gkn2* | 6,12 | 1,49 | 0,0052 | -4,12 |
| *Rtn4rl2* | 232,35 | 135,31 | 0,0053 | -1,72 |
| *Slc4a7* | 77,49 | 46,80 | 0,0054 | -1,66 |
| *Adamts2* | 38,81 | 24,05 | 0,0054 | -1,61 |
| *Toe1* | 112,28 | 143,18 | 0,0054 | 1,28 |
| *Efhd2* | 2526,42 | 1835,35 | 0,0055 | -1,38 |
| *Srcin1* | 2755,90 | 2250,67 | 0,0055 | -1,22 |
| *Isy1* | 223,85 | 268,43 | 0,0056 | 1,20 |
| *Nr4a2* | 1163,99 | 844,60 | 0,0056 | -1,38 |
| *Ap3d1* | 3509,35 | 3151,96 | 0,0056 | -1,11 |
| *Fbxw7* | 948,70 | 813,57 | 0,0056 | -1,17 |
| *Ywhag* | 12565,17 | 10485,04 | 0,0058 | -1,20 |
| *Bves* | 10,16 | 3,68 | 0,0058 | -2,76 |
| *Dlgap1* | 3050,82 | 2435,65 | 0,0058 | -1,25 |
| *Slc7a11* | 266,60 | 174,32 | 0,0058 | -1,53 |
| *Calb1* | 1813,48 | 2200,78 | 0,0060 | 1,21 |
| *Acaa2* | 272,67 | 395,37 | 0,0061 | 1,45 |
| *Ism2* | 29,27 | 45,41 | 0,0061 | 1,55 |
| *Slc39a12* | 48,06 | 144,22 | 0,0061 | 3,00 |
| *Nploc4* | 1061,33 | 935,10 | 0,0061 | -1,13 |
| *Irf2bpl* | 1001,04 | 869,56 | 0,0062 | -1,15 |
| *Zdhhc14* | 1115,04 | 949,95 | 0,0063 | -1,17 |
| *Sptbn5* | 64,05 | 89,12 | 0,0063 | 1,39 |
| *Stxbp1* | 9502,70 | 8055,02 | 0,0063 | -1,18 |
| *Hs2st1* | 1069,74 | 953,63 | 0,0063 | -1,12 |
| *Pex7* | 81,78 | 110,78 | 0,0064 | 1,35 |
| *Vwa5b1* | 12,18 | 24,13 | 0,0064 | 1,98 |
| *Elavl4* | 794,52 | 606,75 | 0,0065 | -1,31 |
| *Grk4* | 62,43 | 86,83 | 0,0065 | 1,39 |
| *Sncb* | 5497,04 | 4282,33 | 0,0066 | -1,28 |
| *Hmgcs1* | 4480,84 | 3942,37 | 0,0066 | -1,14 |
| *ENSRNOG00000070376* | 57,37 | 37,97 | 0,0066 | -1,51 |
| *Myo18b* | 5,19 | 12,66 | 0,0067 | 2,44 |
| *Epha5* | 651,59 | 461,05 | 0,0067 | -1,41 |
| *Kcnk9* | 89,81 | 55,09 | 0,0067 | -1,63 |
| *Krcc1* | 290,01 | 359,36 | 0,0068 | 1,24 |
| *Fam83g* | 6,55 | 2,22 | 0,0068 | -2,95 |
| *Lrfn2* | 193,85 | 126,82 | 0,0068 | -1,53 |
| *Kcnt2* | 125,82 | 79,40 | 0,0069 | -1,58 |
| *Uck2* | 181,61 | 143,60 | 0,0069 | -1,26 |
| *Farp1* | 419,96 | 340,16 | 0,0070 | -1,23 |
| *Acad8* | 350,66 | 422,83 | 0,0070 | 1,21 |
| *Gdap1* | 1070,65 | 895,69 | 0,0070 | -1,20 |
| *LOC120102832* | 46,45 | 74,09 | 0,0072 | 1,60 |
| *Stx1a* | 1555,55 | 1093,39 | 0,0072 | -1,42 |
| *Sdk2* | 264,98 | 203,12 | 0,0073 | -1,30 |
| *Lrfn1* | 388,93 | 294,94 | 0,0073 | -1,32 |
| *Chmp4c* | 1,77 | 7,09 | 0,0073 | 4,01 |
| *Wasf1* | 4312,24 | 3867,36 | 0,0075 | -1,12 |
| *Nfic* | 736,07 | 643,84 | 0,0075 | -1,14 |
| *Hdac11* | 4835,35 | 5381,09 | 0,0076 | 1,11 |
| *Rab6a* | 7761,53 | 6913,50 | 0,0076 | -1,12 |
| *Ndufaf6* | 123,63 | 165,18 | 0,0077 | 1,34 |
| *Mapk8* | 1205,30 | 1033,96 | 0,0077 | -1,17 |
| *Tef* | 5018,19 | 5841,92 | 0,0078 | 1,16 |
| *Plekhg5* | 523,41 | 421,02 | 0,0079 | -1,24 |
| *Ptprcap* | 14,09 | 25,44 | 0,0081 | 1,81 |
| *Ctso* | 329,49 | 390,61 | 0,0082 | 1,19 |
| *Zdhhc22* | 111,34 | 73,97 | 0,0082 | -1,51 |
| *Islr2* | 926,49 | 670,65 | 0,0082 | -1,38 |
| *AABR07027854.1* | 129,30 | 96,22 | 0,0082 | -1,34 |
| *Slc16a4* | 34,62 | 52,09 | 0,0084 | 1,50 |
| *Serpini1* | 1976,98 | 1462,04 | 0,0084 | -1,35 |
| *Kif24* | 57,41 | 38,91 | 0,0086 | -1,48 |
| *Cdk18* | 302,43 | 243,10 | 0,0086 | -1,24 |
| *Rap1gds1* | 2691,16 | 2242,28 | 0,0087 | -1,20 |
| *Gale* | 156,42 | 124,96 | 0,0088 | -1,25 |
| *Ncdn* | 15960,32 | 17576,26 | 0,0089 | 1,10 |
| *ENSRNOG00000069845* | 21,29 | 32,99 | 0,0090 | 1,55 |
| *Wdr76* | 70,17 | 50,54 | 0,0091 | -1,39 |
| *Atp6v0a1* | 5739,69 | 5035,61 | 0,0094 | -1,14 |
| *Phospho2* | 557,07 | 649,12 | 0,0094 | 1,17 |
| *ENSRNOG00000066989* | 28,52 | 16,49 | 0,0094 | -1,73 |
| *RT1-N2* | 18,21 | 4,85 | 0,0095 | -3,75 |
| *ENSRNOG00000068880* | 46,26 | 79,45 | 0,0095 | 1,72 |
| *P2ry1* | 167,04 | 204,87 | 0,0095 | 1,23 |
| *Gnao1* | 9491,18 | 8430,60 | 0,0096 | -1,13 |
| *AABR07000989.1* | 4,17 | 0,86 | 0,0096 | -4,84 |
| *Olfm1* | 7581,24 | 5486,15 | 0,0096 | -1,38 |
| *Incenp* | 161,82 | 127,60 | 0,0096 | -1,27 |
| *Khdrbs3* | 2024,31 | 1635,55 | 0,0097 | -1,24 |
| *Ptpn3* | 181,97 | 113,55 | 0,0097 | -1,60 |
| *Igfbpl1* | 46,40 | 65,75 | 0,0097 | 1,42 |
| *LOC100910996* | 442,45 | 339,67 | 0,0098 | -1,30 |
| *Asic4* | 2309,27 | 2783,51 | 0,0099 | 1,21 |
| *Tbr1* | 1252,51 | 734,26 | 0,0099 | -1,71 |
| *Atp6v1c1* | 2975,89 | 2674,48 | 0,0099 | -1,11 |
| *Mdp1* | 285,08 | 344,10 | 0,0100 | 1,21 |
| *Homer2* | 93,51 | 70,47 | 0,0100 | -1,33 |
| *Gpr34* | 399,85 | 479,82 | 0,0101 | 1,20 |
| *Cep170b* | 3014,11 | 2484,79 | 0,0101 | -1,21 |
| *Slc35c2* | 447,54 | 533,60 | 0,0101 | 1,19 |
| *Nek3* | 92,06 | 70,42 | 0,0101 | -1,31 |
| *Hk1* | 5918,96 | 5210,89 | 0,0103 | -1,14 |
| *Sema3e* | 353,06 | 268,98 | 0,0103 | -1,31 |
| *ENSRNOG00000064457* | 211,75 | 159,68 | 0,0103 | -1,33 |
| *ENSRNOG00000068410* | 76,42 | 45,48 | 0,0104 | -1,68 |
| *Tmem150c* | 348,24 | 279,80 | 0,0104 | -1,24 |
| *ENSRNOG00000069152* | 13,98 | 24,14 | 0,0104 | 1,73 |
| *Rapgefl1* | 1829,05 | 1557,98 | 0,0104 | -1,17 |
| *Trim68* | 216,30 | 168,72 | 0,0105 | -1,28 |
| *Trh* | 71,64 | 98,06 | 0,0105 | 1,37 |
| *Dlx2* | 152,32 | 104,65 | 0,0106 | -1,46 |
| *Rbm43* | 33,08 | 20,79 | 0,0106 | -1,59 |
| *Tcerg1l* | 95,92 | 71,31 | 0,0107 | -1,35 |
| *ENSRNOG00000068741* | 4041,00 | 5052,62 | 0,0107 | 1,25 |
| *ENSRNOG00000062722* | 0,20 | 4,16 | 0,0107 | 20,76 |
| *Ecm2* | 36,22 | 51,75 | 0,0107 | 1,43 |
| *Slc29a3* | 335,88 | 393,36 | 0,0108 | 1,17 |
| *Vgf* | 3551,46 | 2900,22 | 0,0108 | -1,22 |
| *Nuak1* | 670,62 | 537,76 | 0,0108 | -1,25 |
| *Kcnk12* | 59,02 | 34,07 | 0,0109 | -1,73 |
| *Gpd1l* | 1329,69 | 1193,05 | 0,0109 | -1,11 |
| *Spint2* | 285,13 | 359,00 | 0,0110 | 1,26 |
| *Clec2d* | 4,58 | 1,10 | 0,0110 | -4,18 |
| *Fgf12* | 758,40 | 653,78 | 0,0110 | -1,16 |
| *Pxk* | 526,86 | 442,04 | 0,0110 | -1,19 |
| *Slc25a22* | 2565,22 | 2201,72 | 0,0110 | -1,17 |
| *ENSRNOG00000071119* | 4071,99 | 3548,79 | 0,0111 | -1,15 |
| *Glra2* | 290,26 | 238,84 | 0,0112 | -1,22 |
| *Lap3* | 1345,91 | 1554,41 | 0,0112 | 1,15 |
| *Arhgef15* | 108,79 | 135,00 | 0,0112 | 1,24 |
| *Tle3* | 526,33 | 449,60 | 0,0112 | -1,17 |
| *Castor2* | 1589,67 | 1418,85 | 0,0112 | -1,12 |
| *Chrna7* | 59,29 | 34,75 | 0,0112 | -1,71 |
| *Sdk1* | 152,44 | 117,90 | 0,0113 | -1,29 |
| *Adra1d* | 92,32 | 56,67 | 0,0113 | -1,63 |
| *Bbc3* | 102,47 | 71,96 | 0,0114 | -1,42 |
| *Cdh24* | 190,68 | 138,03 | 0,0114 | -1,38 |
| *Sgsm1* | 1112,38 | 934,75 | 0,0115 | -1,19 |
| *Canx* | 8189,61 | 7401,58 | 0,0115 | -1,11 |
| *Ccdc34* | 162,49 | 206,23 | 0,0115 | 1,27 |
| *Sap18* | 749,03 | 841,80 | 0,0115 | 1,12 |
| *Rassf5* | 298,60 | 228,85 | 0,0115 | -1,30 |
| *Kcnj6* | 466,16 | 307,27 | 0,0116 | -1,52 |
| *Nxph3* | 966,90 | 671,25 | 0,0116 | -1,44 |
| *Septin9* | 1101,20 | 884,99 | 0,0116 | -1,24 |
| *Mal2* | 1771,21 | 1533,00 | 0,0116 | -1,16 |
| *RGD1561149* | 150,48 | 107,54 | 0,0117 | -1,40 |
| *Sstr2* | 311,38 | 235,17 | 0,0118 | -1,32 |
| *Otx1* | 86,74 | 61,67 | 0,0118 | -1,41 |
| *Hivep2* | 2437,25 | 2106,28 | 0,0118 | -1,16 |
| *Apbb2* | 969,02 | 843,79 | 0,0118 | -1,15 |
| *Rnf38* | 159,27 | 111,22 | 0,0118 | -1,43 |
| *Dlgap4* | 2482,90 | 2217,76 | 0,0119 | -1,12 |
| *Alas2* | 7,52 | 1,73 | 0,0120 | -4,35 |
| *Rpa3* | 148,50 | 179,77 | 0,0120 | 1,21 |
| *Dclk1* | 11348,57 | 10010,06 | 0,0122 | -1,13 |
| *Ddx11* | 53,19 | 39,14 | 0,0122 | -1,36 |
| *Ptx3* | 13,75 | 6,75 | 0,0122 | -2,04 |
| *Ddit4l* | 18,87 | 9,71 | 0,0123 | -1,94 |
| *Fam81a* | 1043,27 | 766,84 | 0,0123 | -1,36 |
| *Krt8* | 16,29 | 29,04 | 0,0123 | 1,78 |
| *Serp1* | 1190,60 | 1353,20 | 0,0124 | 1,14 |
| *Abca7* | 208,05 | 168,70 | 0,0125 | -1,23 |
| *Kcne4* | 20,82 | 36,93 | 0,0126 | 1,77 |
| *Cnih3* | 287,50 | 213,10 | 0,0127 | -1,35 |
| *Ube2j1* | 2057,04 | 1883,93 | 0,0127 | -1,09 |
| *Nxpe1* | 24,98 | 40,20 | 0,0128 | 1,61 |
| *Tmem35a* | 960,72 | 804,61 | 0,0129 | -1,19 |
| *Csmd2* | 466,03 | 369,98 | 0,0129 | -1,26 |
| *Htr2a* | 279,76 | 226,64 | 0,0129 | -1,23 |
| *Lcorl* | 109,47 | 75,90 | 0,0129 | -1,44 |
| *Rhof* | 306,55 | 232,62 | 0,0130 | -1,32 |
| *LOC102554670* | 16,49 | 8,29 | 0,0130 | -1,99 |
| *Itpa* | 1010,54 | 1160,64 | 0,0130 | 1,15 |
| *Ccdc88b* | 85,67 | 107,39 | 0,0131 | 1,25 |
| *Mki67* | 73,51 | 47,16 | 0,0131 | -1,56 |
| *Gtf2h2* | 101,61 | 126,41 | 0,0132 | 1,24 |
| *Gng7* | 19148,46 | 24292,91 | 0,0133 | 1,27 |
| *Naf1* | 206,71 | 174,06 | 0,0133 | -1,19 |
| *Sema3c* | 334,82 | 254,15 | 0,0134 | -1,32 |
| *Crispld1* | 11,82 | 22,05 | 0,0136 | 1,87 |
| *ENSRNOG00000070783* | 54,92 | 79,76 | 0,0136 | 1,45 |
| *Itga2b* | 93,58 | 116,00 | 0,0136 | 1,24 |
| *Itgb1bp1* | 609,42 | 724,67 | 0,0136 | 1,19 |
| *Ptpru* | 336,73 | 244,47 | 0,0137 | -1,38 |
| *Lin7c* | 1352,87 | 1538,77 | 0,0138 | 1,14 |
| *Atoh7* | 20,37 | 9,50 | 0,0139 | -2,14 |
| *Nrg3* | 441,20 | 381,90 | 0,0141 | -1,16 |
| *Stau2* | 2986,19 | 2734,06 | 0,0142 | -1,09 |
| *Slc25a25* | 1366,07 | 1785,06 | 0,0142 | 1,31 |
| *ENSRNOG00000068460* | 136,95 | 108,52 | 0,0143 | -1,26 |
| *Gnai1* | 1610,35 | 1315,84 | 0,0144 | -1,22 |
| *Cadps* | 2117,34 | 1825,44 | 0,0145 | -1,16 |
| *Kctd18* | 191,29 | 232,65 | 0,0146 | 1,22 |
| *Otud1* | 383,78 | 320,13 | 0,0147 | -1,20 |
| *Gal* | 39,48 | 22,27 | 0,0148 | -1,77 |
| *ENSRNOG00000070008* | 6,58 | 2,43 | 0,0151 | -2,70 |
| *Angptl4* | 90,54 | 116,96 | 0,0151 | 1,29 |
| *Rad23b* | 3443,57 | 3116,56 | 0,0152 | -1,10 |
| *Rspo2* | 145,45 | 99,49 | 0,0152 | -1,46 |
| *Sestd1* | 204,26 | 166,85 | 0,0152 | -1,22 |
| *Nptx1* | 3290,91 | 2119,62 | 0,0153 | -1,55 |
| *Slc29a1* | 190,14 | 235,35 | 0,0153 | 1,24 |
| *Gng11* | 75,24 | 103,29 | 0,0154 | 1,37 |
| *Tmem221* | 1,22 | 5,36 | 0,0154 | 4,40 |
| *Kif28p* | 26,58 | 41,10 | 0,0154 | 1,55 |
| *Rad9a* | 333,08 | 405,43 | 0,0155 | 1,22 |
| *Ascl1* | 160,35 | 109,86 | 0,0155 | -1,46 |
| *Syp* | 13666,31 | 11421,17 | 0,0156 | -1,20 |
| *Ckap4* | 196,27 | 151,72 | 0,0156 | -1,29 |
| *Tmem217* | 4,37 | 1,20 | 0,0156 | -3,65 |
| *Anxa11* | 1258,37 | 1078,69 | 0,0157 | -1,17 |
| *Hsd17b7* | 63,68 | 46,74 | 0,0159 | -1,36 |
| *Rcl1* | 311,47 | 372,09 | 0,0159 | 1,19 |
| *Rnls* | 114,00 | 143,72 | 0,0160 | 1,26 |
| *Gpr17* | 1220,53 | 1034,66 | 0,0160 | -1,18 |
| *Clec12a* | 9,77 | 18,52 | 0,0160 | 1,90 |
| *Cyb561* | 832,94 | 956,61 | 0,0160 | 1,15 |
| *Homez* | 85,27 | 64,86 | 0,0161 | -1,31 |
| *Ptrhd1* | 87,06 | 120,93 | 0,0161 | 1,39 |
| *Plk3* | 110,01 | 83,13 | 0,0161 | -1,32 |
| *Mta3* | 907,49 | 785,66 | 0,0163 | -1,16 |
| *LOC688390* | 16,29 | 8,43 | 0,0164 | -1,93 |
| *Thrb* | 1135,86 | 952,94 | 0,0164 | -1,19 |
| *ENSRNOG00000069443* | 3,40 | 0,72 | 0,0164 | -4,70 |
| *Zbtb16* | 47,58 | 27,79 | 0,0165 | -1,71 |
| *Syn2* | 2564,59 | 2142,86 | 0,0165 | -1,20 |
| *Hacl1* | 229,32 | 272,63 | 0,0166 | 1,19 |
| *Bbof1* | 61,37 | 84,12 | 0,0166 | 1,37 |
| *Mpdz* | 581,43 | 451,82 | 0,0167 | -1,29 |
| *Clic5* | 290,88 | 233,38 | 0,0167 | -1,25 |
| *Nfib* | 121,74 | 70,68 | 0,0168 | -1,72 |
| *Tmem178a* | 684,19 | 495,66 | 0,0169 | -1,38 |
| *Hectd4* | 2395,68 | 2103,75 | 0,0170 | -1,14 |
| *C4bpa* | 10,57 | 19,47 | 0,0170 | 1,84 |
| *Gnpda2* | 263,45 | 311,05 | 0,0170 | 1,18 |
| *Wasl* | 1685,65 | 1532,05 | 0,0171 | -1,10 |
| *Cyp3a9* | 81,58 | 62,16 | 0,0172 | -1,31 |
| *Kcnc2* | 686,01 | 499,03 | 0,0173 | -1,37 |
| *AABR07017236.2* | 0,20 | 2,94 | 0,0173 | 14,39 |
| *R3hdm2* | 2415,75 | 2175,38 | 0,0175 | -1,11 |
| *Syt1* | 3757,32 | 2768,08 | 0,0176 | -1,36 |
| *Tceal6* | 927,05 | 1093,51 | 0,0176 | 1,18 |
| *Smim45* | 506,89 | 603,31 | 0,0177 | 1,19 |
| *Hcn1* | 132,35 | 92,53 | 0,0178 | -1,43 |
| *Rasgrf2* | 150,45 | 102,79 | 0,0178 | -1,46 |
| *Cers4* | 319,75 | 226,51 | 0,0179 | -1,41 |
| *Dhcr24* | 2138,11 | 1761,96 | 0,0179 | -1,21 |
| *Sec22a* | 171,93 | 213,15 | 0,0180 | 1,24 |
| *Ptgis* | 8,38 | 16,87 | 0,0180 | 2,01 |
| *Clec11a* | 121,94 | 90,87 | 0,0182 | -1,34 |
| *LOC120093125* | 2,20 | 12,43 | 0,0182 | 5,66 |
| *Fundc1* | 422,48 | 477,59 | 0,0183 | 1,13 |
| *Dzip1l* | 426,10 | 498,46 | 0,0184 | 1,17 |
| *Nrxn2* | 4902,51 | 4265,30 | 0,0186 | -1,15 |
| *Lxn* | 1217,43 | 1005,80 | 0,0187 | -1,21 |
| *Pgr* | 83,34 | 54,80 | 0,0187 | -1,52 |
| *Sorbs2* | 2317,77 | 1967,14 | 0,0187 | -1,18 |
| *Il17b* | 4,61 | 15,57 | 0,0188 | 3,38 |
| *ENSRNOG00000069313* | 226,41 | 269,49 | 0,0188 | 1,19 |
| *Hbs1l* | 1058,25 | 961,71 | 0,0189 | -1,10 |
| *MAST1* | 625,69 | 474,48 | 0,0189 | -1,32 |
| *Ccdc85a* | 214,19 | 165,32 | 0,0189 | -1,30 |
| *Tpte2* | 8,16 | 3,27 | 0,0189 | -2,50 |
| *Lipm* | 52,89 | 31,58 | 0,0190 | -1,67 |
| *Bgn* | 333,84 | 277,86 | 0,0190 | -1,20 |
| *Neurod6* | 475,17 | 337,73 | 0,0190 | -1,41 |
| *LOC100360573* | 281,57 | 347,65 | 0,0191 | 1,23 |
| *Gsx2* | 6,37 | 2,31 | 0,0192 | -2,76 |
| *Ttpa* | 109,39 | 138,87 | 0,0192 | 1,27 |
| *Panx2* | 1949,43 | 1714,85 | 0,0193 | -1,14 |
| *Zfp286a* | 147,83 | 183,57 | 0,0194 | 1,24 |
| *Dnm1* | 11559,59 | 9717,79 | 0,0194 | -1,19 |
| *Ogfrl1* | 487,29 | 409,92 | 0,0194 | -1,19 |
| *LOC100912617* | 0,80 | 4,03 | 0,0194 | 5,04 |
| *Scp2* | 3052,03 | 3529,19 | 0,0196 | 1,16 |
| *Chrna5* | 121,73 | 77,78 | 0,0196 | -1,57 |
| *Rdx* | 1505,81 | 1343,51 | 0,0197 | -1,12 |
| *Amph* | 2224,93 | 1879,87 | 0,0197 | -1,18 |
| *Etnk2* | 32,30 | 47,38 | 0,0198 | 1,47 |
| *AABR07039334.1* | 275,25 | 219,67 | 0,0198 | -1,25 |
| *Spout1* | 375,62 | 440,78 | 0,0200 | 1,17 |
| *Ap2b1* | 5172,47 | 4623,98 | 0,0201 | -1,12 |
| *Cd109* | 9,98 | 5,10 | 0,0203 | -1,96 |
| *Cth* | 60,23 | 80,95 | 0,0203 | 1,34 |
| *Fbxl2* | 454,93 | 392,07 | 0,0204 | -1,16 |
| *Tmem130* | 7909,76 | 7258,74 | 0,0204 | -1,09 |
| *Lmnb1* | 150,15 | 116,10 | 0,0204 | -1,29 |
| *Etl4* | 1298,78 | 1028,00 | 0,0205 | -1,26 |
| *Hormad1* | 0,20 | 2,98 | 0,0205 | 14,79 |
| *Zeb2* | 1156,68 | 964,34 | 0,0205 | -1,20 |
| *Shank1* | 2964,59 | 2242,67 | 0,0205 | -1,32 |
| *Cadps2* | 716,98 | 503,48 | 0,0205 | -1,42 |
| *Meaf6* | 912,00 | 802,88 | 0,0206 | -1,14 |
| *Slc16a10* | 15,92 | 25,96 | 0,0208 | 1,63 |
| *Clip1* | 967,89 | 844,48 | 0,0208 | -1,15 |
| *Pde1a* | 560,32 | 385,46 | 0,0208 | -1,45 |
| *Sebox* | 1,61 | 6,28 | 0,0209 | 3,89 |
| *ENSRNOG00000060956* | 2428,54 | 2690,14 | 0,0209 | 1,11 |
| *Adcy2* | 1484,13 | 1242,27 | 0,0212 | -1,19 |
| *Olr59* | 22,15 | 13,58 | 0,0212 | -1,63 |
| *Crim1* | 414,19 | 308,66 | 0,0212 | -1,34 |
| *Stt3b* | 636,75 | 548,28 | 0,0212 | -1,16 |
| *Tctn1* | 217,92 | 254,73 | 0,0213 | 1,17 |
| *Clip3* | 7104,00 | 6369,22 | 0,0213 | -1,12 |
| *Surf6* | 327,70 | 380,16 | 0,0214 | 1,16 |
| *Zfyve21* | 333,62 | 411,84 | 0,0215 | 1,23 |
| *Dgkz* | 5601,03 | 4743,51 | 0,0215 | -1,18 |
| *Tbrg4* | 386,35 | 445,47 | 0,0216 | 1,15 |
| *Pten* | 766,92 | 595,23 | 0,0217 | -1,29 |
| *Wfdc2* | 90,62 | 119,04 | 0,0217 | 1,31 |
| *Aen* | 83,74 | 107,27 | 0,0218 | 1,28 |
| *Nt5dc1* | 95,41 | 75,68 | 0,0218 | -1,26 |
| *Adora2a* | 4996,01 | 5902,69 | 0,0218 | 1,18 |
| *Rnf138l1* | 10,48 | 4,46 | 0,0219 | -2,35 |
| *Cpxm1* | 48,24 | 62,86 | 0,0220 | 1,30 |
| *Il12a* | 22,95 | 35,48 | 0,0220 | 1,55 |
| *ENSRNOG00000064754* | 225,68 | 303,05 | 0,0220 | 1,34 |
| *Tm4sf4* | 1,81 | 6,73 | 0,0220 | 3,72 |
| *Napb* | 3206,57 | 2569,09 | 0,0221 | -1,25 |
| *Cntnap5b* | 21,11 | 11,89 | 0,0221 | -1,77 |
| *RGD1564492* | 55,89 | 73,99 | 0,0221 | 1,32 |
| *Apobr* | 28,89 | 42,75 | 0,0222 | 1,48 |
| *Slc6a17* | 4501,31 | 3821,70 | 0,0222 | -1,18 |
| *Mamdc2* | 61,84 | 43,35 | 0,0222 | -1,43 |
| *Aldh16a1* | 210,79 | 248,24 | 0,0223 | 1,18 |
| *Cap1* | 8280,63 | 9393,26 | 0,0223 | 1,13 |
| *Rbl1* | 33,93 | 23,63 | 0,0223 | -1,44 |
| *Slc25a5* | 3578,69 | 4195,63 | 0,0224 | 1,17 |
| *Osbpl11* | 252,35 | 209,47 | 0,0226 | -1,20 |
| *Mmd2* | 2448,34 | 2820,05 | 0,0226 | 1,15 |
| *Phf23* | 488,17 | 559,40 | 0,0227 | 1,15 |
| *Map1b* | 11869,77 | 9994,49 | 0,0228 | -1,19 |
| *AABR07058017.2* | 5,16 | 1,46 | 0,0229 | -3,53 |
| *Smap1* | 1171,20 | 1015,82 | 0,0229 | -1,15 |
| *Ptprn* | 9381,13 | 8575,97 | 0,0229 | -1,09 |
| *Atxn2* | 1157,08 | 1038,34 | 0,0229 | -1,11 |
| *Fam222a* | 376,18 | 432,30 | 0,0229 | 1,15 |
| *Ccn3* | 1038,47 | 617,94 | 0,0230 | -1,68 |
| *Sgtb* | 3226,68 | 2925,92 | 0,0230 | -1,10 |
| *Arhgap44* | 2249,98 | 1895,45 | 0,0231 | -1,19 |
| *P2rx6* | 29,74 | 58,42 | 0,0231 | 1,96 |
| *Otud7a* | 717,76 | 628,14 | 0,0231 | -1,14 |
| *Nwd2* | 84,20 | 53,21 | 0,0232 | -1,58 |
| *LOC120096327* | 0,40 | 3,21 | 0,0232 | 8,09 |
| *Arhgef16* | 15,91 | 26,17 | 0,0233 | 1,64 |
| *ENSRNOG00000064080* | 515,62 | 400,39 | 0,0233 | -1,29 |
| *Rcn3* | 154,18 | 119,49 | 0,0234 | -1,29 |
| *Lamp5* | 2494,17 | 3032,13 | 0,0234 | 1,22 |
| *Optn* | 928,57 | 777,39 | 0,0235 | -1,19 |
| *Stap2* | 47,91 | 68,54 | 0,0235 | 1,43 |
| *Ubash3a* | 0,79 | 3,85 | 0,0236 | 4,88 |
| *Lratd2* | 332,04 | 378,65 | 0,0236 | 1,14 |
| *Gap43* | 5231,61 | 4697,37 | 0,0238 | -1,11 |
| *Cmip* | 1123,70 | 867,27 | 0,0238 | -1,30 |
| *Emx1* | 67,73 | 39,00 | 0,0238 | -1,74 |
| *Oasl2* | 53,27 | 75,99 | 0,0240 | 1,43 |
| *C5h9orf152* | 1,60 | 6,13 | 0,0241 | 3,83 |
| *ENSRNOG00000064622* | 1,39 | 4,79 | 0,0241 | 3,43 |
| *Dlx1* | 274,47 | 220,59 | 0,0241 | -1,24 |
| *Megf9* | 441,92 | 327,23 | 0,0242 | -1,35 |
| *RGD1566093* | 239,69 | 202,02 | 0,0242 | -1,19 |
| *Ucp2* | 410,63 | 350,45 | 0,0243 | -1,17 |
| *Tspan5* | 2278,47 | 2076,63 | 0,0243 | -1,10 |
| *Foxe1* | 3,58 | 0,93 | 0,0243 | -3,84 |
| *ENSRNOG00000063744* | 6,74 | 14,91 | 0,0244 | 2,21 |
| *Rpl26-ps2* | 317,08 | 425,09 | 0,0244 | 1,34 |
| *Sv2b* | 4101,32 | 3220,78 | 0,0245 | -1,27 |
| *Mgarp* | 3,16 | 8,20 | 0,0245 | 2,59 |
| *Dus2* | 154,19 | 197,06 | 0,0245 | 1,28 |
| *Cpne8* | 296,37 | 253,64 | 0,0247 | -1,17 |
| *Cmtr2* | 128,79 | 154,21 | 0,0247 | 1,20 |
| *Dpysl2* | 12003,36 | 10855,74 | 0,0247 | -1,11 |
| *Tmem145* | 179,09 | 139,09 | 0,0248 | -1,29 |
| *Uqcc1-ps1* | 4,61 | 1,27 | 0,0248 | -3,63 |
| *RGD1564409* | 58,29 | 29,50 | 0,0248 | -1,98 |
| *Micall2* | 19,68 | 29,84 | 0,0249 | 1,52 |
| *Pcsk4* | 141,67 | 104,30 | 0,0249 | -1,36 |
| *Sorl1* | 1819,88 | 1515,29 | 0,0250 | -1,20 |
| *Bean1* | 72,86 | 57,43 | 0,0250 | -1,27 |
| *RGD1561796* | 80,66 | 103,84 | 0,0250 | 1,29 |
| *Ccdc71* | 643,52 | 572,60 | 0,0252 | -1,12 |
| *Ip6k2* | 1036,85 | 1162,57 | 0,0255 | 1,12 |
| *Lratd1* | 1062,51 | 1198,62 | 0,0255 | 1,13 |
| *Homer1* | 1841,13 | 1554,21 | 0,0255 | -1,18 |
| *Pknox2* | 427,87 | 364,97 | 0,0255 | -1,17 |
| *Crmp1* | 1866,69 | 1617,63 | 0,0256 | -1,15 |
| *Cfap47* | 11,95 | 6,32 | 0,0258 | -1,89 |
| *Atg16l2* | 51,48 | 71,70 | 0,0259 | 1,39 |
| *Wdr53* | 100,90 | 125,87 | 0,0260 | 1,25 |
| *Iqsec3* | 1216,93 | 980,81 | 0,0261 | -1,24 |
| *Sfxn3* | 886,97 | 746,98 | 0,0261 | -1,19 |
| *Kyat3* | 160,81 | 199,58 | 0,0261 | 1,24 |
| *Lsm11* | 107,71 | 77,80 | 0,0261 | -1,38 |
| *Adcyap1* | 79,53 | 49,32 | 0,0261 | -1,61 |
| *Sobp* | 594,45 | 461,92 | 0,0261 | -1,29 |
| *Snrpel1* | 32,24 | 46,18 | 0,0262 | 1,43 |
| *Ace2* | 22,09 | 34,87 | 0,0263 | 1,58 |
| *Ppm1h* | 285,46 | 228,95 | 0,0263 | -1,25 |
| *ENSRNOG00000069729* | 7,57 | 3,66 | 0,0266 | -2,07 |
| *Csf1r* | 1640,88 | 1851,35 | 0,0266 | 1,13 |
| *Nxt1* | 75,85 | 97,99 | 0,0266 | 1,29 |
| *Cck* | 3500,84 | 2220,49 | 0,0267 | -1,58 |
| *AABR07004881.1* | 293,73 | 232,64 | 0,0267 | -1,26 |
| *ENSRNOG00000062933* | 23,34 | 33,21 | 0,0267 | 1,42 |
| *Ncald* | 1846,95 | 1426,07 | 0,0267 | -1,30 |
| *Ephx3* | 0,98 | 5,39 | 0,0268 | 5,48 |
| *ENSRNOG00000070152* | 6,39 | 11,81 | 0,0270 | 1,85 |
| *Aste1* | 47,97 | 63,47 | 0,0270 | 1,32 |
| *Tfap2d* | 26,80 | 15,21 | 0,0272 | -1,76 |
| *Sfpq* | 2269,64 | 1986,44 | 0,0272 | -1,14 |
| *Tubb3* | 5327,24 | 4676,66 | 0,0273 | -1,14 |
| *Zfp623* | 147,85 | 177,94 | 0,0273 | 1,20 |
| *Avil* | 14,56 | 23,98 | 0,0274 | 1,65 |
| *Tbc1d22b* | 637,25 | 563,82 | 0,0274 | -1,13 |
| *Pnma3* | 211,99 | 164,57 | 0,0276 | -1,29 |
| *Ubr7* | 556,44 | 628,14 | 0,0276 | 1,13 |
| *LOC120093361* | 0,39 | 5,59 | 0,0277 | 14,31 |
| *Vom1r47* | 1,61 | 5,17 | 0,0277 | 3,21 |
| *LOC108352348* | 190,85 | 273,63 | 0,0277 | 1,43 |
| *Srsf4* | 1014,25 | 893,81 | 0,0278 | -1,13 |
| *Rxrb* | 1127,20 | 1280,05 | 0,0279 | 1,14 |
| *Dnajc6* | 3699,82 | 3208,20 | 0,0280 | -1,15 |
| *Tbc1d9* | 1150,34 | 1006,27 | 0,0280 | -1,14 |
| *Art4* | 19,26 | 11,73 | 0,0280 | -1,64 |
| *Fads6* | 395,60 | 443,77 | 0,0280 | 1,12 |
| *Nrn1* | 791,98 | 504,01 | 0,0281 | -1,57 |
| *Gucd1* | 258,63 | 305,55 | 0,0282 | 1,18 |
| *Tcp11l2* | 313,57 | 366,36 | 0,0282 | 1,17 |
| *Slc26a11* | 97,60 | 75,26 | 0,0282 | -1,30 |
| *Sgcg* | 3,16 | 9,54 | 0,0284 | 3,01 |
| *Sirt3* | 688,84 | 794,83 | 0,0285 | 1,15 |
| *ENSRNOG00000068298* | 481,66 | 571,10 | 0,0285 | 1,19 |
| *Ing1* | 431,75 | 373,84 | 0,0286 | -1,15 |
| *Akt2* | 738,47 | 652,91 | 0,0287 | -1,13 |
| *Satb1* | 373,87 | 278,85 | 0,0287 | -1,34 |
| *Arid3b* | 103,91 | 131,70 | 0,0287 | 1,27 |
| *Ogfr* | 844,61 | 1002,72 | 0,0287 | 1,19 |
| *Cfb* | 279,80 | 338,37 | 0,0288 | 1,21 |
| *Mef2c* | 858,68 | 637,64 | 0,0290 | -1,35 |
| *Vrk2* | 34,33 | 48,87 | 0,0291 | 1,42 |
| *Acot3* | 155,07 | 182,23 | 0,0291 | 1,18 |
| *Atp4a* | 0,80 | 3,90 | 0,0291 | 4,85 |
| *Pip5k1c* | 4526,12 | 4103,97 | 0,0292 | -1,10 |
| *Upp1* | 87,27 | 122,43 | 0,0292 | 1,40 |
| *Syt13* | 2393,52 | 2178,69 | 0,0293 | -1,10 |
| *Ppp1r10* | 864,42 | 739,99 | 0,0293 | -1,17 |
| *Golga7b* | 269,81 | 193,58 | 0,0294 | -1,39 |
| *Aida* | 691,30 | 613,90 | 0,0294 | -1,13 |
| *Dmkn* | 521,70 | 616,46 | 0,0295 | 1,18 |
| *ENSRNOG00000068226* | 212,83 | 259,63 | 0,0295 | 1,22 |
| *Atp10a* | 148,17 | 185,69 | 0,0296 | 1,25 |
| *Cdkn2b* | 7,18 | 2,80 | 0,0297 | -2,57 |
| *Dnajc12* | 244,97 | 198,84 | 0,0297 | -1,23 |
| *Psme3* | 1387,15 | 1277,09 | 0,0298 | -1,09 |
| *B3galt2* | 569,12 | 461,35 | 0,0298 | -1,23 |
| *Prdm8* | 97,48 | 64,00 | 0,0299 | -1,52 |
| *Abcg2* | 213,07 | 267,72 | 0,0299 | 1,26 |
| *Atad2b* | 77,40 | 53,91 | 0,0299 | -1,44 |
| *Cacng5* | 132,35 | 110,37 | 0,0300 | -1,20 |
| *Cpxm2* | 9,53 | 19,52 | 0,0300 | 2,05 |
| *LOC103690870* | 31,73 | 20,71 | 0,0301 | -1,53 |
| *Hmox1* | 68,64 | 93,30 | 0,0301 | 1,36 |
| *Cntnap4* | 314,83 | 248,63 | 0,0301 | -1,27 |
| *Shank2* | 1924,45 | 1692,58 | 0,0303 | -1,14 |
| *Prmt8* | 1011,77 | 916,00 | 0,0304 | -1,10 |
| *Eno3* | 152,49 | 179,25 | 0,0305 | 1,18 |
| *Dclk3* | 1077,92 | 1223,43 | 0,0306 | 1,13 |
| *Nrsn1* | 2231,42 | 1956,86 | 0,0307 | -1,14 |
| *Fndc5* | 1424,96 | 1630,84 | 0,0309 | 1,14 |
| *Mppe1* | 167,84 | 202,65 | 0,0311 | 1,21 |
| *Sult1d1* | 246,10 | 297,44 | 0,0311 | 1,21 |
| *Zbtb18* | 2084,94 | 1811,46 | 0,0312 | -1,15 |
| *LOC102550011* | 55,53 | 76,11 | 0,0312 | 1,37 |
| *Lrrc23* | 273,39 | 332,03 | 0,0313 | 1,21 |
| *Gpr62* | 151,88 | 108,91 | 0,0315 | -1,39 |
| *Mpeg1* | 1142,20 | 1470,17 | 0,0315 | 1,29 |
| *Atp6v1a* | 5858,06 | 5214,83 | 0,0315 | -1,12 |
| *Taf13* | 572,93 | 664,80 | 0,0317 | 1,16 |
| *Pcdhb10* | 34,83 | 48,85 | 0,0318 | 1,40 |
| *Nckap5l* | 364,00 | 417,27 | 0,0318 | 1,15 |
| *Mfsd12* | 214,73 | 176,51 | 0,0320 | -1,22 |
| *Scml4* | 46,95 | 33,58 | 0,0320 | -1,40 |
| *Cbx1* | 269,64 | 225,96 | 0,0320 | -1,19 |
| *Chrna4* | 415,07 | 301,59 | 0,0322 | -1,38 |
| *Gpr149* | 174,85 | 215,27 | 0,0322 | 1,23 |
| *Gng10* | 1512,23 | 1752,28 | 0,0322 | 1,16 |
| *P4ha3* | 47,01 | 27,30 | 0,0324 | -1,72 |
| *Ncaph* | 26,66 | 16,86 | 0,0325 | -1,58 |
| *Trabd* | 217,96 | 258,52 | 0,0325 | 1,19 |
| *Igsf1* | 1189,08 | 1379,25 | 0,0326 | 1,16 |
| *Acad9* | 640,49 | 728,24 | 0,0327 | 1,14 |
| *ENSRNOG00000065147* | 248,61 | 307,77 | 0,0329 | 1,24 |
| *Stambpl1* | 530,55 | 472,13 | 0,0329 | -1,12 |
| *Disp2* | 3043,36 | 2607,68 | 0,0330 | -1,17 |
| *Unc5d* | 39,74 | 25,67 | 0,0332 | -1,55 |
| *Pfkfb4* | 70,37 | 88,53 | 0,0333 | 1,26 |
| *Bcl2a1* | 19,56 | 30,53 | 0,0333 | 1,56 |
| *Cnbp* | 5378,13 | 6024,46 | 0,0336 | 1,12 |
| *Elf1* | 118,40 | 143,08 | 0,0336 | 1,21 |
| *Adamtsl5* | 48,46 | 64,25 | 0,0336 | 1,33 |
| *Rab6b* | 5623,38 | 4840,09 | 0,0337 | -1,16 |
| *Fbf1* | 795,66 | 885,37 | 0,0338 | 1,11 |
| *Rarres2* | 349,78 | 439,66 | 0,0338 | 1,26 |
| *LOC100365810* | 610,35 | 760,99 | 0,0338 | 1,25 |
| *ENSRNOG00000014538* | 4,00 | 0,67 | 0,0340 | -6,00 |
| *Limk1* | 763,54 | 613,80 | 0,0340 | -1,24 |
| *St6gal2* | 200,33 | 137,93 | 0,0340 | -1,45 |
| *Btbd3* | 538,40 | 425,40 | 0,0341 | -1,27 |
| *Slc25a26* | 217,42 | 252,18 | 0,0341 | 1,16 |
| *Snora52* | 0,99 | 5,45 | 0,0341 | 5,52 |
| *Rft1* | 127,76 | 151,47 | 0,0342 | 1,19 |
| *Atp1b1* | 18727,59 | 17011,36 | 0,0342 | -1,10 |
| *Ubash3b* | 119,05 | 92,93 | 0,0342 | -1,28 |
| *Dscaml1* | 469,63 | 374,25 | 0,0344 | -1,25 |
| *Mfap3l* | 668,42 | 536,52 | 0,0344 | -1,25 |
| *Zfp108* | 176,87 | 205,80 | 0,0346 | 1,16 |
| *Nod2* | 3,98 | 10,97 | 0,0347 | 2,76 |
| *Vip* | 134,51 | 92,29 | 0,0348 | -1,46 |
| *Sfr1* | 1232,62 | 1397,28 | 0,0349 | 1,13 |
| *Ggh* | 443,74 | 514,13 | 0,0349 | 1,16 |
| *Myh6* | 40,71 | 60,22 | 0,0350 | 1,48 |
| *Ccdc190* | 28,85 | 42,46 | 0,0351 | 1,47 |
| *Cpt2* | 157,78 | 191,61 | 0,0351 | 1,21 |
| *Adgra1* | 373,47 | 282,63 | 0,0352 | -1,32 |
| *Ccdc177* | 151,00 | 111,13 | 0,0353 | -1,36 |
| *S1pr2* | 69,16 | 87,57 | 0,0353 | 1,27 |
| *Abcg1* | 1441,93 | 1589,65 | 0,0354 | 1,10 |
| *Vwc2l* | 35,23 | 21,41 | 0,0355 | -1,65 |
| *Mex3a* | 31,27 | 21,71 | 0,0355 | -1,44 |
| *Foxd1* | 2,61 | 7,04 | 0,0356 | 2,70 |
| *Myom3* | 33,48 | 20,34 | 0,0356 | -1,65 |
| *Tamalin* | 586,37 | 509,81 | 0,0357 | -1,15 |
| *Polr3gl* | 207,79 | 247,48 | 0,0358 | 1,19 |
| *Med8* | 410,95 | 469,57 | 0,0358 | 1,14 |
| *ENSRNOG00000066309* | 34,86 | 49,17 | 0,0359 | 1,41 |
| *Entrep2* | 702,99 | 621,85 | 0,0360 | -1,13 |
| *Snx31* | 2,81 | 0,47 | 0,0363 | -6,00 |
| *Inhbb* | 66,49 | 51,98 | 0,0363 | -1,28 |
| *Acsl4* | 1496,54 | 1324,94 | 0,0363 | -1,13 |
| *Pfkp* | 6507,73 | 5959,02 | 0,0364 | -1,09 |
| *Tusc3* | 3653,45 | 4083,23 | 0,0365 | 1,12 |
| *Nav3* | 576,23 | 425,51 | 0,0365 | -1,35 |
| *Meis3* | 356,23 | 415,09 | 0,0367 | 1,17 |
| *Snrpg* | 224,45 | 285,56 | 0,0368 | 1,27 |
| *Nfkb2* | 59,89 | 78,63 | 0,0371 | 1,31 |
| *Rasal1* | 538,02 | 398,97 | 0,0372 | -1,35 |
| *Kcnk2* | 2014,33 | 2297,71 | 0,0372 | 1,14 |
| *Cdh13* | 2837,31 | 2558,88 | 0,0373 | -1,11 |
| *Tifab* | 165,76 | 213,64 | 0,0373 | 1,29 |
| *Timm9* | 230,81 | 269,94 | 0,0376 | 1,17 |
| *Vash1* | 769,76 | 639,49 | 0,0376 | -1,20 |
| *Trim34* | 48,48 | 64,15 | 0,0377 | 1,32 |
| *Cxcr4* | 25,96 | 36,44 | 0,0377 | 1,40 |
| *Ehmt2* | 2815,10 | 3105,73 | 0,0377 | 1,10 |
| *Slc37a4* | 350,63 | 404,15 | 0,0378 | 1,15 |
| *Clstn3* | 3903,33 | 3416,93 | 0,0378 | -1,14 |
| *Tm7sf3* | 787,39 | 914,08 | 0,0378 | 1,16 |
| *Pole3* | 510,24 | 580,22 | 0,0378 | 1,14 |
| *Camk2d* | 590,20 | 483,94 | 0,0379 | -1,22 |
| *Ppme1* | 2662,97 | 2393,74 | 0,0380 | -1,11 |
| *Iqsec1* | 4338,28 | 3988,07 | 0,0380 | -1,09 |
| *ENSRNOG00000067762* | 200,78 | 168,59 | 0,0381 | -1,19 |
| *Kin* | 138,88 | 166,42 | 0,0382 | 1,20 |
| *Ncbp2* | 961,77 | 1072,19 | 0,0382 | 1,11 |
| *Bpnt1* | 897,77 | 974,53 | 0,0382 | 1,09 |
| *ENSRNOG00000066939* | 44,42 | 32,51 | 0,0382 | -1,37 |
| *Ndc1* | 283,59 | 324,22 | 0,0383 | 1,14 |
| *Prrg3* | 199,35 | 152,29 | 0,0383 | -1,31 |
| *ENSRNOG00000065152* | 0,20 | 2,28 | 0,0383 | 11,66 |
| *Tmem17* | 207,91 | 244,74 | 0,0383 | 1,18 |
| *Mkrn1* | 1706,00 | 1865,20 | 0,0384 | 1,09 |
| *Gnb4* | 430,62 | 342,59 | 0,0385 | -1,26 |
| *AC109737.1* | 5,16 | 1,51 | 0,0385 | -3,41 |
| *Borcs5* | 292,91 | 249,77 | 0,0386 | -1,17 |
| *Higd1a* | 950,32 | 1094,74 | 0,0386 | 1,15 |
| *Chp1* | 1402,36 | 1274,55 | 0,0386 | -1,10 |
| *ENSRNOG00000064308* | 145,34 | 170,07 | 0,0387 | 1,17 |
| *Hacd4* | 24,14 | 15,93 | 0,0388 | -1,51 |
| *Tox* | 207,04 | 174,28 | 0,0389 | -1,19 |
| *Sfrp2* | 74,95 | 57,71 | 0,0390 | -1,30 |
| *Asb6* | 361,12 | 416,27 | 0,0390 | 1,15 |
| *Usp45* | 237,62 | 183,14 | 0,0391 | -1,30 |
| *LOC100359916* | 272,61 | 234,80 | 0,0392 | -1,16 |
| *ENSRNOG00000064885* | 7,21 | 2,90 | 0,0393 | -2,49 |
| *Map3k6* | 18,82 | 11,02 | 0,0394 | -1,71 |
| *Clpx* | 488,94 | 551,69 | 0,0394 | 1,13 |
| *St3gal3* | 599,55 | 678,55 | 0,0395 | 1,13 |
| *Msx1* | 19,05 | 29,44 | 0,0395 | 1,55 |
| *Rnf39* | 102,90 | 84,62 | 0,0396 | -1,22 |
| *ENSRNOG00000066444* | 0,79 | 3,38 | 0,0397 | 4,30 |
| *Slc6a7* | 359,33 | 259,89 | 0,0397 | -1,38 |
| *Capza2* | 3684,93 | 3984,07 | 0,0398 | 1,08 |
| *Exd1* | 7,55 | 3,62 | 0,0398 | -2,09 |
| *Ndufa4l2* | 87,85 | 106,54 | 0,0398 | 1,21 |
| *Nrp2* | 206,44 | 158,64 | 0,0399 | -1,30 |
| *Snx20* | 26,93 | 38,55 | 0,0400 | 1,43 |
| *Zfp473* | 14,51 | 24,75 | 0,0400 | 1,71 |
| *Plat* | 857,73 | 1091,64 | 0,0401 | 1,27 |
| *AABR07054262.1* | 5,97 | 2,47 | 0,0401 | -2,41 |
| *Tbx19* | 14,80 | 7,67 | 0,0401 | -1,93 |
| *Pi4ka* | 3195,27 | 2829,73 | 0,0402 | -1,13 |
| *Rtn4r* | 398,66 | 237,75 | 0,0402 | -1,68 |
| *Ccn2* | 722,29 | 574,79 | 0,0402 | -1,26 |
| *Cmtm4* | 111,39 | 78,37 | 0,0403 | -1,42 |
| *Cd37* | 60,13 | 79,45 | 0,0404 | 1,32 |
| *Doc2a* | 306,31 | 219,79 | 0,0404 | -1,39 |
| *Endov* | 95,22 | 122,52 | 0,0404 | 1,29 |
| *Slc24a3* | 1127,77 | 1005,63 | 0,0405 | -1,12 |
| *LOC100360522* | 1617,17 | 2093,38 | 0,0405 | 1,29 |
| *Frmd4a* | 596,62 | 529,08 | 0,0405 | -1,13 |
| *Haus6* | 41,68 | 29,09 | 0,0405 | -1,43 |
| *Cacng3* | 1805,06 | 1591,07 | 0,0405 | -1,13 |
| *Scn4b* | 14900,48 | 17907,34 | 0,0406 | 1,20 |
| *Uaca* | 151,61 | 203,87 | 0,0407 | 1,34 |
| *Dnai1* | 58,36 | 74,63 | 0,0410 | 1,28 |
| *Chrm1* | 1508,59 | 1328,85 | 0,0413 | -1,14 |
| *Cyp2d3* | 4,52 | 14,48 | 0,0413 | 3,20 |
| *Sox3* | 16,08 | 7,41 | 0,0414 | -2,17 |
| *Shoc1* | 5,15 | 1,53 | 0,0415 | -3,37 |
| *Osbpl3* | 49,68 | 35,91 | 0,0415 | -1,38 |
| *Ccsap* | 626,51 | 530,09 | 0,0416 | -1,18 |
| *LOC120102105* | 29,67 | 43,53 | 0,0417 | 1,47 |
| *Lsm14b* | 1452,41 | 1329,49 | 0,0417 | -1,09 |
| *C16h4orf47* | 21,89 | 13,74 | 0,0418 | -1,59 |
| *ENSRNOG00000066750* | 102,66 | 124,09 | 0,0418 | 1,21 |
| *Prkcg* | 3300,55 | 2847,49 | 0,0418 | -1,16 |
| *Cacna1g* | 511,41 | 392,40 | 0,0419 | -1,30 |
| *Fam118b* | 169,43 | 197,06 | 0,0420 | 1,16 |
| *Cap2* | 2310,74 | 2114,22 | 0,0420 | -1,09 |
| *Fam181b* | 33,47 | 23,00 | 0,0421 | -1,46 |
| *Egr3* | 1400,58 | 1147,85 | 0,0421 | -1,22 |
| *Ndufa10l1* | 231,88 | 158,51 | 0,0421 | -1,46 |
| *Rab1b-ps1* | 17,32 | 10,30 | 0,0423 | -1,68 |
| *Gpr146* | 350,04 | 401,49 | 0,0425 | 1,15 |
| *Sdhaf2* | 1001,50 | 1109,19 | 0,0425 | 1,11 |
| *Elmod3* | 199,81 | 230,68 | 0,0427 | 1,15 |
| *Slc38a11* | 5,18 | 1,48 | 0,0427 | -3,50 |
| *Chst7* | 158,22 | 128,07 | 0,0427 | -1,24 |
| *Rabggtb* | 1145,58 | 1276,53 | 0,0427 | 1,11 |
| *Jun* | 975,58 | 806,54 | 0,0428 | -1,21 |
| *Ccr10* | 7,15 | 13,30 | 0,0428 | 1,86 |
| *RGD1560523* | 39,57 | 28,51 | 0,0428 | -1,39 |
| *ENSRNOG00000062648* | 9,66 | 22,61 | 0,0428 | 2,34 |
| *Ccdc6* | 603,51 | 470,80 | 0,0429 | -1,28 |
| *Cyb5r1* | 481,93 | 405,55 | 0,0429 | -1,19 |
| *LOC681355* | 29,43 | 40,50 | 0,0429 | 1,38 |
| *Col4a5* | 126,25 | 159,13 | 0,0429 | 1,26 |
| *Cdk5r2* | 6549,19 | 5699,69 | 0,0431 | -1,15 |
| *Mboat7* | 1964,33 | 1778,36 | 0,0431 | -1,10 |
| *Chst13* | 17,91 | 9,46 | 0,0431 | -1,89 |
| *Cemip2* | 116,91 | 91,63 | 0,0431 | -1,28 |
| *Lrrn2* | 2189,96 | 1996,78 | 0,0433 | -1,10 |
| *Susd2* | 85,76 | 66,95 | 0,0433 | -1,28 |
| *Mir3561* | 2,79 | 6,62 | 0,0433 | 2,37 |
| *Akirin1* | 1102,72 | 1203,86 | 0,0434 | 1,09 |
| *Sel1l* | 2358,04 | 2045,09 | 0,0435 | -1,15 |
| *Slc30a3* | 373,35 | 219,67 | 0,0436 | -1,70 |
| *Ina* | 3472,21 | 3094,42 | 0,0438 | -1,12 |
| *Fitm2* | 365,93 | 312,66 | 0,0438 | -1,17 |
| *Acsm5* | 149,69 | 201,76 | 0,0440 | 1,35 |
| *Col3a1* | 50,44 | 71,56 | 0,0440 | 1,42 |
| *Cpeb2* | 404,11 | 320,86 | 0,0440 | -1,26 |
| *Itk* | 142,90 | 184,14 | 0,0443 | 1,29 |
| *L2hgdh* | 122,83 | 149,67 | 0,0445 | 1,22 |
| *Ipcef1* | 978,80 | 720,54 | 0,0446 | -1,36 |
| *Ldb2* | 783,94 | 713,42 | 0,0446 | -1,10 |
| *Zfp780b* | 118,70 | 92,20 | 0,0447 | -1,29 |
| *ENSRNOG00000050514* | 133,78 | 101,86 | 0,0448 | -1,31 |
| *Ctnnd2* | 3261,22 | 2911,76 | 0,0449 | -1,12 |
| *Pros1* | 157,60 | 182,60 | 0,0453 | 1,16 |
| *Inf2* | 6768,75 | 7530,91 | 0,0454 | 1,11 |
| *Sclt1* | 174,22 | 202,17 | 0,0455 | 1,16 |
| *Slc26a4* | 7,97 | 3,16 | 0,0456 | -2,52 |
| *Tmem97* | 1058,58 | 938,74 | 0,0457 | -1,13 |
| *Ccdc82* | 304,78 | 261,10 | 0,0460 | -1,17 |
| *Sv2a* | 4594,07 | 4077,64 | 0,0460 | -1,13 |
| *LOC120099460* | 3,19 | 7,17 | 0,0462 | 2,25 |
| *Rap2a* | 1111,78 | 936,26 | 0,0462 | -1,19 |
| *Them4* | 395,74 | 447,95 | 0,0465 | 1,13 |
| *Ccdc117* | 179,92 | 217,38 | 0,0468 | 1,21 |
| *Cd34* | 152,40 | 115,68 | 0,0470 | -1,32 |
| *Nudt18* | 419,66 | 484,90 | 0,0471 | 1,16 |
| *Tdh* | 14,71 | 22,60 | 0,0471 | 1,54 |
| *Galns* | 105,35 | 86,70 | 0,0472 | -1,22 |
| *Hgf* | 19,29 | 11,70 | 0,0472 | -1,65 |
| *Cpa2* | 32,58 | 43,86 | 0,0473 | 1,35 |
| *LOC100361457* | 1674,58 | 1497,78 | 0,0473 | -1,12 |
| *Sulf2* | 1786,78 | 1535,60 | 0,0473 | -1,16 |
| *Fdft1* | 1596,48 | 1399,58 | 0,0475 | -1,14 |
| *Acp4* | 20,86 | 31,69 | 0,0477 | 1,52 |
| *Necab2* | 2533,37 | 2965,13 | 0,0477 | 1,17 |
| *Magi1* | 767,35 | 646,84 | 0,0478 | -1,19 |
| *ENSRNOG00000071035* | 102,49 | 123,67 | 0,0478 | 1,21 |
| *Ndel1* | 1199,63 | 1098,93 | 0,0478 | -1,09 |
| *Hacd1* | 48,49 | 66,70 | 0,0479 | 1,38 |
| *Rbm20* | 16,40 | 8,94 | 0,0479 | -1,83 |
| *Rmnd5b* | 375,08 | 310,74 | 0,0481 | -1,21 |
| *Pgrmc2* | 1409,37 | 1292,12 | 0,0482 | -1,09 |
| *Angptl1* | 16,54 | 9,68 | 0,0484 | -1,71 |
| *Tox3* | 149,41 | 119,82 | 0,0484 | -1,25 |
| *Stk4* | 562,51 | 498,97 | 0,0486 | -1,13 |
| *Trmt2a* | 228,94 | 262,74 | 0,0486 | 1,15 |
| *ENSRNOG00000066174* | 7,77 | 13,54 | 0,0487 | 1,74 |
| *AABR07019088.1* | 4,42 | 1,29 | 0,0488 | -3,43 |
| *Gpc4* | 204,06 | 160,25 | 0,0489 | -1,27 |
| *Bend6* | 2428,55 | 2228,40 | 0,0490 | -1,09 |
| *Gemin2* | 108,96 | 138,38 | 0,0492 | 1,27 |
| *Raph1* | 382,14 | 313,75 | 0,0493 | -1,22 |
| *Kcnk1* | 977,72 | 835,42 | 0,0493 | -1,17 |
| *Vsnl1* | 4488,96 | 3464,73 | 0,0493 | -1,30 |
| *Mtfr1l* | 1030,51 | 1177,86 | 0,0493 | 1,14 |
| *Mir3072* | 4,99 | 10,62 | 0,0494 | 2,13 |
| *Cep295nl* | 2103,88 | 1904,59 | 0,0495 | -1,10 |
| *Hlx* | 24,13 | 15,76 | 0,0497 | -1,53 |
| *ENSRNOG00000063392* | 0,20 | 2,47 | 0,0498 | 12,12 |
| *Foxj3* | 1155,84 | 1026,77 | 0,0498 | -1,13 |
| *Spock1* | 1541,29 | 1262,21 | 0,0498 | -1,22 |
| *Strbp* | 1159,59 | 1070,26 | 0,0498 | -1,08 |
| *Tmem163* | 332,24 | 244,67 | 0,0499 | -1,36 |
| *Ccp110* | 650,51 | 525,16 | 0,0499 | -1,24 |
| *Cxxc5* | 1143,72 | 961,13 | 0,0500 | -1,19 |
