## Supplemental Table 2 for "Experimental change in personality: Overexpression of GDNF in the rat striatum converts the low exploratory phenotype into highly explorative"

**Supplementary Table ...** Primers used for quantifying gene expression by RT-qPCR.

F: Forward; R: Reverse; bp: base pairs

| **Gene** | **F primer sequence**  **(5’ to 3’)** | **R primer sequence**  **(5’ to 3’)** | **Product**  **(pb)** |
| --- | --- | --- | --- |
| *Vmat1* | GCCTTCGAAAGTGTCTCCTG | GCCAACACACCAAAGAGGTT | 243 |
| *Pdzd7* | TGAGGACGAAGCTGGGAATG | CCTGAGATGCTGATGCCCAA | 90 |
| *P2rx6* | GAACTGGGAGCATGGCTTCT | TGTCCCATTCCTGGTAGCCT | 192 |
| *Mettl3* | ATGTGCAGCCCAACTGGATT | CTGTGCTTAAACCGGGCAAC | 88 |
| *Zfp692* | CTCCAACCGGCAGTATTTGAATC | CAGGGCAGCAAAACGACTTC | 71 |
| *Lrpprc* | AGCCGAAAAGCAAGATGTCG | TTTCACATGCATGGCTCGTG | 100 |
| *Gapdh* | CTCTGCTCCTCCCTGTTCTA | GATACGGCCAAATCCGTTCA | 105 |

*Vmat1:* solute carrier family 18 member A1; *Pdzd7:* PDZ domain containing 7; *P2rx6,* purinergic receptor P2X 6; *Mettl3,* methyltransferase 3, N^6^-adenosine-methyltransferase complex catalytic subunit; *Zfp692,* zinc finger protein 692; *Lrpprc,* leucine-rich pentatricopeptide repeat containing, *Gapdh:* glyceraldehyde-3-phosphate dehydrogenase
