## Supplementary figures and images for "Experimental change in personality: Overexpression of GDNF in the rat striatum converts the low exploratory phenotype into highly explorative"

### Supplemental Figure 1

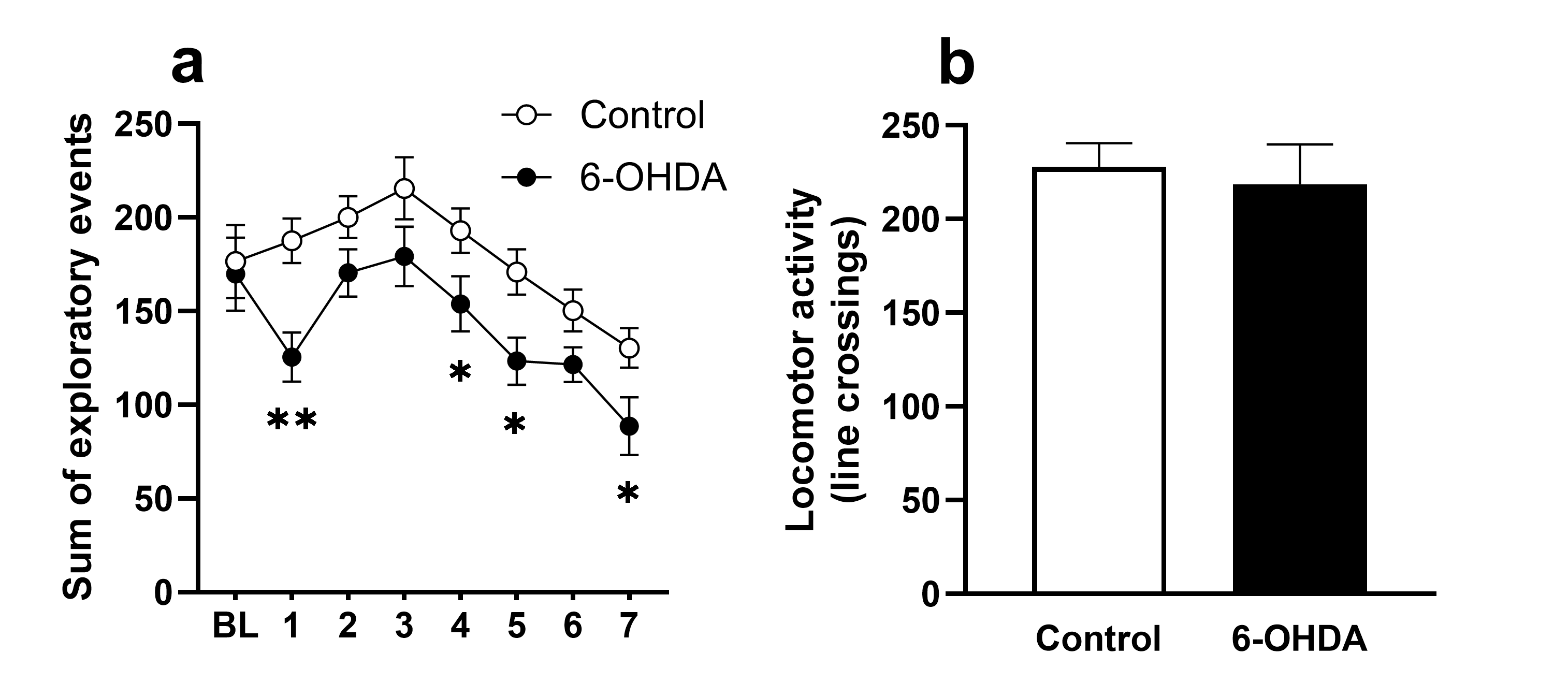

### Supplemental Figure 2

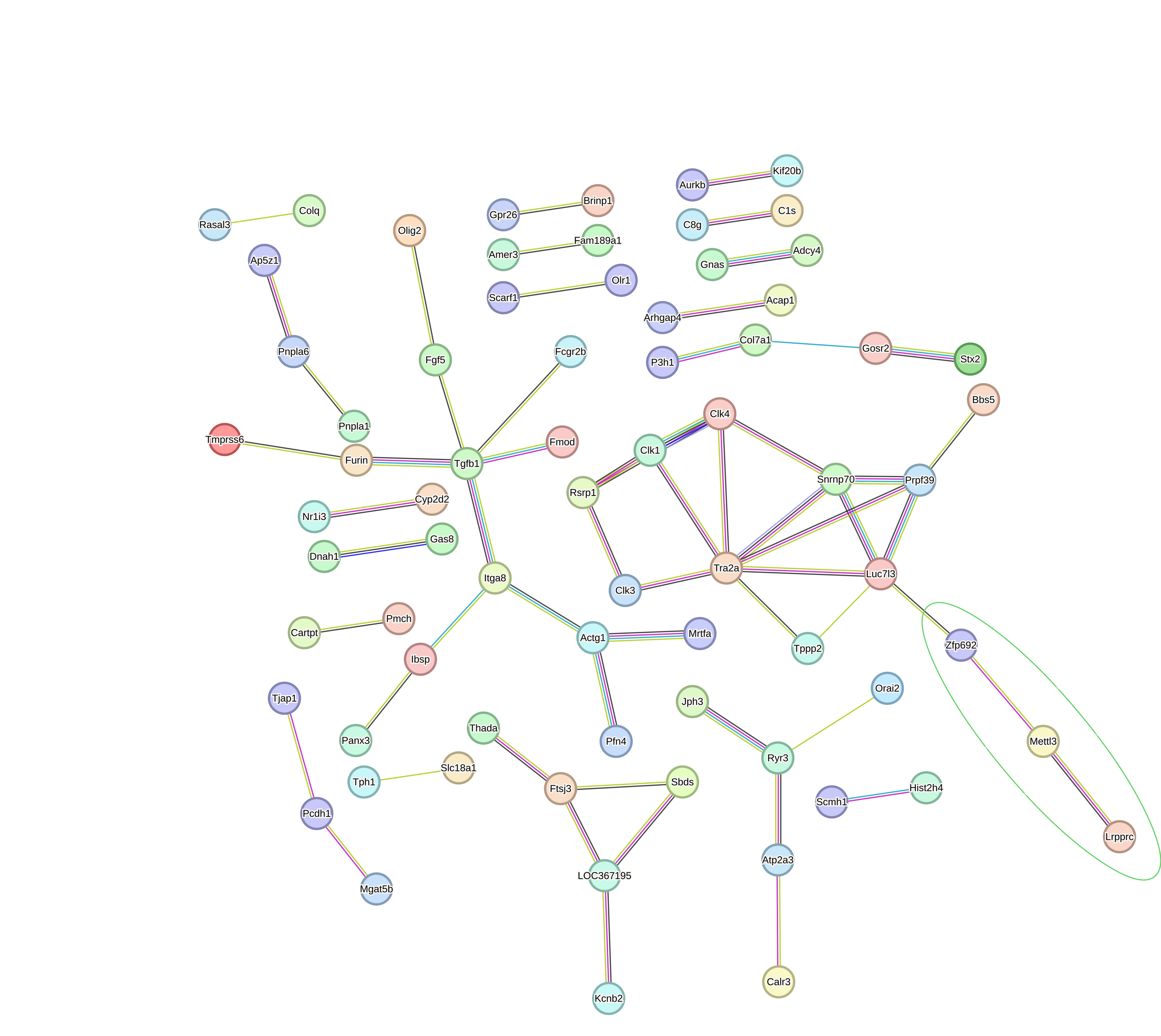
